# Osmotic pressure enables high yield assembly of giant vesicles in solutions of physiological ionic strengths

**DOI:** 10.1101/2022.10.08.511425

**Authors:** Alexis Cooper, Vaishnavi Girish, Anand Bala Subramaniam

## Abstract

Giant unilamellar vesicles (GUVs) are micrometer scale minimal cellular mimics that are useful for synthetic biology and drug delivery. Unlike assembly in low-salt solutions, assembly of GUVs in solutions with ionic concentrations of 100-150 mM Na/KCl (salty solutions) is challenging. Chemical compounds deposited on the substrate or in the lipid mixture could assist in the assembly of GUVs. Here, we investigate quantitatively the effects of temperature and chemical identity of six assisting polymeric and small molecule compounds on the molar yields of GUVs composed of three different lipid mixtures using high resolution confocal microscopy and large dataset image analysis. All the polymers moderately increased the yields of GUVs either at 22 or 37 degrees Celsius, whereas the small molecule compound was ineffective. Low gelling temperature agarose is the singular compound that consistently produces yields of GUVs of greater than 10 %. We propose a free energy model of budding to explain the effects of polymers in assisting the assembly of GUVs. The osmotic pressure exerted on the membranes by the dissolved polymer balances the increased adhesion between the membranes, thus reducing the free energy for bud formation. Data obtained by modulating the ionic strength and ion valency of the solution shows that the evolution of the yield of GUVs support our model’s prediction. In addition, polymer specific interactions with the substrate and the lipid mixture effects yields. The uncovered mechanistic insight provides a quantitative experimental and theoretical framework to guide future studies. Additionally, this work shows a facile means for obtaining GUVs in solutions of physiological ionic strengths.

## Introduction

Giant unilamellar vesicles, GUVs, vesicles with a single bimolecular wall of phospholipids with diameters ≥ 1 μm, mimic the minimal configuration of biological cells^1,2^. GUVs are useful for applications in bottom-up synthetic biology and biomedicine^3–12^. Conventionally it has been challenging to assemble GUVs in solutions with ionic concentrations ∼100 – 150 mM Na/KCl (salty solutions) thin-film hydration^1,13,14^. This limitation presents a longstanding challenge for the use of GUVs in applications since biomolecules such as proteins, nucleic acids, and polysaccharides require salty solutions to function^15–19^.

Several approaches have been proposed to improve the yields of GUVs in salty solutions. Soluble hexoses such as fructose^20^, macromolecular polymeric films composed of ultralow gelling temperature agarose^21^, polyvinyl alcohol (PVA)^22^, crosslinked polyacrylamide^23^, and crosslinked dextran(polyethylene glycol)^24^ have been used to assist the assembly of GUVs in salty solutions. The relative effectiveness of these various compounds compared to each other and the yields of GUVs that they produce however, is unknown. Importantly, the lack of quantitative data hinders a physicochemical understanding of the mechanism through which these compounds exert their effects on the assembly of GUVs.

Here, we investigate the molar yields of GUVs in salty solutions as a function of the temperature and chemical identity of the assisting compounds. We performed experiments using three lipid mixtures, the zwitterionic phospholipid diolyeoyl-*sn*-glycero-3-phosphocholine (DOPC), a lipid mixture that minimally mimics the composition of the endoplasmic-reticulum-Golgi intermediate compartment (ERGIC) membrane^25^, and a mixture that minimally mimics the composition of the exoplasmic leaflet of the mammalian cellular membrane (mammalian exoplasmic leaflet (MEL))^26^. We evaluated the small molecule fructose, the synthetic macromolecular polymer PVA, and four natural macromolecular polysaccharides, agaroses with varying gelling temperatures.

The yield of GUVs assembled in low-salt solutions without assisting compounds ranged from a low of 5 % for the MEL mixture to a high of 25 % for the ERGIC mixture. The yield of GUVs from the DOPC mixture was 17 %. In salty solutions, the yields of GUVs for all the lipid mixtures were very low, < 1 %. All the polymeric compounds were able to assist the assembly of GUVs in salty solutions either at an incubation temperature of 22 °C or 37 °C. Apart from low gelling temperature agarose, experiments with all the polymers as assisting compounds resulted in only moderate to low yields of GUVs in salty solutions, ranging from 2 % to 10 %. Fructose was ineffective as an assisting compound, producing very low yields, < 1 %, at all temperatures.

We develop a free energy model of budding to explain the effects of macromolecular polymers on the yield of GUVs. High concentrations of ions increase the intermembrane adhesion potential between lipid bilayers in lamellar stacks. Our model shows that the gradient in osmotic pressure imposed by the dissolving polymer can balance the increased adhesion potential of the membranes in salty solutions. We test the model by experimentally modulating the adhesion potential using buffers of varying ionic strengths and ion valencies. Our results support the prediction that the osmotic pressure of the dissolving polymer acts to oppose adhesion between membranes. The polymers’ contribution to the osmotic pressure results in a net reduction in the free energy for bud formation.

Additionally, we find that interactions of the polymers with the substrate and with the lipid mixture effects yields. Dewetting of the polymer from the substrate results in the formation of polymer-lipid ‘pseudobuds’ that morphologically resemble GUV buds but do not produce free-floating vesicles. When the moderately anionic low gelling temperature agarose^27^ was used as the assisting compound, the yield of GUVs composed of the negatively charged ERGIC mixture decreased by 7 % and 10 % respectively. Conversely, when the highly anionic PVA^28^ was used as the assisting compound, the yield of GUVs composed of the ERGIC mixture increased by 5 % when compared to the DOPC and MEL mixtures. These additional polymer and lipid specific interactions are challenging to incorporate in a minimal model for budding. In aggregate, we find that low gelling temperature agarose is the singular compound that consistently produces yields of GUVs of ≥ 10 %. Partial dissolution of the polymer with minimal dewetting is essential for obtaining high yields of GUVs in salty solutions.

## Results and Discussion

### Properties of the compounds tested

Figure 1 shows the chemical structure and properties of the compounds that we tested. PVA is a water-soluble highly anionic synthetic polymer composed of vinyl monomer units^28^. The PVA used in this study has a molecular weight of 146-186 kDa and is ≥ 99 % hydrolyzed. Uncrosslinked PVA at a concentration of 5 wt % or less does not gel and remains a viscous liquid at room temperature^28^. Dehydrated PVA forms a partially soluble swollen polymer film when rehydrated below its glass transition temperature of ∼ 85 °C^28^.The solubility of this polymer in water is 6 % at 20 °C and 13 % at 40 °C^28^.

**Figure 1.**
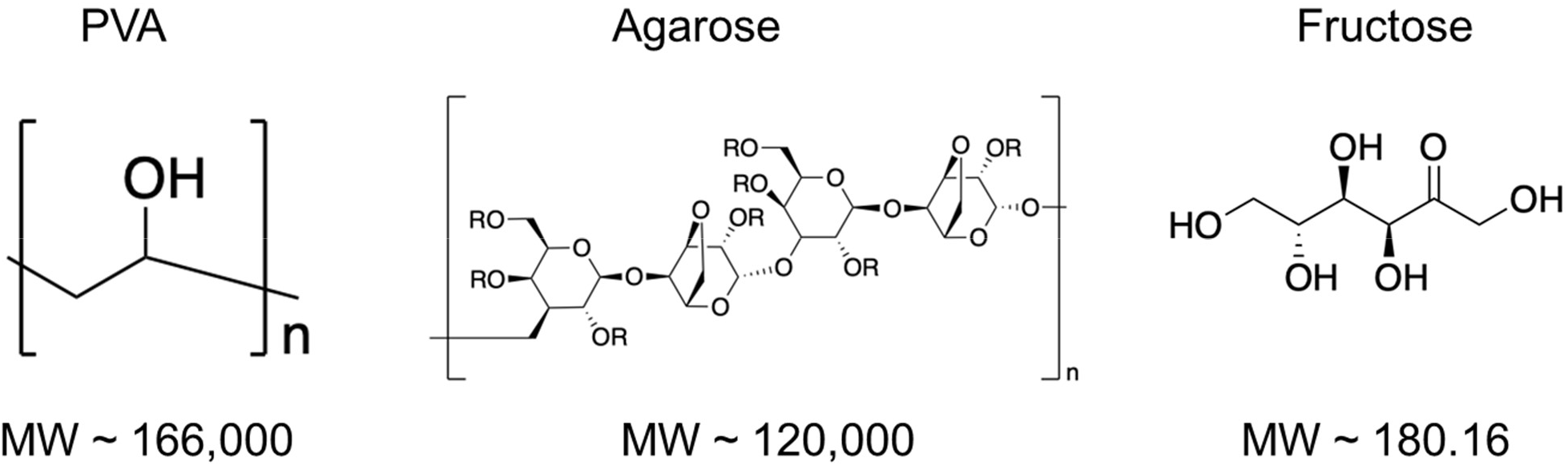
Structural formulas and molecular weight of the compounds. Structural formula of agarose^59,60^ R = H, CH_3_ or CH_2_CH_2_OH ^30^.

Agarose is a naturally derived polysaccharide composed of 1,3-linked β-D-galactopyranose and 1,4-linked 3,6-anhydro-α-L-galactopyranose with an average molecular weight of 120 kDa^29^. Solutions of agarose are conventionally prepared by dissolving 2 wt % or less of agarose powder at elevated temperatures^27^. When the solution is cooled to below the ‘gelling’ temperature, the agarose polymers transition from random coils into double helices^30^. The double helices hydrogen bond to form a percolated gel network^30^. Heating to above the ‘melting’ temperature dissolves the gel into a liquid polymeric solution consisting of random agarose coils^30^. Agarose exhibits thermal hysteresis. The melting temperature is significantly higher than the gelling temperature^31^. Synthetic hydroxyethylation can modify the gelling and melting temperature of agarose^27,30^. We refer to the agaroses by the manufacturer’s classification as ultralow gelling temperature agarose (ULGT), low gelling temperature agarose (LGT), medium gelling temperature agarose (MGT), and high gelling temperature agarose (HGT). We show the gelling temperatures of the agaroses in Supporting Information Table S1. We expect that at room temperature ULGT, LGT, and MGT agarose to demonstrate partial solubility, while HGT agarose is expected to be insoluble.

Fructose is a highly water-soluble small molecule monosaccharide with a molecular weight of 180.16 Da^32^. Fructose is thus ∼ 600× smaller than PVA and agarose.

### Poorly soluble HGT agarose and highly soluble fructose are ineffective at assisting the assembly of GUVs in salty solutions at 22 °C

We assembled GUVs composed of the zwitterionic lipid DOPC by hydrating the lipid-coated surfaces in a solution of phosphate buffered saline (PBS) + 100 mM sucrose at 22 °C (room temperature). This incubation temperature was above the gelling temperature of the ULGT agarose and below the gelling temperature of the LGT, MGT, and HGT agaroses. The sucrose was necessary to obtain a density gradient for sedimentation and is present in all the hydration buffers that we used. After 2 hours of incubation, we harvested the vesicle buds from the surfaces and compare the molar yields of the resultant GUVs. The molar yield measures the moles of lipid in the membranes of the population of harvested GUVs relative to the moles of lipids that were initially deposited on the substrate^33^. The molar yield is an objective measure that allows quantitative comparison of the effects of experimental variables on the yield of GUVs^33^. Similar to our previous work, which used no assisting compounds, we allow the GUVs to sediment for 3 hours prior to obtaining high resolution single plane open-pinhole tile scan images with a confocal microscope ^33^.

We confirmed that the GUVs assembled with the use of assisting compounds showed similar sedimentation behavior to those assembled without any assisting compounds over the 3-hour time frame (See Supporting Text, Supporting Figure S1). Any statistically significant differences in the measured molar yields are thus because of the assisting compounds and the experimentally controlled assembly conditions and not because of differences in sedimentation behavior or measurement techniques. To account for experimental variation, experiments for each condition was repeated three independent times and the data is reported as the mean of the three independent repeats. We perform balanced one-way analysis of variance (ANOVA) tests to analyze the statistical significance of differences among multiple means. If the ANOVA revealed at least one of the conditions had a significant effect on the molar yield, we performed Tukey’s Honestly Significant Difference (HSD) post hoc tests to determine the significance between pairs of conditions. For comparison of two pairs of means we performed Student’s t-tests. Tables of F- and p-values of the statistical tests are shown in Table S2 – S9 in the Supporting Information.

Figure 2 shows a stacked bar plot of the molar yields. We divide the yield data into small GUVs (diameters *d*, 1 μm ≤ *d* < 10 μm), large GUVs (10 μm ≤ *d* < 50 μm), and very large GUVs (*d* ≥ 50 μm). Error bars are the standard deviation from the mean. We show the histogram of the distribution of diameters in Supporting Information Figure S2 and representative images of the harvested vesicles in Supporting Information Figure S3.

**Figure 2.**
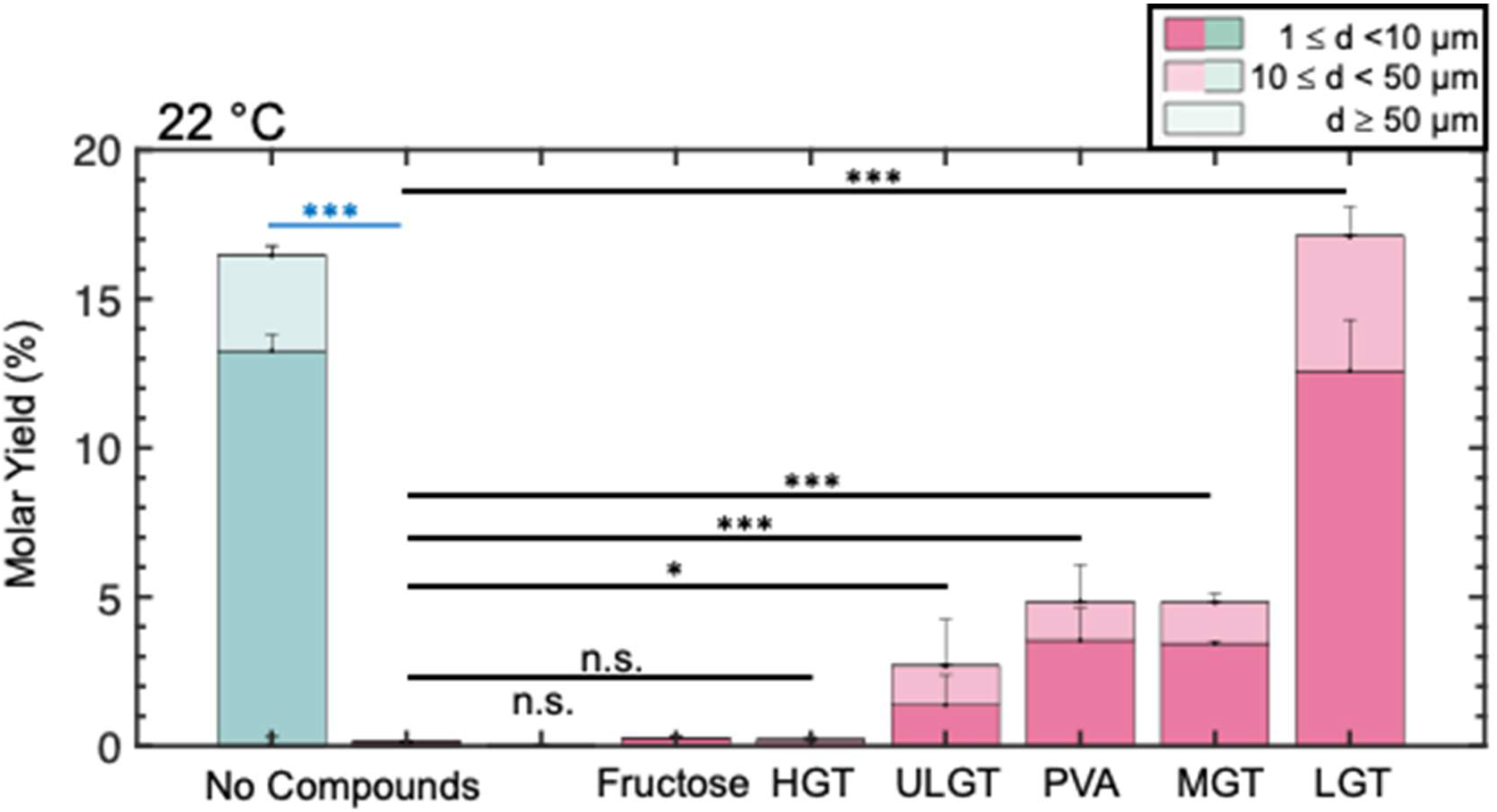
Stacked bar plots of the molar yields of GUVs in salty solutions at 22 °C. The blue bar is for samples hydrated in a low salt solution consisting of 100 mM of sucrose. The pink bars are molar yields for samples hydrated in PBS doped with 100 mM sucrose. The two leftmost bars show the molar yield without any assisting compounds. Each bar is split into 3 regions corresponding to the diameter ranges specified in the legend. Statistical significance was determined using a balanced one-way ANOVA and Tukey’s HSD post hoc tests (black) and Student’s t-test (blue). * = p < 0.05, ** = p < 0.01, *** = p < 0.001, n.s. = not significant.

To isolate the effects of the assisting compounds on assembly in salty solutions, we perform experiments on bare glass surfaces without any assisting compounds in PBS+100 mM sucrose and in a low-salt buffer consisting only of 100 mM sucrose. At room temperature, the yields of GUVs from bare glass in a hydrating solution consisting of 100 mM of sucrose is 16.5 ± 0.3%. This result is consistent with previous reports for gentle hydration on glass surfaces^33^. The yield of GUVs that we obtained from the lipids deposited on bare coverslip surfaces in the buffer containing PBS was ∼ 100 times lower, 0.2 ± 0.0 % (*p* = 6.73 × 10^−8^), than the yield in buffer consisting only of sucrose. We next assessed the effect of assisting compounds on the yields of GUVs in PBS. Use of fructose-doped lipid and HGT agarose as the assisting compounds resulted in a yield of 0.2 ± 0.0 % and 0.3 ± 0.0 % which was statistically indistinguishable from bare glass (both *p* ≥ 0.999). Use of ULGT agarose, MGT agarose, and PVA as the assisting compounds resulted in statistically significant increases in the yields of GUVs compared to assembly without any assisting compounds (all *p* < 0.05). However, the yields were statistically indistinguishable from each other at 2.7 ± 0.2 %, 4.8 ± 0.5 %, and 5.0 ± 1.0 % respectively (all *p* > 0.05). Use of LGT agarose as the assisting compound resulted in the highest yield of GUVs at 17 ± 1 %. This yield is more than three times higher than the yield of GUVs obtained when ULGT agarose and PVA are used as the assisting compounds. This result is notable since ULGT agarose and PVA are both used extensively in the literature^34–42^, whereas, as far as we know, we are the first to report the use of LGT agarose. Our results are general. Use of ULGT and LGT agaroses with different catalog numbers as the assisting compounds resulted in similar yields to those shown in Figure 2 (Supporting Information Figures S4 and S5).

### An increase in the incubation temperature to 37 °C increases the yields of GUVs for most of the compounds but decreases the yield for LGT agarose

HGT agarose is the least soluble of the polymeric compounds tested and fructose is the smallest and most soluble of the compounds tested. Both these compounds were ineffective at assisting the assembly of GUVs (Fig. 2). Since the size and apparent solubility of the compounds appears to have an effect on the yields of GUVs, we devised an experiment to test for the effect of polymeric solubility by assembling the GUVs at 37 °C. Naively, we expect that increasing the temperature should, 1) enhance the yield of GUVs by increasing the solubility of the polymers, 2) have no effect on the yield of GUVs when the small-molecule fructose is used as an assisting compound. Figure 3 shows the results of our experiments. We show representative images of the harvested GUVs in Supporting Information Figure S6 and the histogram of the distribution of diameters in Supporting Information Figure S7.

**Figure 3.**
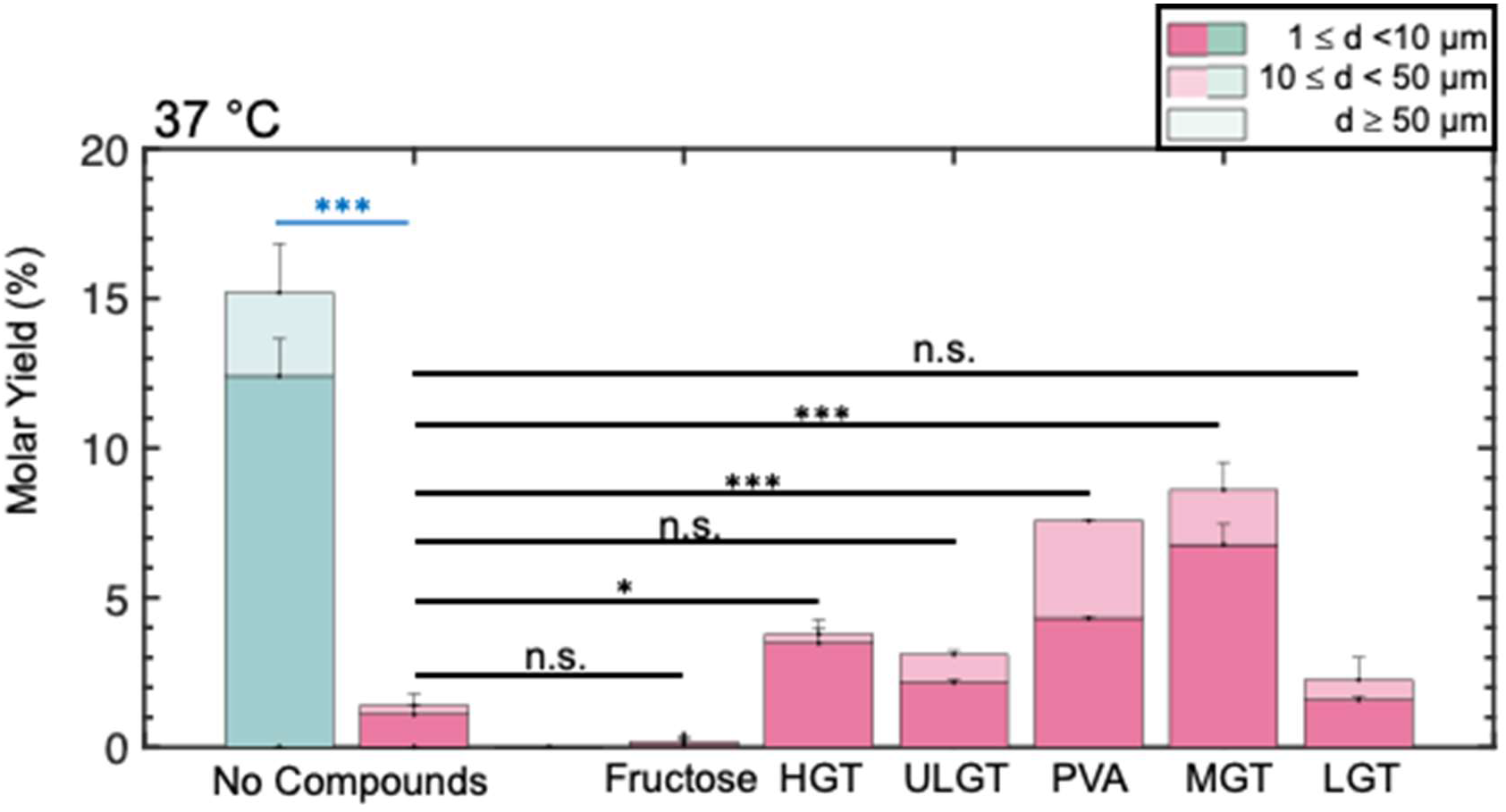
Stacked bar plots of the molar yields of GUVs in salty solutions at 37 °C. The blue bar is for samples hydrated in a low salt solution consisting of 100 mM of sucrose. The pink bars are molar yields for samples hydrated in PBS doped with 100 mM sucrose. The two leftmost bars show the molar yield without any assisting compounds. Each bar is split into 3 regions corresponding to the diameter ranges specified in the legend. Statistical significance was determined using a balanced one-way ANOVA and Tukey’s HSD post hoc tests (black) and Student’s t-test (blue). * = p < 0.05, ** = p < 0.01, *** = p < 0.001, n.s. = not significant.

An increase in temperature could in principle increase the yields of GUVs independent of any effects of the assisting compounds We thus performed experiments to determine the effect of the increase in temperature on the yield of GUVs without any assisting compounds. The yield of GUVs on bare glass in the low-salt sucrose buffer was 15 ± 1 %. This yield was statistically indistinguishable from assembly at room temperature. Conversely, the increased temperature resulted in a modest but statistically significant increase in the yields of GUVs on bare glass in PBS to 1.4 ± 0.4 % (*p*= 0.00480).

Use of PVA, MGT agarose, and HGT agarose as the assisting compounds resulted in statistically significant increases in the yield of GUVs compared to assembly performed at room temperature, 7.6 ± 0.8 % (*p* = 0.0314), 9.0 ± 0.3 % (*p =* 0.0224), and 3.8 ± 0.5 % (*p =* 2.20 × 10^−4^) respectively. The yield of GUVs from the fructose-doped lipid was unaffected by the change in temperature. These four results are consistent with our naïve expectations of the effect of temperature on the yields of GUVs.

Use of ULGT agarose showed no change in yields at 3.1 ± 0.1 % while use of LGT agarose showed an almost 9-fold *decrease* in the yield of GUVs to 2.0 ± 0.1 % (*p* = 1.40 ×10^−5^). This dramatic drop in yield for LGT agarose resulted in the yields of GUVs from the samples prepared with ULGT agarose and LGT agarose as the assisting compounds to be statistically indistinguishable from simple gentle hydration in PBS at 37 °C. Note that 37 °C was above the gelling temperature of ULGT and LGT agarose. We conclude that assembly at temperatures exceeding the gelling temperature of the agarose results in dramatically lowered yields of GUVs. Further, the increase in the yield of GUVs with the increase in temperature when PVA,

MGT agarose, and HGT agarose are used as assisting compounds shows the importance of the solubility of the polymer for assisting in the assembly of GUVs in salty solutions. The decrease in the yield of GUVs when LGT agarose is used as the assisting compound and the unchanged yields when ULGT agarose is used as the assisting compounds, however, shows that additional factors play a role.

### Characterization of the hydrated lipid films reveals the formation of polymer-lipid pseudobuds due to the dewetting of the polymer

To explore potential factors that could explain our results, we imaged the lipid-coated surfaces prior to harvesting using high resolution single plane confocal microscopy. On bare coverslips, we observe flat fluorescent surfaces with stepped differences in intensity and few spherical buds (Fig. 4a). These images are reminiscent of supported lipid bilayers on glass surfaces^43^. We interpret the stepped difference in fluorescence intensity as overlapping bilayers in a stack. Spherical buds can be seen in typical fields of views on the surfaces incubated at 37 °C compared to surfaces incubated at room temperature (Fig. 4b, white arrow). Images of the fructose-doped lipid did not show any regions that appeared to be lipid bilayers or vesicle buds (Fig. 4c). Instead, the surface was characterized by irregular punctate structures. We interpret these structures as being lipid aggregates. The lipid film appeared smooth on HGT agarose at room temperature (Fig. 4d).

**Figure 4.**
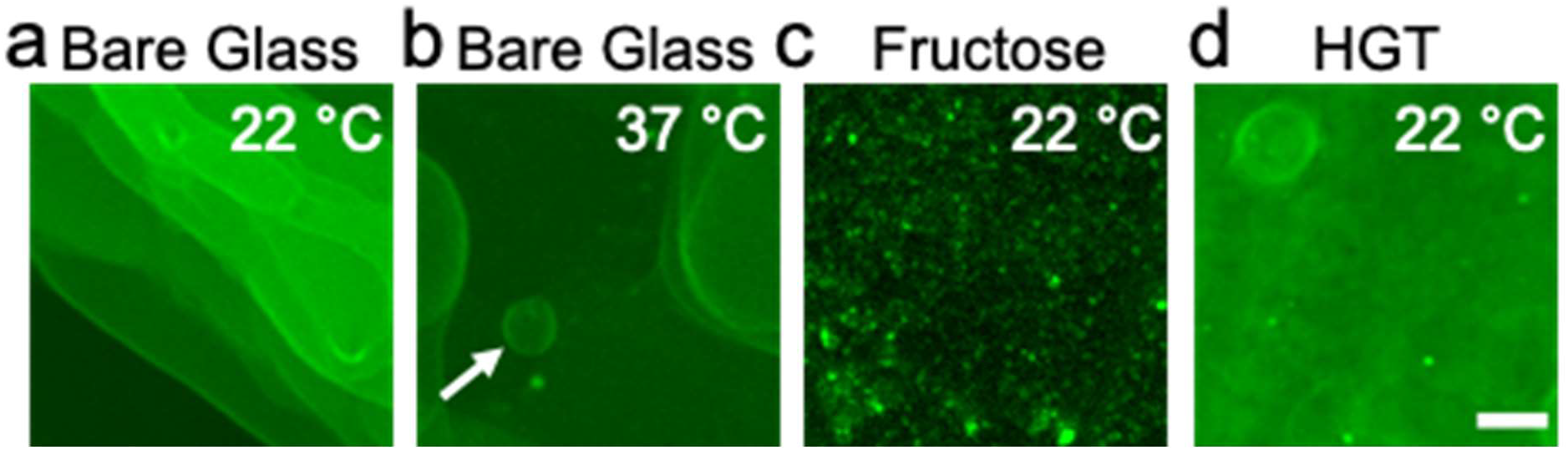
Surfaces that show little to no spherical buds. All samples hydrated in PBS doped with 100 mM sucrose. a) Bare glass, 22 °C, b) bare glass, 37 °C. White arrow points to a spherical bud. c) Fructose, 22 °C, d) HGT agarose, 22 °C. The scale bar is 15 μm.

We find a high density of spherical structures reminiscent of GUV buds on all the other polymer coated surfaces (Fig. 5). Similar to assembly without assisting compounds^33^, the GUV buds remain attached to the surface prior to harvesting (Supporting Information Figure S8). Puzzlingly, surfaces that appeared to have a high density of large spherical buds such as PVA-coated surfaces and ULGT agarose-coated surfaces at room temperature and ULGT, LGT, and MGT agarose coated surfaces at 37 °C had relatively low yields of free-floating GUVs. Furthermore, these surfaces often had buds of larger diameters and of higher surface densities compared to the high yielding LGT agarose-coated surfaces at room temperature.

**Figure 5.**
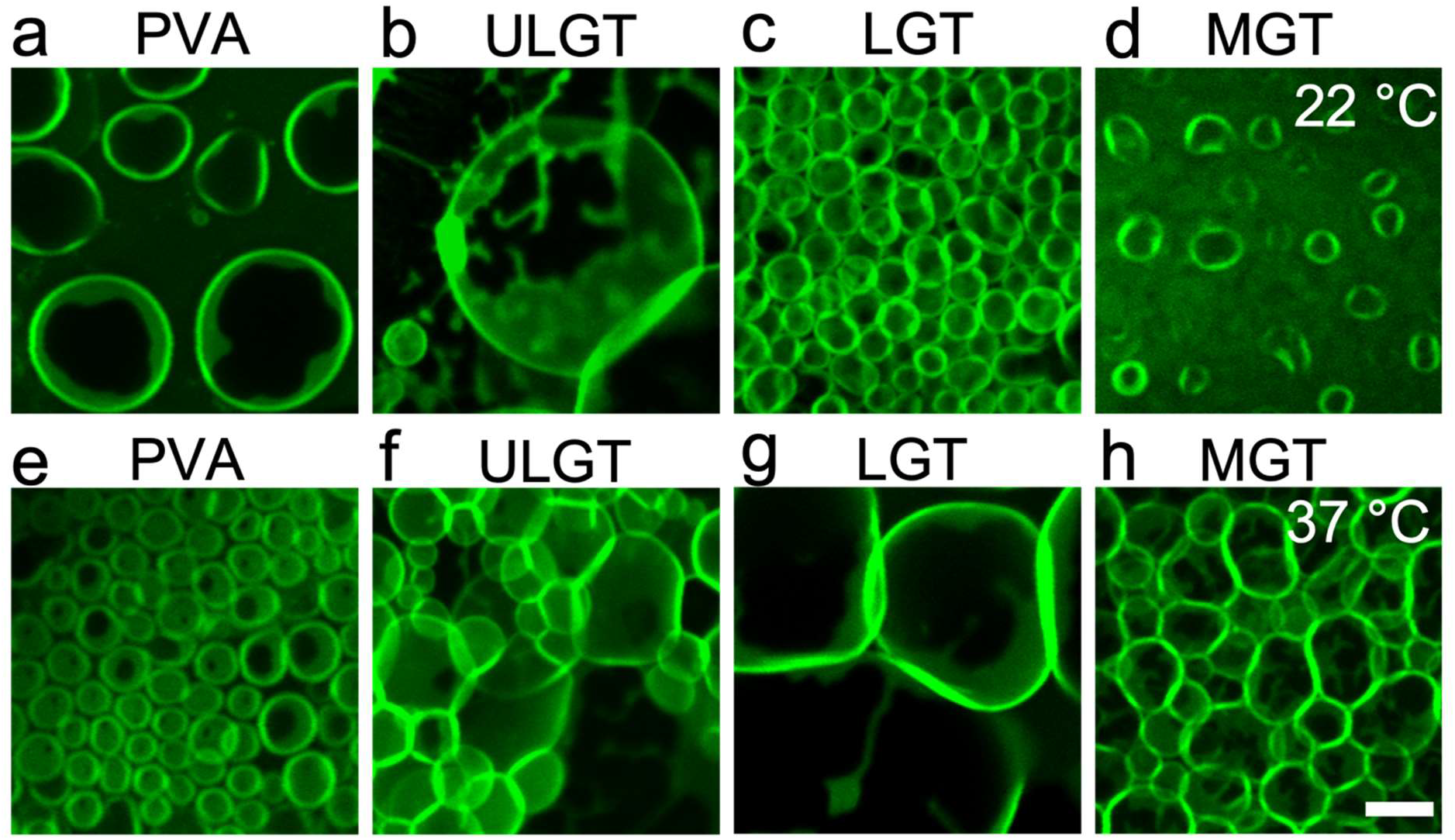
Surfaces that show circular buds. The top row shows images from samples hydrated at 22 °C. The bottom row shows images from samples hydrated at 37 °C. All samples hydrated in PBS doped with 100 mM of sucrose. The scale bar is 15 μm.

To understand further why surfaces with high numbers of apparent buds showed low yields of free-floating GUVs, we evaluated the structure of the buds by examining the distribution of fluorescence intensity. GUV buds imaged using confocal microscopy have a uniform fluorescence intensity^33^. Consistent with this previous observation, all the buds on surfaces that were incubated in low salt solutions have a uniform distribution of fluorescence intensities (Fig. 6a). On the polymer-coated surfaces in PBS, we find two types of structures. Buds with uniform fluorescence intensities reminiscent of GUV buds, (white arrows in Fig. 6b,c) and novel buds with dark regions of low fluorescence intensity (red arrows in Fig. 6b,c). These dark regions had shapes that could be spinodal-like (red arrows Fig. 6b) or circular (red arrows in Fig. 6c). Further, buds that showed these patterns often were larger and had higher membrane fluorescence intensities compared to the buds without dark regions (compare Fig. 6a with Fig. 6d).

**Figure 6.**
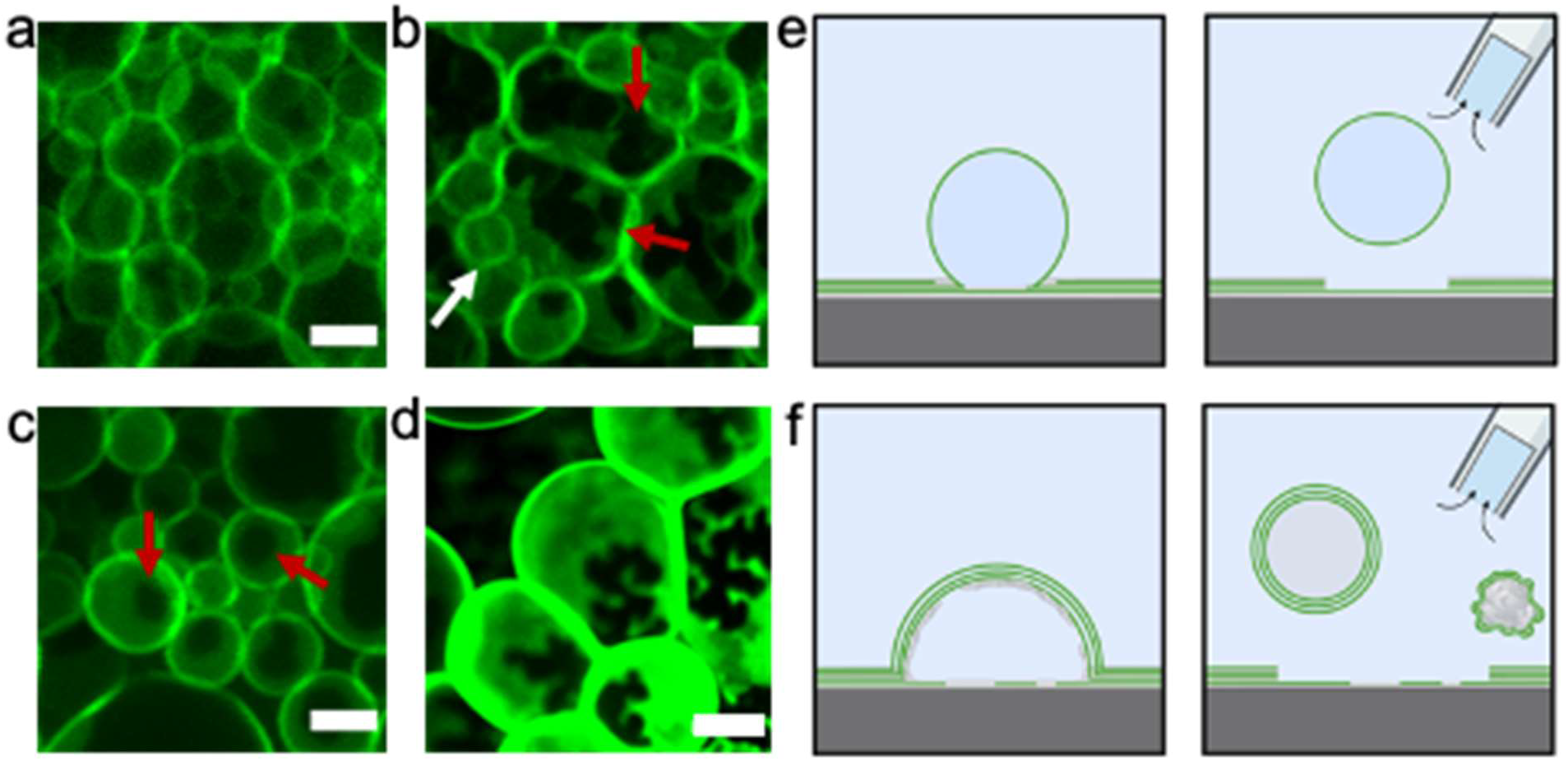
Polymer surfaces form pseudobuds as well as GUV buds. a) Spherical GUV buds with uniform fluorescence intensity as seen in on LGT agarose hydrated in 100 mM sucrose. b) Pseudobuds with spinodal dewetting pattern on LGT agarose (red arrows). GUV buds are highlighted by the white arrows. c) Pseudobuds with circular dewetting pattern on PVA (red arrows). White arrows point to regular GUV buds. d) Large pseudobuds with high fluorescence intensity on ULGT agarose. Schematic showing that e) GUV buds are expected to spontaneously close to form GUVs when harvested from the surface. f) Pseudobuds form non-GUV structures and lipid-coated polymer aggregates. b-d hydrated in PBS doped with 100 mM sucrose. The scale bars for a-c are 10 μm. The scale bar for d is 20 μm.

Based on these observations, we propose that on polymeric surfaces, two types of buds form (Fig. 6e,f). The first are regular GUV buds that self-close to form free-floating GUVs when scissioned from the surface during harvesting (Fig. 6e)^33^. The second are hybrid polymer-lipid ‘pseudobuds’ that nominally resemble GUV buds. We propose that the polymer-lipid pseudobuds arise when the lipid-coated polymer film swells and dewets from the surface (Fig. 6f). This interpretation explains the dark region at the base of the pseudobuds where the polymer has lifted with the lipid. Indeed, the spinodal and circular patterns at the base of the pseudobuds are reminiscent of spinodal and heterogenous nucleation patterns observed for polymer and polysaccharide films that dewet from a supporting substrate^44–47^. Localized dewetting of the polymer film and lifting of the stacks of lipid also explains the high membrane fluorescence intensity of the pseudobuds. The high fluorescence intensity suggests that pseudobuds are composed of multiple lipid bilayers. We posit that the layer of dewetted polymer that scaffolds the pseudobuds prevents closure of the multibilayer lipid membrane to form vesicles when scissioned from the surface during harvesting (Fig. 6f). Furthermore, even if the buds self-close, the multiple bilayers and the high amount of encapsulated polymer make these objects not GUVs.

We test for the formation of polymer-lipid pseudobuds using the lipid 1,2-dioleoyl-*sn*-glycero-3-phosphoethanolamine DOPE. DOPE cannot assemble into vesicles because it forms hexagonal phases instead of lamellar phases^48^. We thus expect no GUV buds to form on the surface. We use PVA and ULGT agarose, two polymers that likely cause the formation of high numbers of pseudobuds. Our images show clear structures resembling pseudobuds, and no structures resembling GUV buds (Fig. 7). On harvesting, we obtained no measurable yield of GUVs. We conclude that films of lipids and polymers can form pseudobuds that nominally resemble GUV buds. These pseudobuds are not productive for forming GUVs.

**Figure 7.**
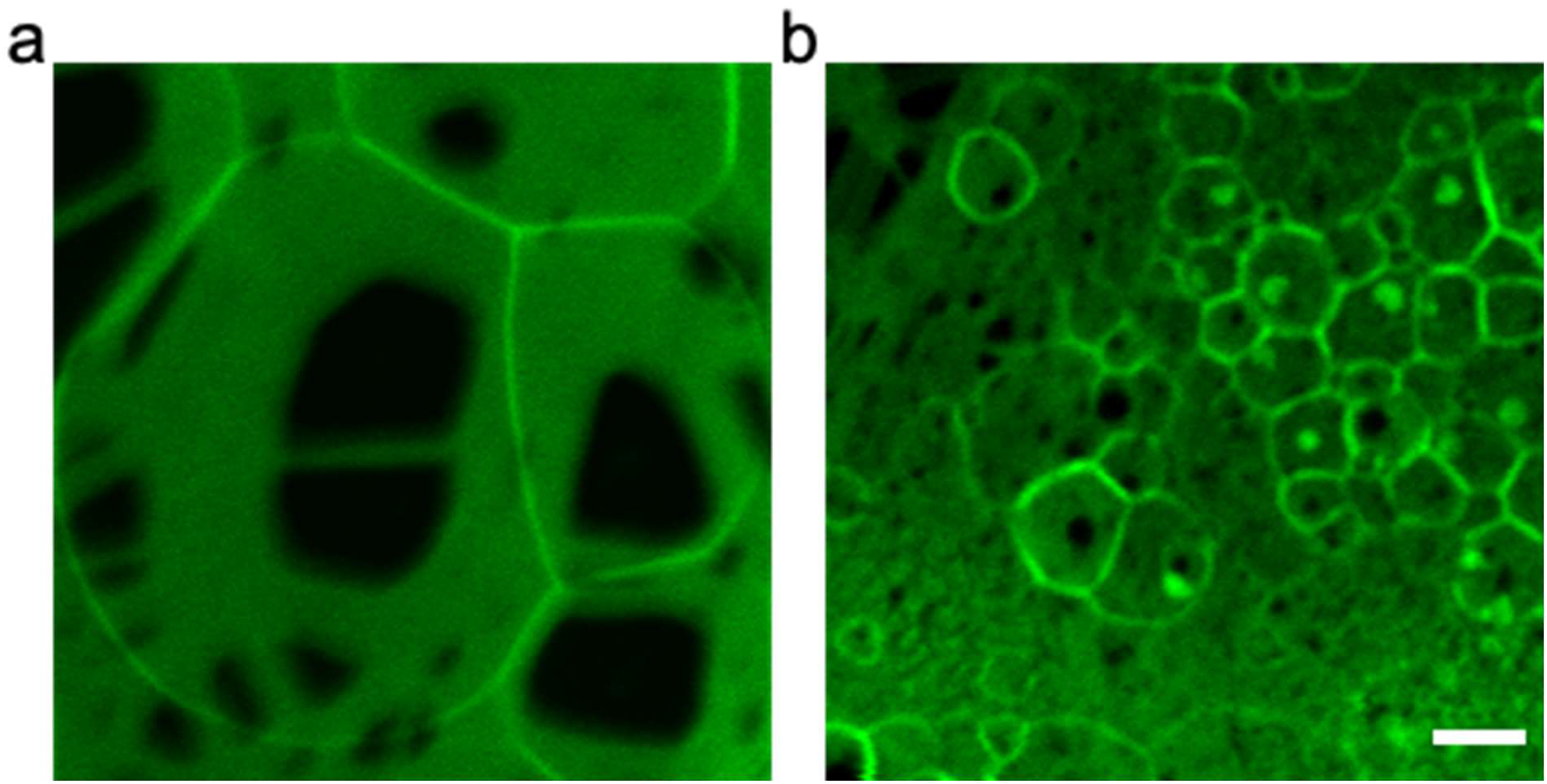
DOPE forms pseudobuds on ULGT agarose and PVA. a) Pseudobuds formed from DOPE on ULGT agarose showing regions of dewetting with low fluorescence intensity. b) Pseudobuds formed from DOPE on PVA showing dewetting patterns. DOPE does not form lamellar structures, so these buds are not GUV buds. Samples were hydrated in PBS doped with 100 mM of sucrose. Scale bar is 10 μm.

Analysis of our images showed that the number of pseudobuds were low for the LGT agarose-coated surface at room temperature, explaining the high yield of free-floating GUVs. At 37 °C which is above the gelling temperature of LGT agarose, the number of pseudobuds increases at the expense of GUV buds. The increased formation of pseudobuds due to dewetting explains the low yield of free-floating GUVs. All the other polymer surfaces have a high number of pseudobuds. Thus, although the surfaces of polymers can appear to be covered with a large number of spherical buds (Fig. 5), most of the buds cannot be harvested to form GUVs. Our discovery of pseudobuds explains previous observations of low yields of free-floating GUVs despite the apparent high numbers of spherical buds on the surfaces^13,22^. We surmise that dewetting of the polymer acts antagonistically to polymer solubility and reduces the yields of GUVs.

### Membrane composition modifies the yields of GUVs obtained from LGT agarose and PVA

We studied the effect of three assisting compounds, PVA, LGT agarose, and HGT agarose on the yields of GUVs obtained using an ERGIC mimicking mixture and a MEL mimicking mixture. The ERGIC mixture is negatively charged since it contains a high mol fraction of phosphatidylserine and phosphatidylinositol. Figure 8 shows the results of our experiments. We show the histogram of the distribution of diameters in Supporting Information Figures S9 and S10.

**Figure 8.**
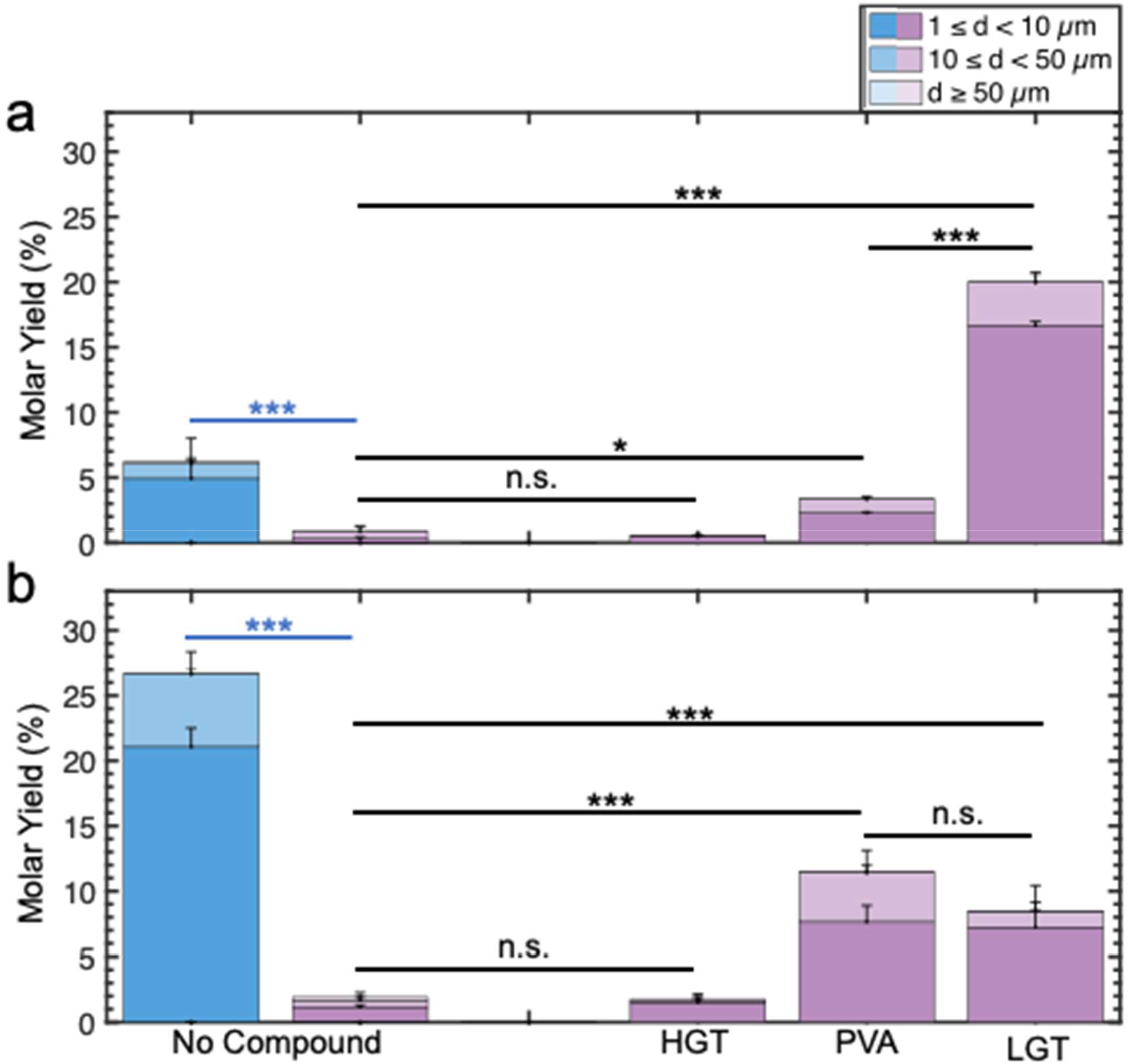
Stacked bar plots of the molar yields of GUV assembly in salty solutions. The blue bar is for samples hydrated in a low salt solution consisting of 100 mM of sucrose. The purple bars are molar yields for samples hydrated in PBS doped with 100 mM sucrose. a) MEL mixture. The two leftmost bars show the molar yield without any assisting compounds. b) ERGIC mixture. The two leftmost bars show the molar yield without any assisting compounds. Each bar is split into 3 regions corresponding to the diameter ranges specified in the legend. Statistical significance was determined using a balanced one-way ANOVA and Tukey’s HSD post hoc tests (black) and Student’s t-test (blue). * = p < 0.05, ** = p < 0.01, *** = p < 0.001, n.s.= not significant.

Similar to our approach with DOPC, we first measured the yield of GUVs obtained in low-salt and salty solutions without any assisting compounds. In low-salt solutions, we obtained a significantly lower yield of GUVs composed of the MEL mixture compared to the DOPC mixture, 6.2 ± 1.8 % (p = 6.07 × 10^−4^), and a significantly higher yield of GUVs composed of the ERGIC mixture compared to the DOPC mixture, 27 ± 1.7 % (p = 4.98 × 10^−4^). These results show that in low-salt solutions, the yield of GUVs depends on the composition of the membrane. Both the MEL mixture and ERGIC mixture had very low yields of GUVs, 0.8 ± 0.3 % and 1.9 ± 0.4 % respectively, in salty solutions. We conclude that assembly in salty solutions without any assisting compounds results in universally low yields of GUVs, < 2%.

In salty solutions, use of LGT agarose as the assisting compound resulted in an increase in the yield of GUVs from the MEL mixture to 20 ± 0.8 %, which was significantly higher than the yield obtained without the use of assisting compounds (*p* = 1.46 × 10^−13^). Use of PVA as the assisting compound resulted in a more modest yet statistically significant increase in molar yield of GUVs composed of the MEL mixture to 3.4 ± 0.1 % (*p* = 4.28× 10^−4^). Similar to our results with DOPC, use of HGT agarose as the assisting compound did not result in an increase in the yield of GUVs composed of the MEL mixture compared to bare glass, 0.9 ± 0.3 % (*p* = 0.738).

Use of LGT agarose and PVA as the assisting compounds resulted in statistically significant increases in the yield of GUVs composed of the ERGIC mixture to 8.5 ± 1.9 % and 11 ± 1.7 %, respectively. The difference in the mean yields of GUVs obtained between the two polymers was not statistically significant. Interestingly, the effect of the membrane composition and the chemical identity of the polymer was significant. When LGT agarose was used as the assisting compound, the resulting yields of GUVs composed of the ERGIC mixture was lower by 8.5 % (p = 5.46 × 10^−4^) and 11.5 % (p = 1.08 × 10^−4^) when compared to the zwitterionic DOPC and MEL mixtures respectively. Conversely, when PVA was used as the assisting compound, the resulting yields of GUVs composed of the ERGIC mixture increased by 6.2 % (p = 1.33 × 10^−3^) and 7.6 % (p = 4.43 × 10^−4^) when compared to the zwitterionic DOPC and MEL mixtures respectively. Similar to the DOPC and MEL mixtures, use of HGT agarose as the assisting compound, had no significant effect on the yield of GUVs composed of the ERGIC mixture, 1.7 ± 0.5 % (*p* = 0.998).

We surmise that in salty solutions, the poorly soluble HGT agarose was ineffective at increasing yields for all lipid mixtures, while the soluble LGT agarose and PVA increased yields for all lipid mixtures. Additionally, the membrane composition, likely the anionic nature of the ERGIC membrane, causes polymer specific changes in the yields of GUVs.

### The osmotic pressure exerted by dissolving polymers promotes the assembly of GUVs

Currently, there is no consensus on how assisting compounds promote the assembly of GUVs^13,20,22,49^. Upon hydration, lipids assemble into multibilayer stacks that conform to the geometry of the supporting solid substrate^50^. We had previously shown that in the absence of assisting compounds, gentle hydration of lipid films on surfaces composed of nanoscale cylindrical fibers (using nanocellulose paper in the PAPYRUS method (Paper-Abbetted amPhiphile hYdRation in aqUeous Solutions)) resulted in twice the yields of GUVs compared to flat surfaces^33^. We explained this result by showing that the free energy change for the formation of spherical buds from membranes templated on cylindrical fibers was lower than the free energy change from membranes templated on flat surfaces. In conditions where the energy to perform work is fixed, processes with low positive changes in free energy or high negative changes in free energy result in high yields of GUVs^33^.

Our results here show that on flat substrates, polymers that have partial solubility such as ULGT agarose, LGT agarose, MGT agarose, and PVA at 22 °C and HGT agarose at 37 °C can increase the yields of GUVs in salty solutions relative to bare glass. Dewetting of the soluble polymers from the surface on the other hand, favors the formation of pseudobuds and reduces the yields of GUVs. Further, when compared to the zwitterionic DOPC and MEL mixtures, the yield of GUVs from the anionic ERGIC mixture is enhanced when the highly anionic PVA is used as an assisting compound and is decreased when LGT agarose is used as an assisting compound. Clearly, interactions between the assisting polymers with the solid glass support and the membrane can disfavor or enhance the formation of GUV buds. Consistently, we find that in conditions where the polymer has low solubility such as when HGT agarose is used as the assisting compound at room temperature, the yields of GUVs remain unchanged compared to bare glass (Figure 2, Figure 8). To understand the mechanistic importance of polymer dissolution on the formation of buds we examine Equation 1.

Equation 1 shows the change in free energy for budding on flat surfaces, Δ*E*, retaining the pressure-volume term and dropping the edge energy term (See Supporting Information Text for further details).

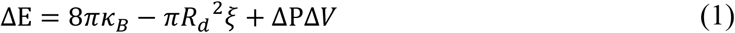

In this equation *κ_B_* is the bending rigidity of the membrane, *R_d_* is the radius of the flat lipid disk that forms the spherical GUV bud, *λ* is the edge energy, *ξ* is the adhesion potential between the membranes in a stack, Δ*P* is the difference in osmotic pressure, and Δ*V* is the difference between the volume of the spherical bud and the interlamellar volume enclosed by the disk in the stack. *ξ* is negative for attractive interactions. In the absence of an osmotic pressure, that is Δ*P* = 0, the free energy change for the formation of buds is always positive^33^. Energy due to hydrodynamic flows or temperature gradients^51^ provide work to form GUV buds.

Our data shows that the yield of GUVs in low-salt solutions depends on the composition of the lipid membrane. This result is consistent with the expected differences in membrane properties such as adhesion and bending rigidity due to differences in composition (Fig. 2, 8). Our data also shows that in salty solutions, the yields of GUVs from all three lipid compositions that we tested is very low (Fig. 2, 3, 8). This result suggests that the energy due to flows and temperature gradients, which was sufficient to produce high yields of GUVs in low-salt solutions, is insufficient to form GUVs in salty solutions. Dissolved ions increase adhesion between surfaces in aqueous solutions by screening electrostatic charges^48,52^. Using characteristic values of adhesion energy of *ξ* = 1 × 10 J m^2^ for DOPC membranes in low salt, *ξ* = 1 × 10 J m^2^ in salty solutions, *κ_B_* = 8.5 × 10^2^ J, and *R_d_* = 1.0 μm, in low-salt solutions Δ*E* = 1284 k T and in salty solutions Δ*E* = 76958 k T. The increase in the magnitude of the energy to form buds due to electrostatic screening is expected to decrease the number of buds formed. This result is consistent with our observed dramatic decrease in the yield of GUVs obtained through gentle hydration without assisting compounds in PBS compared to low-salt solutions (the leftmost bars in Fig. 2, 3, 8).

For polymer-coated surfaces, dissolution of the polymer can create a difference in osmotic pressure in the interlamellar space of the bilayer stacks. The osmotic pressure acts in the opposite direction to the adhesion potential since there is a high concentration of polymers on the surface and no polymers in the bulk solution. Thus Δ*P* is non-zero and negative. The negative third term in the right-hand side of Equation 1 allows the possibility for no change or even a net decrease in free energy that can balance an increase in the magnitude of the adhesion potential. Our estimates show that the concentration of polymer in the interlamellar space with a distance of 4 nm in a stack consisting of 5 bilayers is approximately 2.6 M. Approximately 0.009 % of the polymer molecules must dissolve in the interlamellar space, 0.25 mM, for the contribution of the osmotic pressure to result in similar budding energies between the polymer-free low-salt solutions and the polymer-assisted salty solutions (See Supporting Text for further discussion).

Since it is challenging to measure the dissolution of small amounts of polymer, we devise experiments to probe for the effects of osmotic pressure by modulating the magnitude of the adhesion potential of the membranes relative to PBS. The adhesion potential between membranes is lowest in solutions of low ionic strength. High concentrations of monovalent ions reduce the double layer screening length^48,52^, while mM concentrations of divalent cations can neutralize surface charges or function as ionic bridges^53,54^. These conditions promote adhesion between membranes^52,55,56^ (See Supporting Information Text, Table S10, Table S11 for further discussion and calculations of the double layer screening length). We reduced the adhesion potential relative to PBS by using 100 mM sucrose with no added salts and increased the adhesion potential by using 600 mM NaCl, a solution with a monovalent salt concentration that is more than four times that of PBS. We also used buffers with millimolar amounts of the divalent cations Mg^2+^ and Ca^2+^, PBS + 5 mM MgCl_2_ and 150 mM KCl + 5 mM CaCl_2_. The combinations of salts served to test for generality.

Figure 9 shows the molar yields of GUVs obtained when LGT agarose was used as the assisting compound. We show the histogram of the distribution of diameters in Supporting Information Figure S11. The yield of GUVs was affected significantly by the composition of the hydrating buffer. In solutions devoid of salt, the yield of GUVs doubled to 40 ± 3% compared to assembly in PBS (*p* =1.43 × 10^−7^). In 600 mM NaCl, PBS + 5 mM MgCl_2_, and 150 mM KCl + 5 mM CaCl_2_, the yield was approximately halved to 6.0 ± 1.0 % (*p* = 8.14 × 10^−5^), 9.0 ± 1.0 % (*p* = 0.00124), and 9.0 ± 1.0% (*p* = 0.00137) respectively. Clearly, buffers that decrease the adhesion potential result in an increase in the yield while buffers that increase the adhesion potential result in a decrease in the yield. These results are consistent with the osmotic pressure of dissolving polymers as an important mechanism behind the formation of GUVs on polymer-coated surfaces. Since dissolution of the polymer appears to be key for the formation of GUVs in salty solutions, extrapolating from our results for HGT agarose, cross-linking polymers to minimize dissolution will likely result in low yields of GUVs in salty solutions.

**Figure 9.**
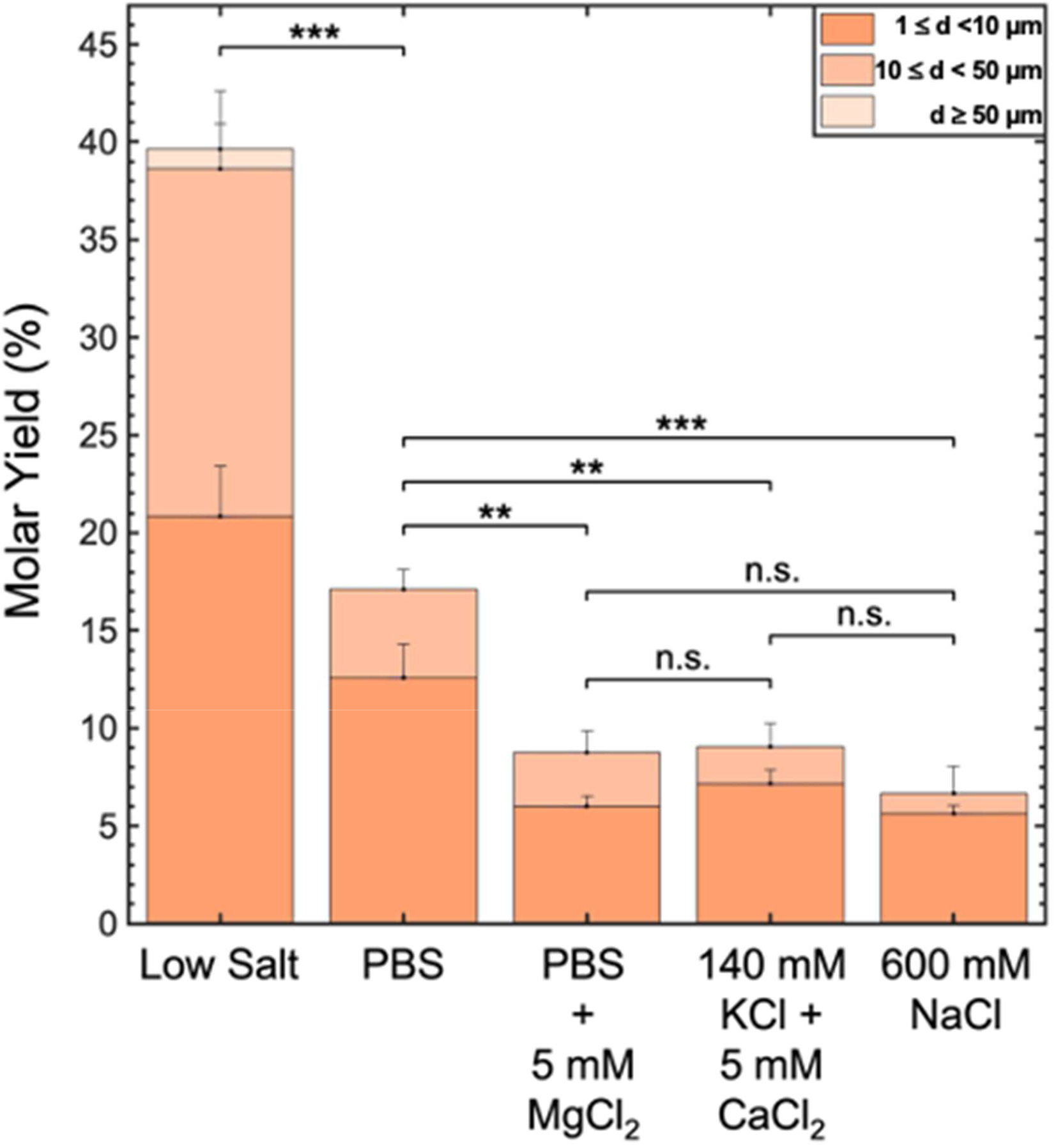
The yield of GUVs depends on the concentration and valency of ions. Stacked bar plot showing yields of GUVs assembled on LGT agarose hydrated in solutions containing low salt, PBS, PBS + 5 mM MgCl_2_, 140 mM KCl + 5 mM CaCl_2_, and 600 mM NaCl. The PBS data is reproduced from Figure 2. Each bar is split into 3 regions corresponding to the diameter ranges specified in the legend. Each bar is the average of 3 samples. Statistical significance was determined using a one-way ANOVA and Tukey’s HSD posthoc tests. * = p < 0.05, ** = p < 0.01, *** = p < 0.001, n.s. = not significant. All the buffers containing salts were also doped with 100 mM of sucrose.

Our model shows that the dissolution of the polymer is sufficient in principle to cause the formation of GUV sized buds in salty solutions. The concentration of polymer in the interlamellar volume is an important parameter. In all samples, the lipid that is dissolved in an organic solvent is deposited onto the dry polymer films. Rearrangement of the polymer and lipid in the transient milieu of the evaporating organic solvent and the subsequent hydration of the dry polymer/lipid film in the aqueous solvent to form lipid stacks interspersed with polymers likely determines the efficiency of the osmotic pressure mechanism. It is reasonable that the chemical composition of the polymer such as the presence of hydrophobic groups or charged groups effects the amount of polymer that incorporates in the interlamellar space of the lipid stack. Small molecule sugars are ineffective at increasing yields compared to simple gentle hydration on bare glass. The lack of effect of small sugars on yields that we find here are consistent with data from previous reports^21^ (Supporting Information Text). We suggest that the high solubility of sugars and their small size allows them to escape through defects in the bilayer stacks making sugars unable to exert an osmotic pressure against the membrane.

## Conclusion

Quantitative experiments reveal that the formation of GUVs from films of lipids in salty solutions depends significantly on the chemistry of the compounds, the assembly temperature, and the composition of the lipid membrane. Use of LGT agarose at room temperature as an assisting compound consistently resulted in the highest yields of free floating GUVs in salty solutions for all the lipid mixtures tested. Use of other polymers as assisting compounds resulted in moderate to low yields of GUVs. Experiments with solutions of varying ionic strengths shows that the difference in osmotic pressure due to dissolving polymers promotes the formation of GUVs. Although partial dissolution of the polymer is essential for increasing yields, specific interactions of the polymer with the substrate and the lipids can influence the yield. These results demonstrate the importance of measuring quantitative yields when novel assisting compounds, lipid mixtures, or temperatures are used in the assembly of GUVs. Looking forward, we propose that our quantitative experimental framework and our minimal free energy model provides a mechanistic guide for rational studies for discovering novel polymers that can further improve yields of giant vesicles in salty solutions, for example from amphiphilic block copolymers^57,58^.

## Methods Materials

We purchased glass coverslips (Corning, 22 mm × 22 mm) and premium plain glass microscope slides (75 mm × 25 mm) from Thermo Fisher Scientific (Waltham, MA).

### Chemicals

We purchased sucrose (BioXtra grade, purity ≥ 99.5%), glucose (BioXtra grade, purity ≥ 99.5%), potassium chloride (molecular biology grade, ≥ 99.0%), magnesium chloride (purity ≥ 98%), casein from bovine milk (BioReagent grade), agarose type IX-A: ultra-low gelling temperature (Catalog number: A2576, molecular biology grade), agarose: low gelling point (Catalog number: A9414, molecular biology grade), agarose type II-A medium EEO (Catalog number: A9918), agarose type VI-A: high gelling temperature (Catalog number: A7174), ultra-low gelling temperature agarose (Catalog number: A5030), low gelling temperature agarose (Catalog number: A0701) and poly(vinyl alcohol) (M_W_ 146,000 – 186,000 99+% hydrolyzed) (PVA) from Sigma-Aldrich (St. Louis, MO). We purchased chloroform (ACS grade, purity ≥ 99.8%, with 0.75% ethanol as preservative), Invitrogen 10× PBS Buffer (pH 7.4, 0.2 μm filtered, 1.37 M sodium chloride, 0.027 M potassium chloride, 0.080 sodium phosphate dibasic, 0.020 M potassium phosphate monobasic), sodium chloride (BioXtra grade, purity ≥99.5%) and D-(-)-Fructose (HPLC grade, purity ≥99%) from Thermo Fisher Scientific (Waltham, MA). We obtained 18.2 MΩ ultrapure water from an ELGA Pure-lab Ultra water purification system (Woodridge, IL). We purchased 1,2-dioleoyl-*sn*-glycero-3-phosphocholine (18:1 (Δ9-cis) PC (DOPC)), 23-(dipyrrometheneboron difluoride)-24-norcholesterol (TopFluor-Chol), 1,2-dioleoyl-*sn*-glycero-3-phosphoethanolamine (DOPE), 1-palmitoyl-2-oleoyl-glycero-3-phosphocholine (POPC), cholesterol (ovine wool, >98%), 1-palmitoyl-2-oleoyl-*sn*-glycero-3-phosphoethanolamine (POPE), 1-palmitoyl-2-oleoyl-*sn*-glycero-3-phosphoinositol (ammonium salt) (POPI), 1-palmitoyl-2-oleoyl-*sn*-glycero-3-phospho-L-serine (sodium salt) (POPS) and 1,2-distearoyl-*sn*-glycero-3-phosphoethanolamine-N-[methoxy(polyethylene glycol)-2000](ammonium salt) (PEG2000-DSPE) from Avanti Polar Lipids, Inc. (Alabaster, AL).

### Lipid mixtures

Lipid mixtures were prepared as previously described with minor adaptations^33^. Briefly, we prepared working solutions of 96.5:3:0.5 mol% DOPC: PEG2000-DSPE:TopFluor-Chol and 96.5:3:0.5 DOPE: PEG2000-DSPE:TopFluor-Chol, 66.5:29:3:0.5 POPC:Chol:PEG2000-DSPE 0:TopFluor-Chol and 41.5:20:13:7:15:3:0.5 POPC:POPE:POPI:POPS:Chol: PEG2000-DSPE:TopFluor-Chol at a concentration of 1 mg/mL. All lipid solutions were stored in Teflon-capped glass vials, purged with argon, and stored in a -20°C freezer. Lipid solutions were remade weekly.

### Formation of polymer films on glass

Films of polymer on glass coverslips (Corning, 22 mm × 22 mm) were prepared by applying 300 μL of 1 wt % (w/w) polymer on a coverslip and evenly spreading the solution with the side of a pipette tip^58^. The coated glass coverslips were allowed to dehydrate on a hotplate for a minimum of 2 hours set at a temperature of 40 °C. At the end of the process, the coverslip appeared flat and clear.

### Deposition of lipids

To ensure standardized conditions that allows comparison between samples, we deposit lipid solutions as described previously^33^. Briefly, circular disks with a diameter of 9.5 mm were traced on the underside of the polymer-coated coverslip, or bare coverslips using a template made from a circle hole punch (EK Tools Circle Punch, 3/8 in.). We evenly deposited 10 μL of the lipid working solution onto the polymer-coated or bare side of the glass within the traced area using a glass syringe (Hamilton). All lipid coated substrates were placed into a laboratory vacuum desiccator for 1 hour to remove any traces of organic solvent before hydration.

### Procedure for assembly

Circular poly(dimethyl)siloxane (PDMS) gaskets (inner diameter × height = 12 × 1 mm) were affixed to bare coverslips or polymer coated coverslips to construct a barrier around the dry solvent-free lipid films. We added 150 μL of PBS doped with 100 mM sucrose into the gaskets. To minimize evaporation, we place the gaskets and a water-saturated Kimwipe in a sealed 150 mm diameter Petri dish. The films were allowed to hydrate for 2 hours on a laboratory bench at room temperature. For assembly at 37 °C, we preheated the buffers to 37 °C in a water bath. The films were allowed to hydrate for 2 hours on a hotplate set to 37°C. To prevent evaporation, we covered the gaskets with glass coverslips, and placed the gaskets and a water saturated Kimwipe in a sealed 150 mm diameter Petri dish.

### Fructose-doped lipid method

Following a previously published protocol^20^, we prepared 1 mM (0.785 mg/mL) DOPC:TopFluor-Chol at 99.5:0.5 mol % in chloroform and 20 mM fructose dissolved in neat methanol. We mixed 50 μL of 1 mM 99.5:0.5 mol % DOPC:TopFluor-Chol in chloroform with 25 μL of 20 mM fructose in methanol to create a 2:1 chloroform:methanol working solution. We applied 15 μL of the solution onto the coverslip and place the coverslip in a vacuum chamber for 1 hour. The dried lipid film was then hydrated in PBS doped with 100 mM sucrose.

### Procedure for harvesting the GUVs

Harvesting GUVs from the substrated was conducted as previously described ^33^. Briefly, the GUVs were harvested by pipetting 100 μL of the hydrating solution with a cut 1000 μL pipet tip on 6 different regions of the lipid-coated surface to cover the whole area. We aspirated all the GUV containing liquid on the seventh time and transferred the liquid into a 0.5 mL Eppendorf tube. The final sample volume was ∼150 μL. Aliquots were taken immediately for imaging.

### Confocal microscopy of harvested vesicles

Imaging of harvested vesicles was conducted as previously described^33^. Briefly, we constructed imaging chambers by placing PDMS gaskets with a square opening (width × length × height = 6 × 6 × 1 mm) on glass microscope slides. Before use, we passivated the chamber with a solution of 1 mg/mL casein in PBS to prevent the rupture of GUVs on the surface of bare glass. Chambers were thoroughly rinsed with ultrapure water after passivation. We filled the passivated chamber with 58 μL of a 100 mM solution of glucose in PBS and evenly distributed a 2 μL aliquot of harvested GUV suspension into the 100 mM glucose in PBS solution by repeatedly pipetting 2 μL of the mixed suspension in glucose. We allowed the GUVs to sediment for 3 hours in a sealed 150 mm Petri dish with a water saturated Kimwipe to prevent evaporation before imaging. We captured images using an upright confocal laser-scanning microscope (LSM 880, Axio Imager.Z2m, Zeiss, Germany), using a 488 nm argon laser and a 10× Plan-Apochromat objective with a numerical aperture of 0.45. We imaged using an automated tile scan routine (64 images [850.19 μm × 850.19 μm (3212 pixels × 3212 pixels)]) to capture the entire area of the chamber. The routine used an autofocus feature at each tile location. Out of focus tiles were imaged manually. The pinhole was set at 15.16 Airy units which gave a confocal slice thickness of 79.3 μm.

### Imaging the surface of hydrated lipid films

We captured images of the surfaces using an upright confocal laser-scanning microscope (LSM 700, Axio Imager.Z2m, Zeiss, Germany), a diode-488 nm laser and a 10× Plan-Apochromat objective with a numerical aperture of 0.45. The frame size was 2048 pixels × 2048 pixels. The pinhole was set to 1 Airy Unit which gave a confocal slice thickness of 5.9 μm. Images were selected to be representative of the whole surface.

### Image processing and analysis

We conducted image processing and analysis as previously described^33^. Briefly, we used a custom MATLAB (Mathworks Inc., Natick, MA) routine to analyze the confocal tile scan images. The routine segmented fluorescent objects from the background. To obtain the diameters and mean intensities of the objects, we used the native *regionprops* function. We used the coefficient of variance of the intensities to select GUVs from the detected fluorescent objects All images were inspected after automated segmentation and erroneously segmented objects were manually corrected.

### Statistical analysis

All statistical analysis was performed using MATLAB. We conducted 1-way balanced analysis of variance (ANOVA) in MATLAB to determine the statistical significance of the mean yields obtained for the different compounds. We conduct a posthoc Tukey’s Honestly Significant Difference (HSD) to determine statistical significance of the differences in the mean between pairs of surfaces. To compare statistical significance of the difference of temperature on the yields we conduct Student’s t-tests.

## Supporting information

SupportingInfo

## ASSOCIATED CONTENT

### Supporting Information

Supporting text describing mathematical details of the molar yield calculations and free-energy model. Figures showing: Histograms of GUV diameters from different substrates. Confocal images of harvested, free-floating GUVs and buds on the surface of the substrate. Tables showing: Compounds tested, p-values for statistical tests, concentrations of salts in buffers, and calculated Debye screening lengths. Figure S1-S11, Table S1-S11, Equation S1-S9. (PDF).

## AUTHOR INFORMATION

### Corresponding Author

*

### Author contributions

A.B.S conceived and directed the study. A.C. performed experiments and V.G helped. A.C analyzed the data. A.B.S. and A.C. interpreted the data. A.C. prepared the figures. A.C and V.G. wrote the first draft of the manuscript. A.B.S developed the model and wrote the final draft. All authors have given approval to the final version of the manuscript.

## Acknowledgements

This work was funded by the National Science Foundation through NSF CAREER DMR-1848573. The data in this work was collected, in part, with a confocal microscope acquired through the National Science Foundation MRI Award Number DMR-1625733. The schematic in Fig. 6 was made using Biorender.

## Conflict of Interest

The authors declare no conflict of interest.

## TOC Graphic

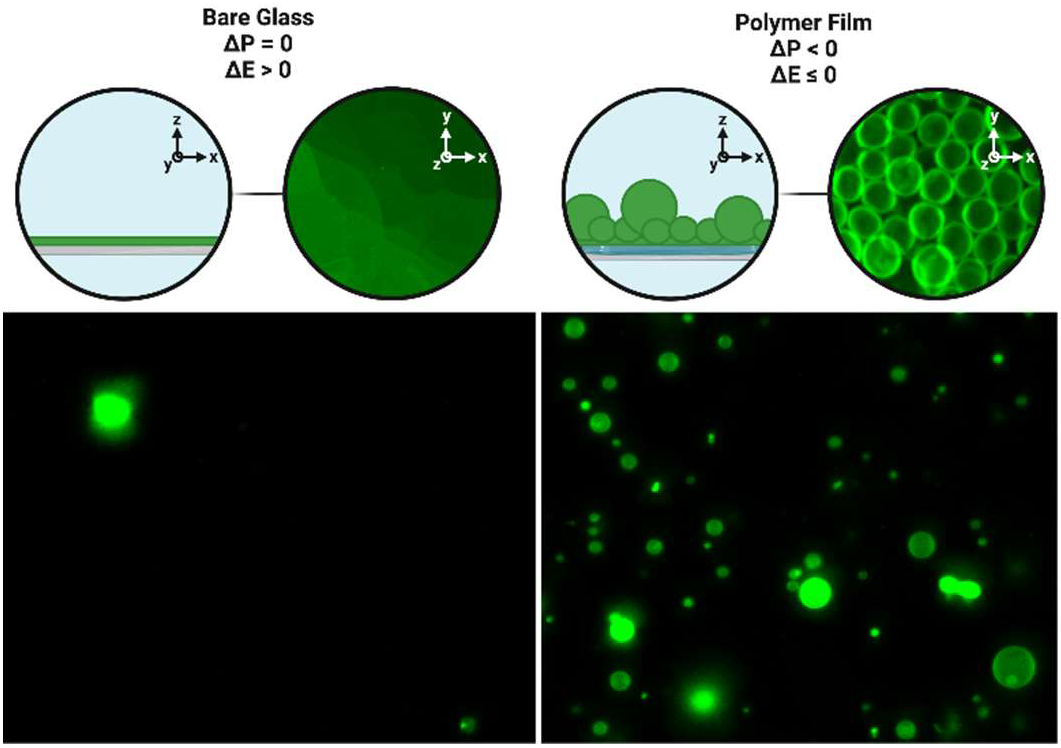

## Supporting Information

### Supporting Text

#### Determining the effect of dissolved polymers on the sedimentation of GUVs

High concentrations of dissolved polymers in solution could in principle decrease the sedimentation velocity of the GUVs by increasing the viscosity of the solution. The polymers are present at low concentration in the sedimentation chamber.

To demonstrate, we calculate the maximum possible concentration of polymer in the solution containing the harvested GUVs (assuming all the polymers dissolve) using Equation S1. We assume all solutions have the density of water *ρ* = 1 g/mL.

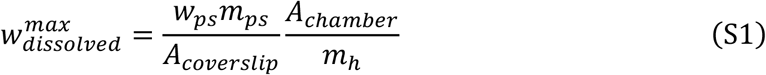

In this equation, 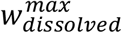 is the maximum concentration (w/w %) of dissolved polymer in the hydrating solution, *w*_ps_ is the concentration of the polymer solution (1 w/w %) deposited on the glass coverslip, *m*_ps_ is the mass of the polymer solution (0.3 mL ≈ 0.3 g) deposited on the glass coverslip, *A*_coverslip_ is the area of the coverslip (484 mm^2^), *A*_cham er_ is the area of the hydration chamber (113 mm^2^), and *m_h_* is the mass of the hydrating buffer (0.15 mL ≈ 0.15 g). 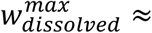 0.47 w/w %. Further, in the sedimentation chamber, the harvested vesicle suspension is diluted 30 times in the sedimentation buffer. The sedimentation buffer is devoid of polymers. Thus, the maximum concentration of dissolved polymers in the sedimentation chamber in the extreme case of complete dissolution is 0.016 w/w%.

The sedimentation chamber that we used has a height of 1000 μm. The vesicles are initially present in all locations in the chamber since they are well-mixed. To test for differences, after 3 hours, we image the solution at the imaging plane (5 μm above the coverslip) and the bulk at 100 μm and 900 μm above the imaging plane for GUVs composed of DOPC harvested from bare glass without any assisting compounds and when LGT agarose is used as the assisting compound. In both cases, we observe a large number of GUVs in the imaging plane and no GUVs in the locations in the bulk (Fig. S1). Our results confirm that the low concentration of polymers, ≪ 0.016 wt %, in the solution in the sedimentation chamber does not have any measurable effect on the sedimentation behavior of GUVs.

### Calculation of molar yield from the literature

Reference^1^ reports that the total number of GUVs with diameters between 10 μm and 80 μm in a 20 μL aliquot from a harvested volume of 500 μL is 63 ± 14. The molar yield is defined as the mols of lipids in GUV membranes divided by the mols of lipids initially used^2^. The mols of lipid initially used in Reference 1 was 1× 10^−9^ mols ^1^. We estimate the maximum possible mols of lipids in the harvested vesicles reported in reference^1^

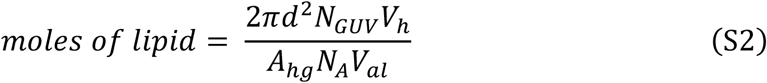

Here *N*_GUV_ is the number of GUVs, taken as 77, *d*_GUV_ is the diameter of the GUVs, taken to be 80 μm, *A*_hg_ is the headgroup area of the lipid which for DOPC is 72.4 × 10^−4^ μm^2^, *N_A_* = 6.023 × 10^23^ is Avogadro’s number, *V_h_* is the harvested volume which is 500 μL, and *V*_al_ is the aliquot counted which is 20 μL. Choosing the largest diameter of GUV reported and the upper range of GUVs counted in the experiment ensures that we are calculating the maximum possible mols of lipids harvested as GUVs from reference 1.

Dividing the moles of lipid in the harvested vesicles with the mols of lipids initially used gives a maximum molar yield of 6.0 × 10^−4^ % for Reference 1. Notably, the diameters of GUVs quantified in Reference 1 were limited to 10 μm ≤ *d* < 100 μm ^1^. When we similarly limit our measurements, we find that the yield of GUVs from the fructose-doped technique that we performed is 3.9 × 10^−2^ %. This result is ∼ 2 orders of magnitude greater than that calculated from reference 1. Nevertheless, these values show that the fructose-doped method produces a very low yield of GUVs in physiological salt solutions.

### Energy for forming spherical buds from flat surfaces

The free energy of a budding membrane that is templated on a surface is modeled by Equation S3 ^2^.

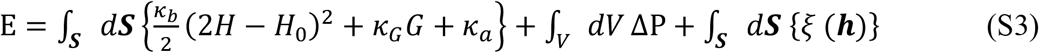

In this equation, *κ_b_* is the bending modulus, *κ_G_* is the Gaussian bending modulus, 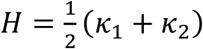 is the mean curvature where *κ* and *κ*_2_ are the principal curvatures on the surface, *G* = *κ κ*_2_ is the Gaussian curvature, ***h***_0_ is the spontaneous curvature, *κ_a_* is the area expansion modulus, Δp is the difference in osmotic pressure, and *ξ*(***h***) is the microscopic interaction potential normal to the surface of the membrane. The magnitude of *ξ*(***h***) depends on the distance, ***h***, between the membrane and the surface^3^. These quantities are integrated as appropriate over the surface, ***S***, of the membrane, or the volume, *V*, that the membrane encloses.

Equation S3 can be simplified by noting that ***h***_0_ = 0 for a symmetric bilayer and that there is no change in the Gaussian curvature for spherical buds that remain attached to the surface^2^. We further simplify by replacing the microscopic interaction potential, *ξ*(***h***), with an effective adhesion contact potential, *ξ* ^2^. We obtain Equation S4 by i) integrating Equation S3 to obtain the energy for State 1, the geometry of a flat disk of radius ***R**_d_*, the energy for State 2, the geometry of a spherical bud of radius 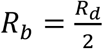, and ii) subtracting the energy of State 1 from State 2.

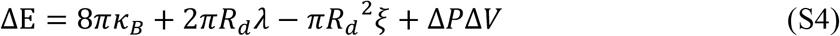

The second term on the RHS of Equation S4 introduces a constraint for a section of the membrane to transition into a spherical bud at a constant area. If there is a lipid source, the membrane can transition without requiring breaks by recruiting lipids from the source^3^. In the absence of a lipid source, the membrane must form breaks, with an edge energy *λ*, to allow the lipids to reconfigure to form a spherical bud. In the main manuscript, for simplicity, we assume that the membrane has a source and does not break during budding, thus *λ* = 0. For gentle hydration on bare glass, Δ*p* = 0. For polymer-coated surfaces Δ*p* depends on the concentration of dissolved polymer.

### Estimate of the osmotic pressure contribution of the dissolved polymer

The behavior of the polymer and lipid is expected to be far from ideal due to the small length scales, the high concentrations, the large molecular weight of the polymer, and the time dependent process of lipid hydration and polymer dissolution. The polymers appear to interact with the lipid, which further changes the behavior. To make progress, we consider a simplified scaling argument. We assume ideal behavior and use characteristic length and energy scales to determine if the osmotic pressure exerted by the dissolving polymer at the concentrations that we use in our experiments can overcome an increase in adhesion between the membranes. Specifically, we estimate the concentration of dissolved polymer that is needed in the interlamellar space of the bilayer stacks so that the contribution of the osmotic pressure results in no change in the free energy between polymer-free budding in low salt solutions and polymer-assisted budding in salty solutions.

We assume that the effects of the salt and polymers on the edge energy, *λ*, and the bending rigidity, *κ_B_*, to be negligible. The change in energy in the low-salt condition, Δ*E_LS_* and salty condition, Δ*E_HS_* is given by Equations S5 and S6.

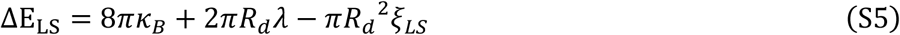

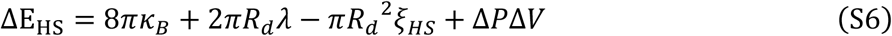

In these equations, *ξ_LS_* is the adhesion energy in the low salt solution, *ξ_HS_* is the adhesion energy in the salty solution, Δ*P* is the osmotic pressure difference, assumed to be constant, and *ΔV* is the change in volume.

The change in volume from a disk-shaped bilayer on the surface with radius *R_d_* and interlamellar spacing height, *z* to a spherical bud of radius *R_b_* is given by Equation S7a

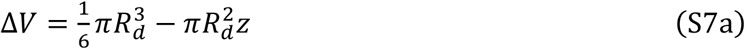

We used 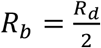 for the transition between a disk to a spherical bud at a constant surface area to express Equation S3a in terms of *R_d_*.

For an ideal small molecule osmolyte at low concentrations, Δ*P* is related to the concentration of the osmolyte, *c* by Equation S7b.

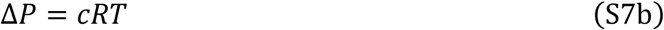

In this equation, *R* is the ideal gas constant, and *T* is the temperature.

We obtain Equations S7c and S7d which satisfies the condition that there be no change in energy between assembly of GUVs in the low salt solution and the salty solution.

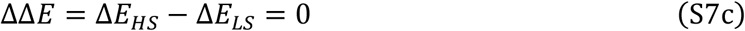

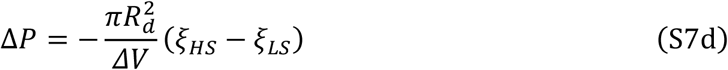

We substitute Equation S7a and S7b into equation S7d to get Equation S8. Equation S8 relates the concentration of the polymer in the interlamellar space that is needed to balance the increase in the adhesion energy in the salty solution compared to the low salt solution.

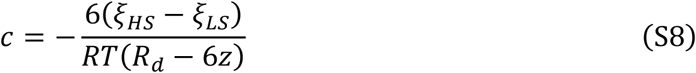

We take *ξ*_LS_ = 1 × 10^−6^ J m^−2^ for DOPC membranes in low-salt solutions and *ξ*_HS_ = 1 × 10^−4^ J m^−2^ for DOPC membranes in salty solutions^3^. We use *z* = 4 nm and *R_d_* = 1 μm for a GUV bud 1 μm in diameter. We get 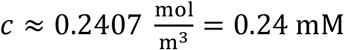. The use of the ideal expression for osmotic pressure in the dilute limit does not change our conclusion that low amounts dissolved polymer is sufficient to exert an osmotic pressure that balances the adhesion energy in high salt solutions. For macromolecular polymers at moderate concentrations, the osmotic pressure is often expressed as Δ*P* = (*c* + *A*_2_(*Mc*)^2^)*RT*, where *A*_2_ is the second virial coefficient and *M* is the molecular weight of the polymer^4^. The magnitude of the osmotic pressure, Δ*P*, is thus expected to be higher for polymers compared to small molecule solutes for the same dissolved concentration, *c*. A lower amount of dissolved polymer than predicted by Equation S8 will be sufficient to balance the increased adhesion between membranes in salty solutions.

We next consider the expected concentration of dry polymer, *c_d_*, in the interlamellar volume of the bilayer stack. The mass of polymer molecules per unit area of substrate (0.003 g of the polymer spread over a coverslip area of 4.84 × 10^−4^ m^2^) is 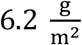. We assume that upon hydration the polymer and lipid form uniform stacks composed of 5 lipid bilayers with an interlamellar spacing of 4 nm. The concentration of dry polymer in the stack with a molecular weight *M* = 120,000 is,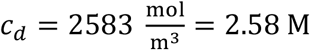. Thus, in this ideal calculation, approximately 0.009 % of the polymer must dissolve to form GUV buds in salty solutions.

### Electrostatic interaction between membranes

The adhesion of membranes is affected by the presence of ions in solutions^5–7^. Ions promote adhesion by screening repulsive electrostatic interactions. The effectiveness of screening can be estimated using the Debye screening length^3^. Smaller screening lengths reflect a shorter range in which electrostatic repulsion is felt, thus allowing attractive van der Waals interaction to dominate^3^. We use Equation S9^3^ to calculate the

Debye screening length, 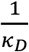.

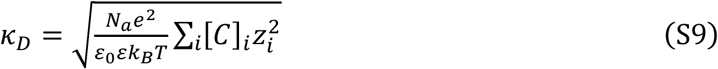

Here *N_A_* is Avogadro’s number, *e* is the elementary charge, [*C*] is the concentration of ionic species *i, z* is the charge of ionic species *i, ε*_0_ is the permittivity of free space, *ε* is the dielectric constant, *k_B_* is the Boltzmann constant, and *T* is the absolute temperature.

We report the ionic composition of the buffers used for the experiments in Figure 9 in Table S10. We report the ionic species and the calculated Debye screening lengths in Table S11. Note that the solutions consisting of 100 mM sucrose and 140 mM KCl + 5 mM CaCl_2_ are unbuffered. Our measurements showed that they have a pH of 5.5. This value of pH is consistent with the formation of carbonic acid when ultrapure water is in equilibrium with atmospheric carbon dioxide^8^. The screening length drops from 170 nm in the solution consisting of 100 mM sucrose to 0.75 nm in the solution consisting of PBS + 100 mM sucrose. The screening length is further reduced to 0.39 nm in the solution consisting of 600 mM NaCl + 100 mM sucrose. The calculated Debye screening lengths of PBS + 5 mM MgCl_2_ and 140 mM KCl + 5 mM CaCl_2_ are 0.73 nm and 0.77 nm respectively. Although the values of the Debye lengths are similar to those of PBS, adhesion is significantly enhanced because mM concentration of divalent cations can bind and neutralize surface charges and serve as ionic bridges between charges^9,10^.

## Supporting Figures

**Figure S1.**
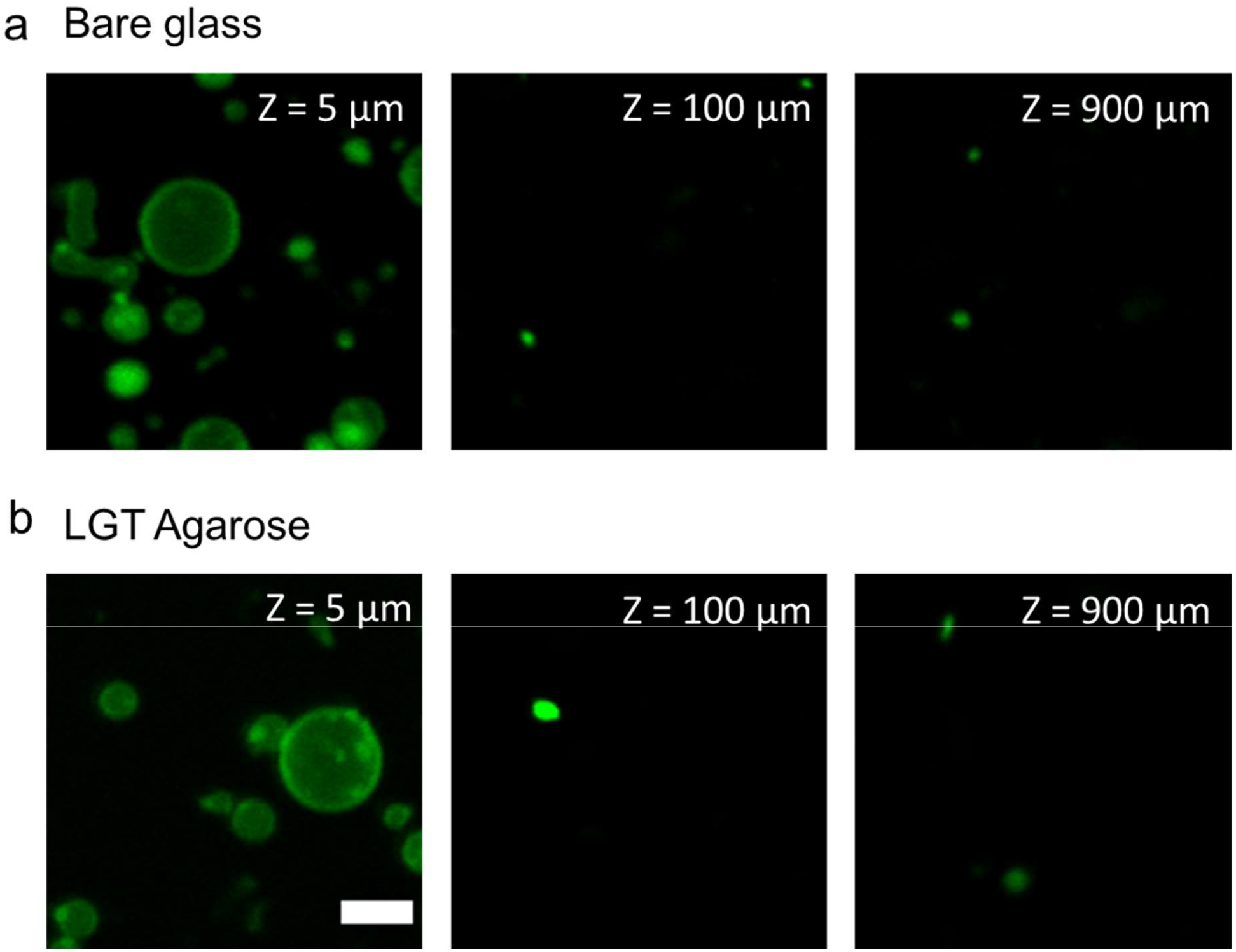
GUVs sediment in imaging chambers similarly in the presence and absence of polymers. Representative images at various *Z* planes of our imaging chamber after 3 hours of sedimentation. a) GUVs obtained from a bare glass surface without the use of assisting compounds. b) GUVs obtained when LGT agarose was used as the assisting compound. The Z position relative to the location of the surface of the bottom glass slide of the imaging chamber is indicated in the images. The scale bar is 10 μm.

**Figure S2.**
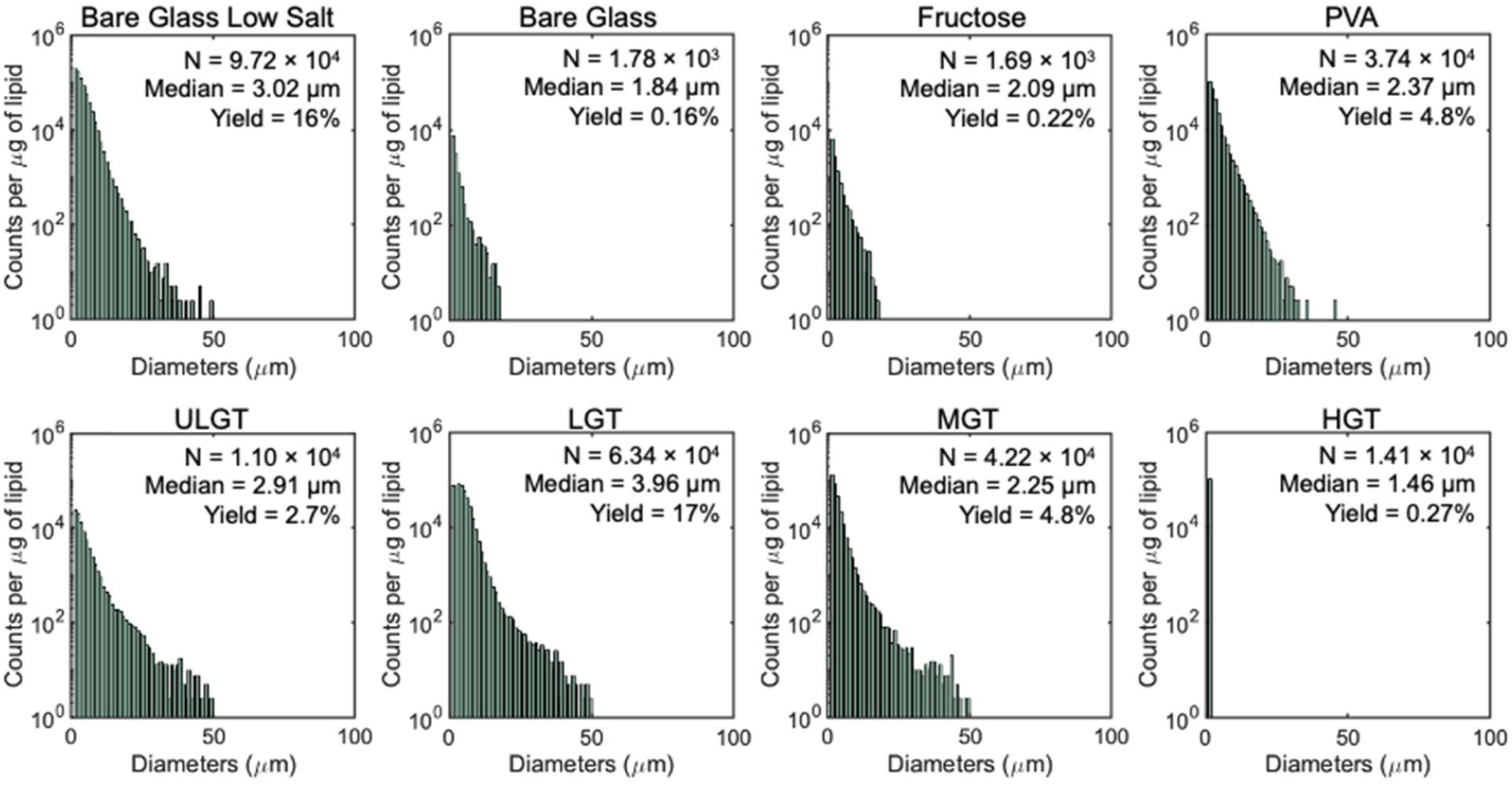
Histograms of GUV diameters of samples shown in Figure 2. Each histogram is the average of 3 independent repeats per sample. Note the logarithmic scale on the y-axis. Bin widths are 1 μm.

**Figure S3.**
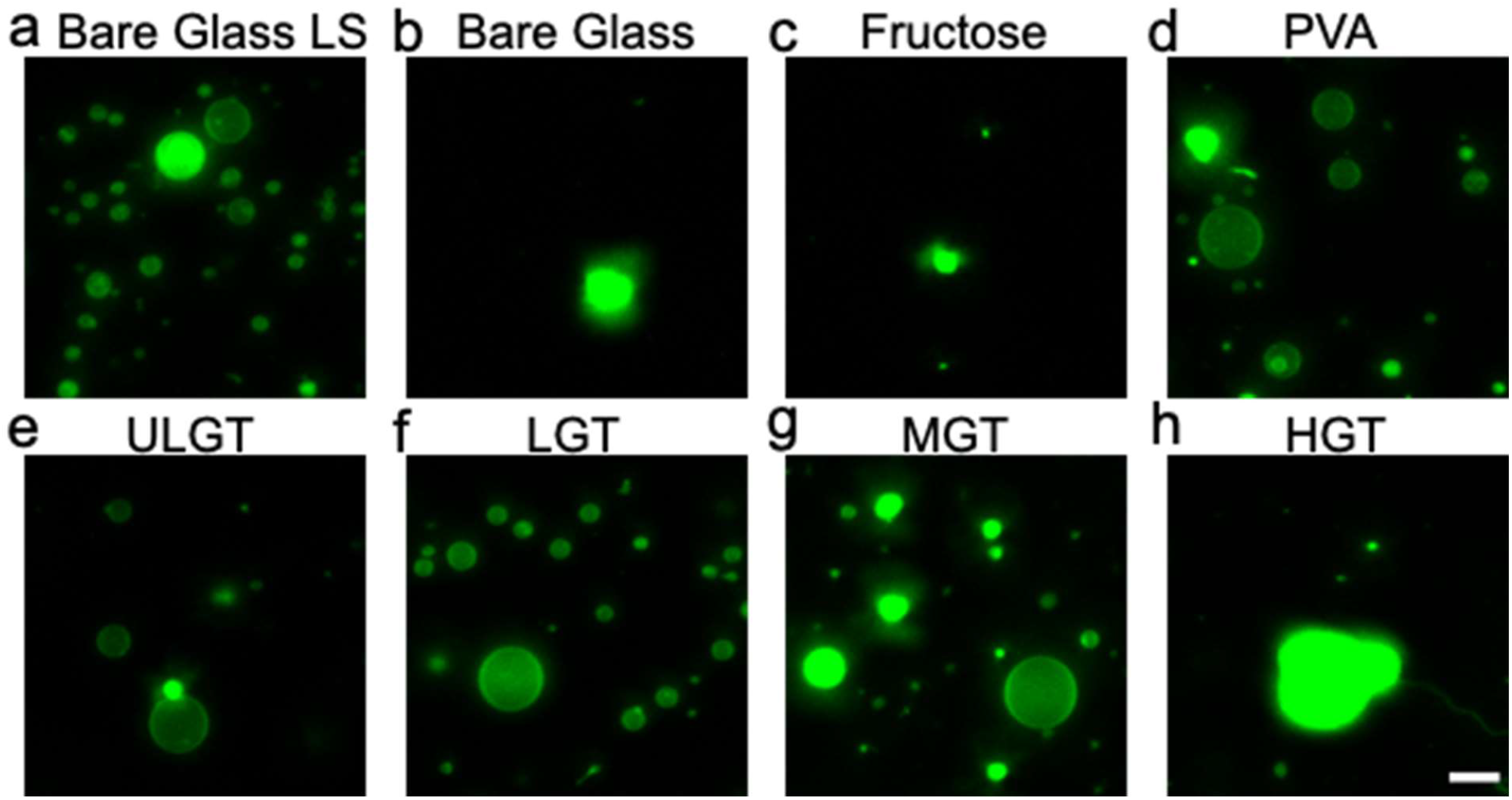
Representative images of the harvested objects for samples shown in Figure 2. Samples hydrated at 22 °C. Bright spots in all samples are lipid aggregates or MLVs. The scale bars are 15 μm.

**Figure S4.**
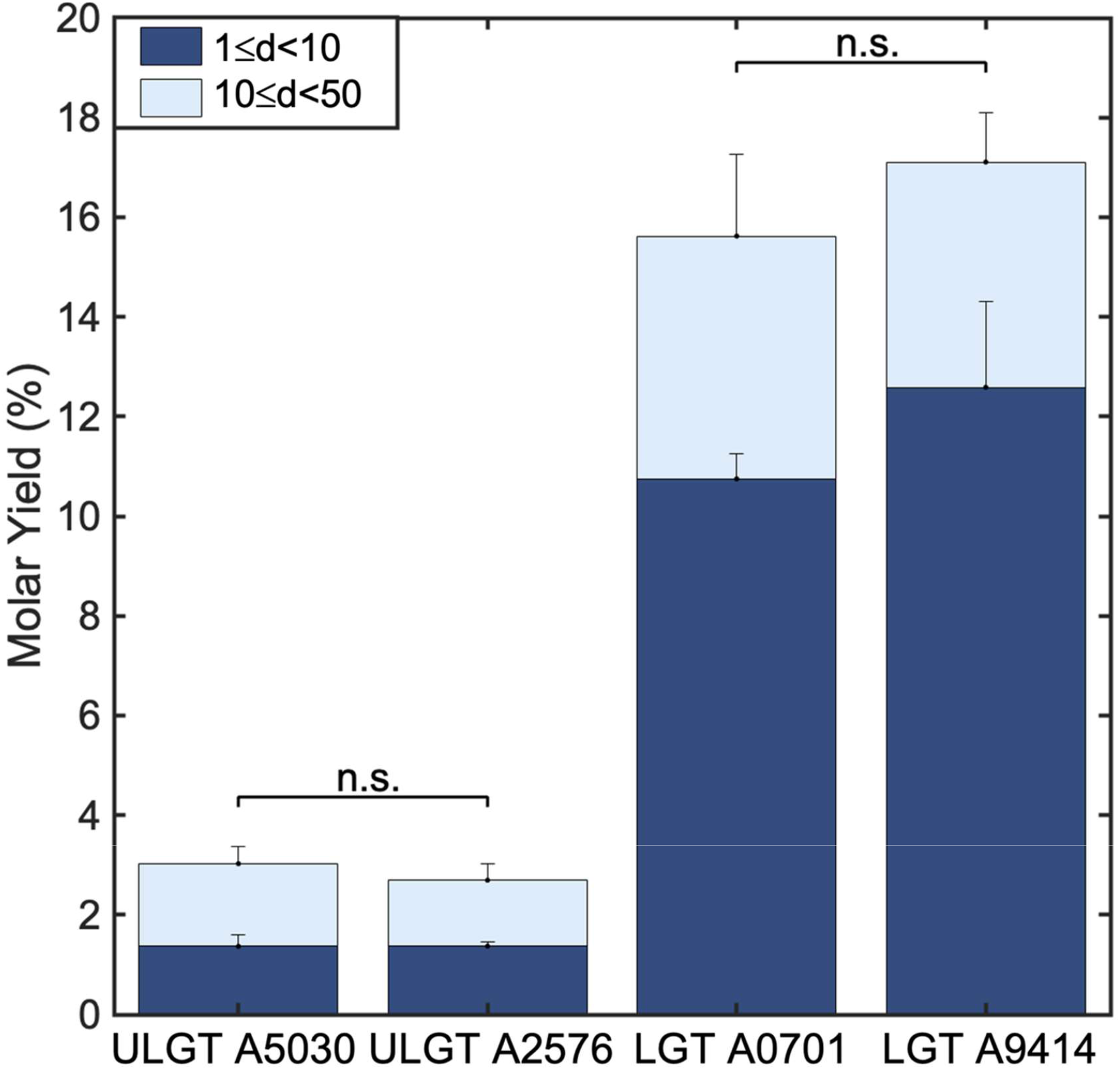
There is no significant difference in yields of GUVs obtained from ULGT and LGT agaroses with different catalog numbers (ultra-low gelling: A5030, A2576 and low gelling: A0701, A9414). Each bar is an average of 3 independent repeats per sample. Statistical significance determined by t-test. * = p < 0.05, ** = p < 0.01, *** = p < 0.001, ns = not significant. The data for A2576 and A9414 are from Figure 2.

**Figure S5.**
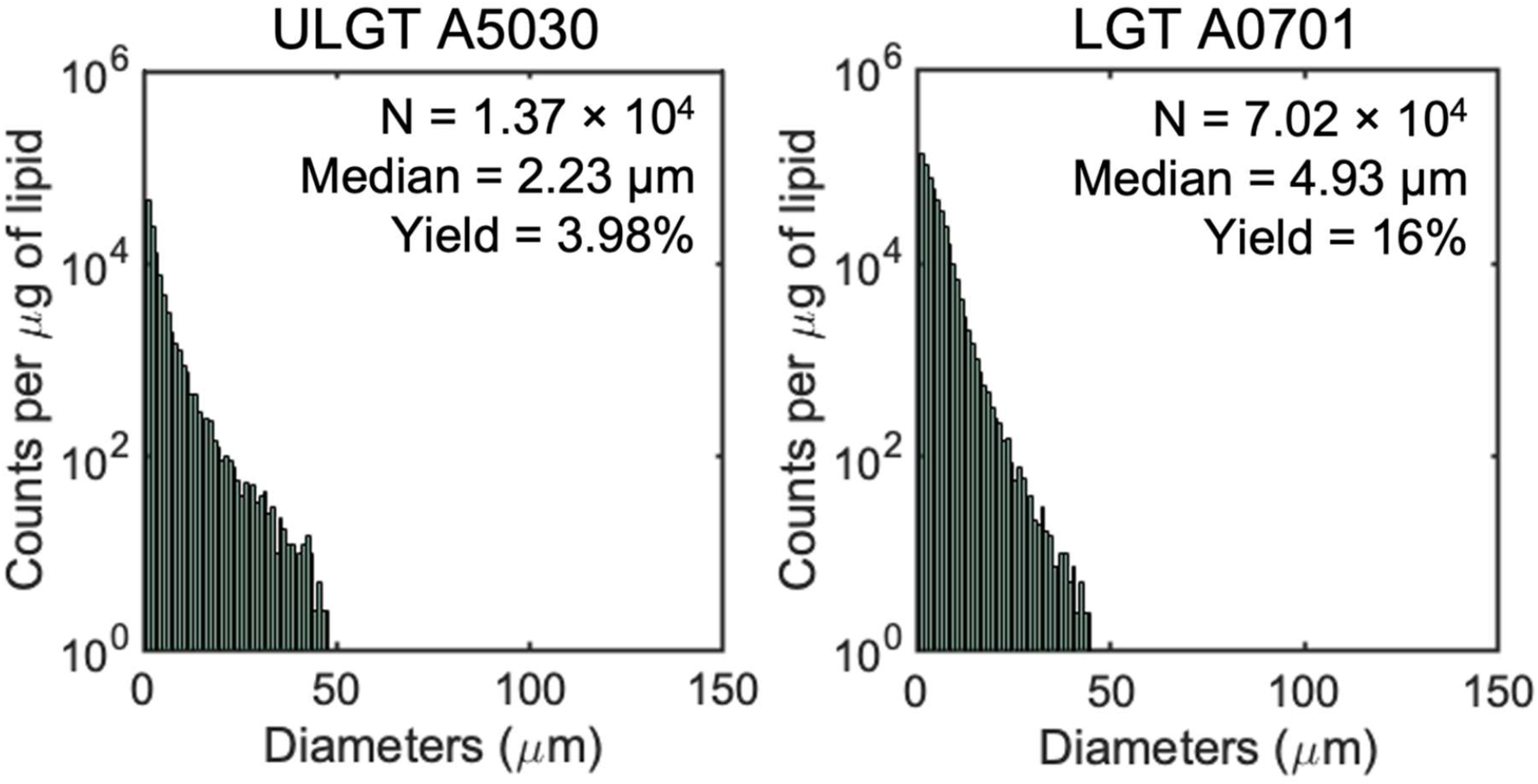
Histograms of GUV diameters of the samples shown in Figure S4. Each histogram is the average of 3 independent repeats per sample. Note the logarithmic scale on the y-axis. Bin widths are 1 μm.

**Figure S6.**
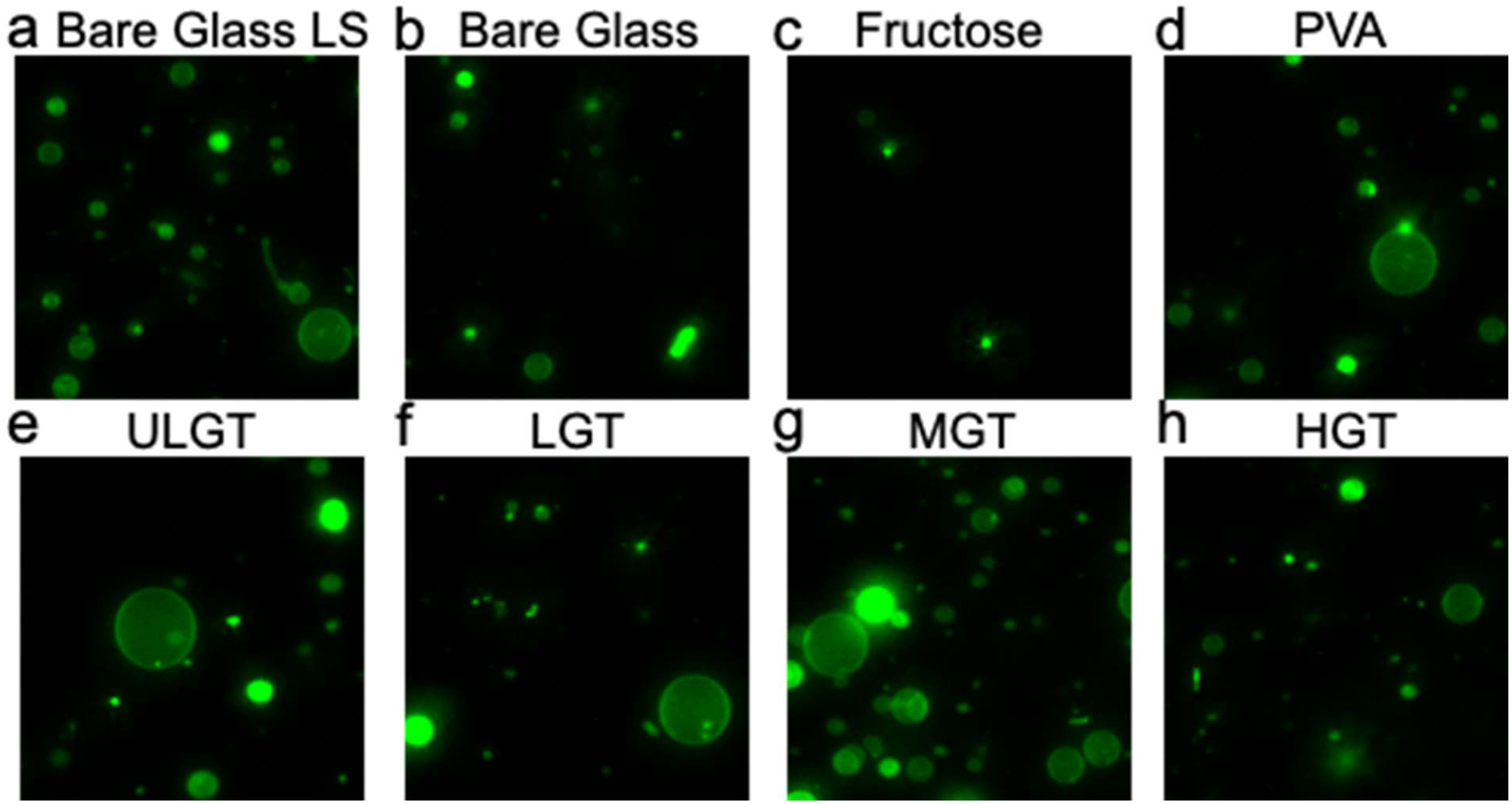
Representative images of the harvested objects for samples shown in Figure 3. Samples hydrated at 37 °C. Bright spots in all samples are lipid aggregates or MLVs. The scale bars are 15 μm.

**Figure S7.**
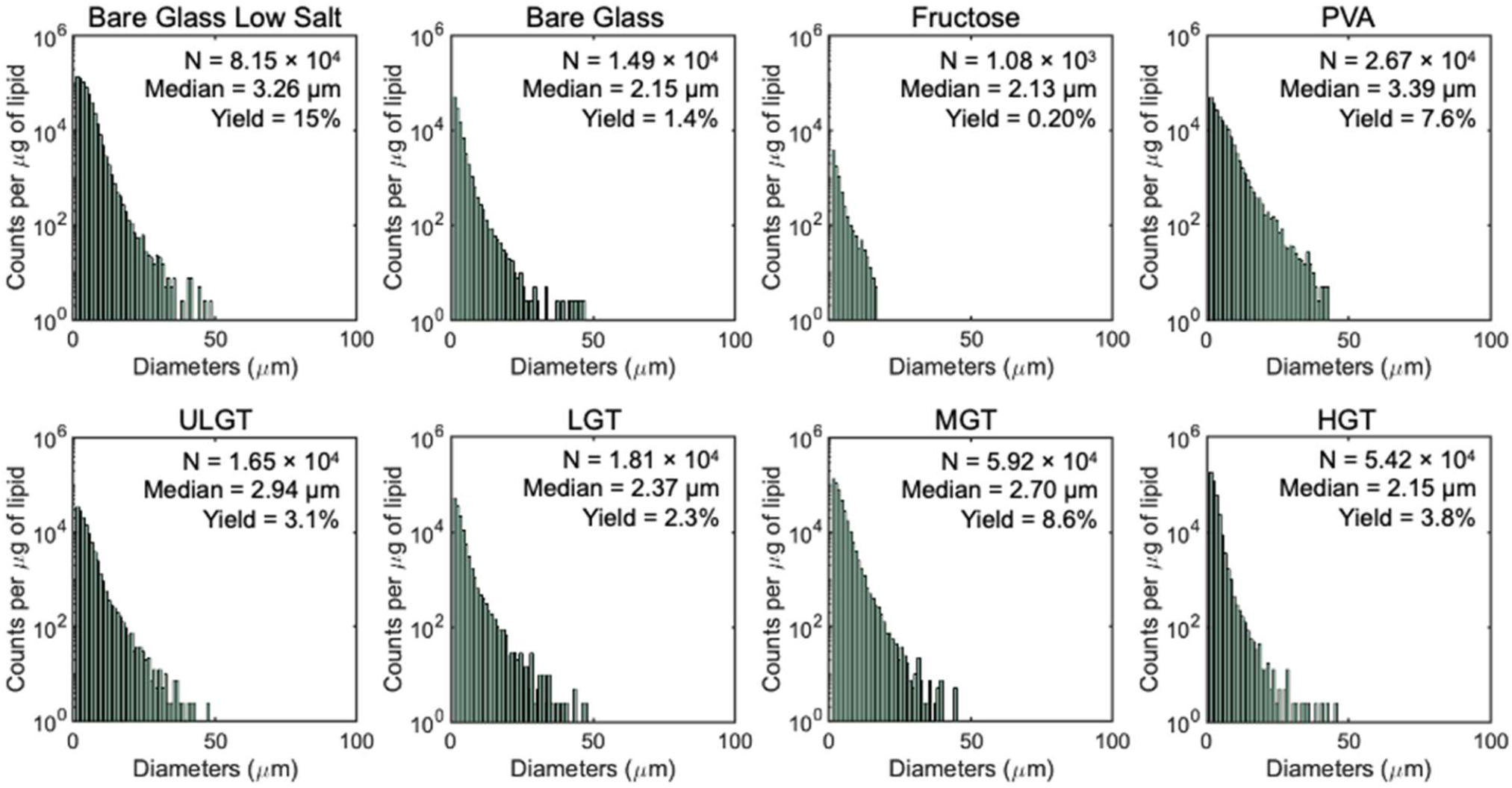
Histograms of GUV diameters of samples shown in Figure 3 for GUVs assembled at 37 °C. Each histogram is the average of 3 independent repeats per sample. Note the logarithmic scale on the y-axis. Bin widths are 1 μm.

**Figure S8.**
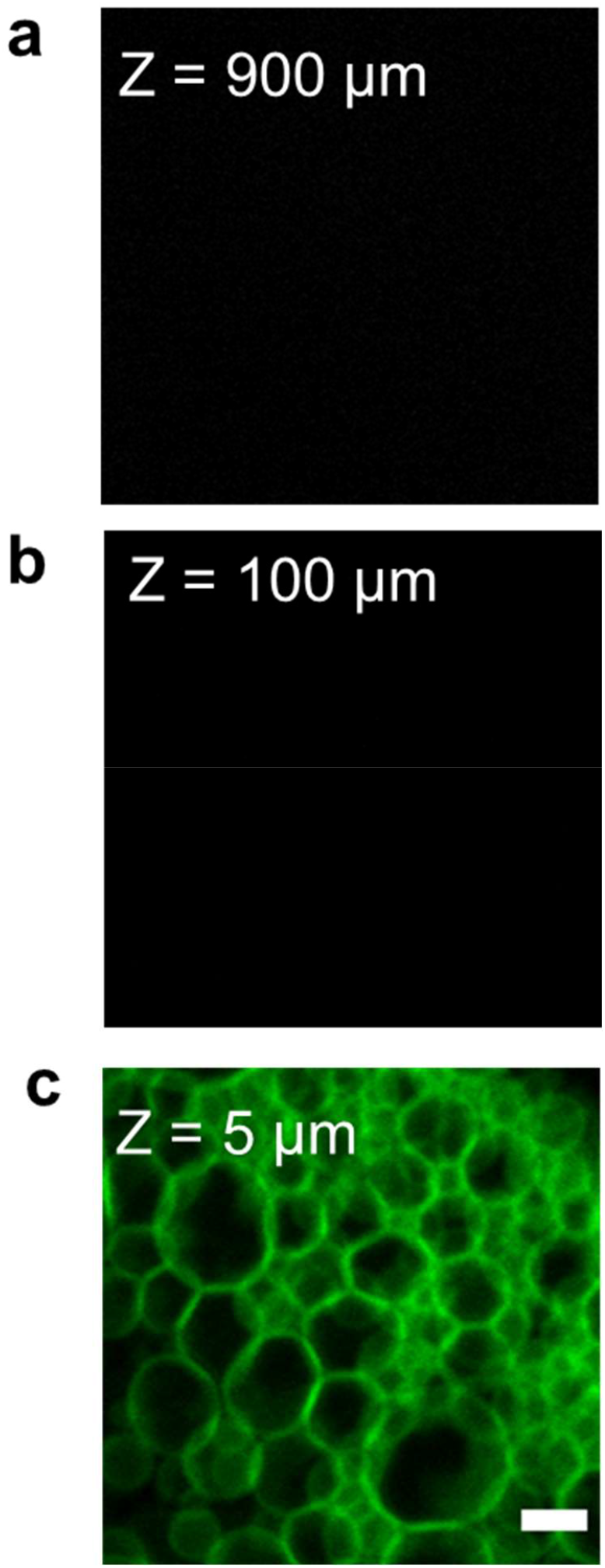
GUV buds remain attached to the surface in the absence of flow. Representative images after 2 hours of incubation at a) 900 μm and b) 100 μm above the LGT-agarose coated coverslip. Both regions had few to no floating structures. c) The surface of the agarose is covered with a high density of buds. The supporting glass coverslip is at Z = 0 μm. The scale bar is 10 μm.

**Figure S9.**
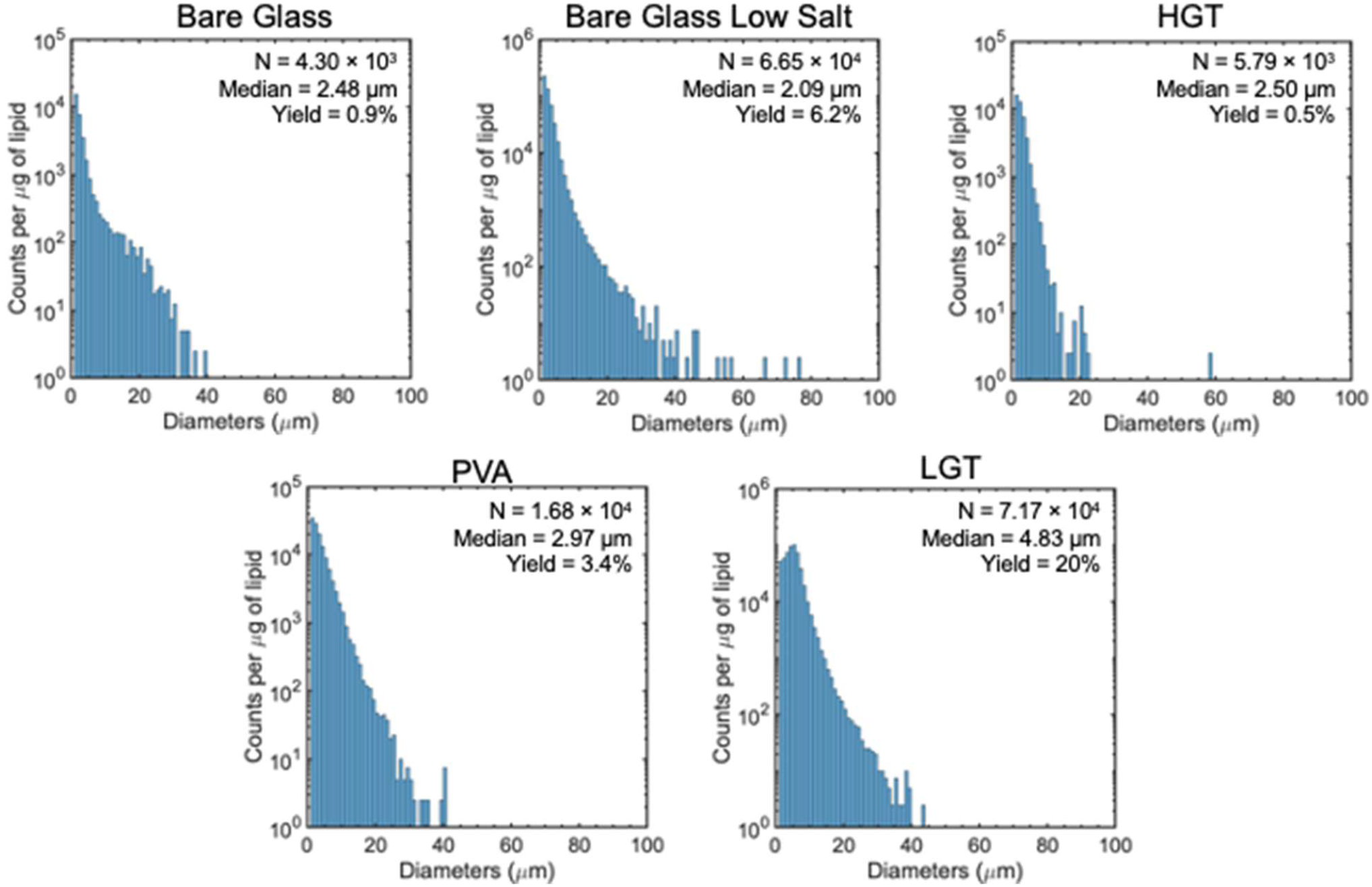
Histograms of GUV diameters of the samples shown in Figure S8a. The membranes of the GUVs are composed of a lipid mixture that minimally mimics the composition of the exoplasmic leaflet of the mammalian cellular membrane (mammalian exoplasmic leaflet (MEL)). Each histogram is the average of 3 independent repeats per sample. Note the logarithmic scale on the y-axis. Bin widths are 1 μm.

**Figure S10.**
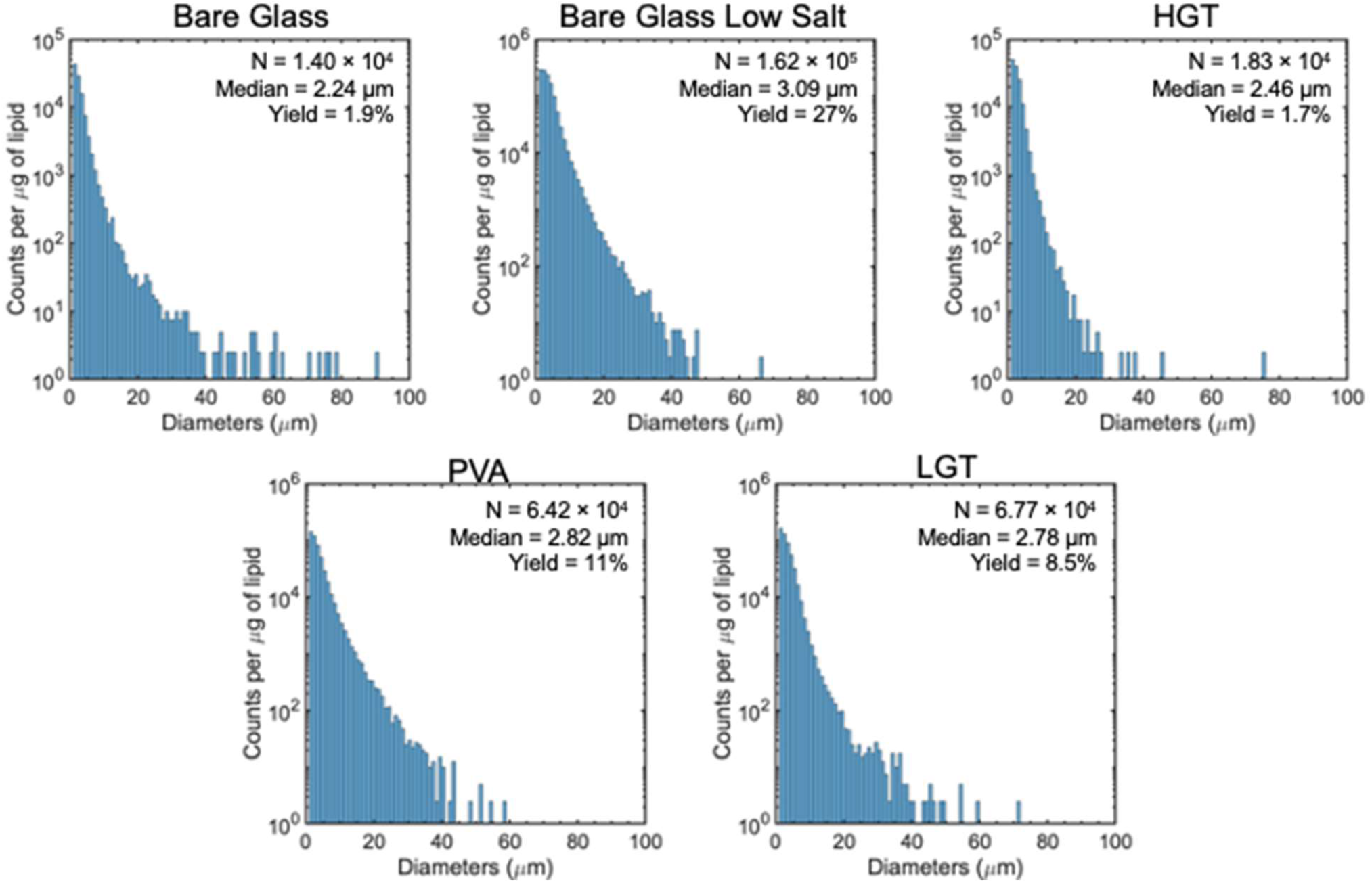
Histograms of GUV diameters of the samples shown in Figure S8b. The membranes of the GUVs are composed of a lipid mixture that minimally mimics the composition of the endoplasmic-reticulum-Golgi intermediate compartment (ERGIC) membrane. Each histogram is the average of 3 independent repeats per sample. Note the logarithmic scale on the y-axis. Bin widths are 1 μm.

**Figure S11.**
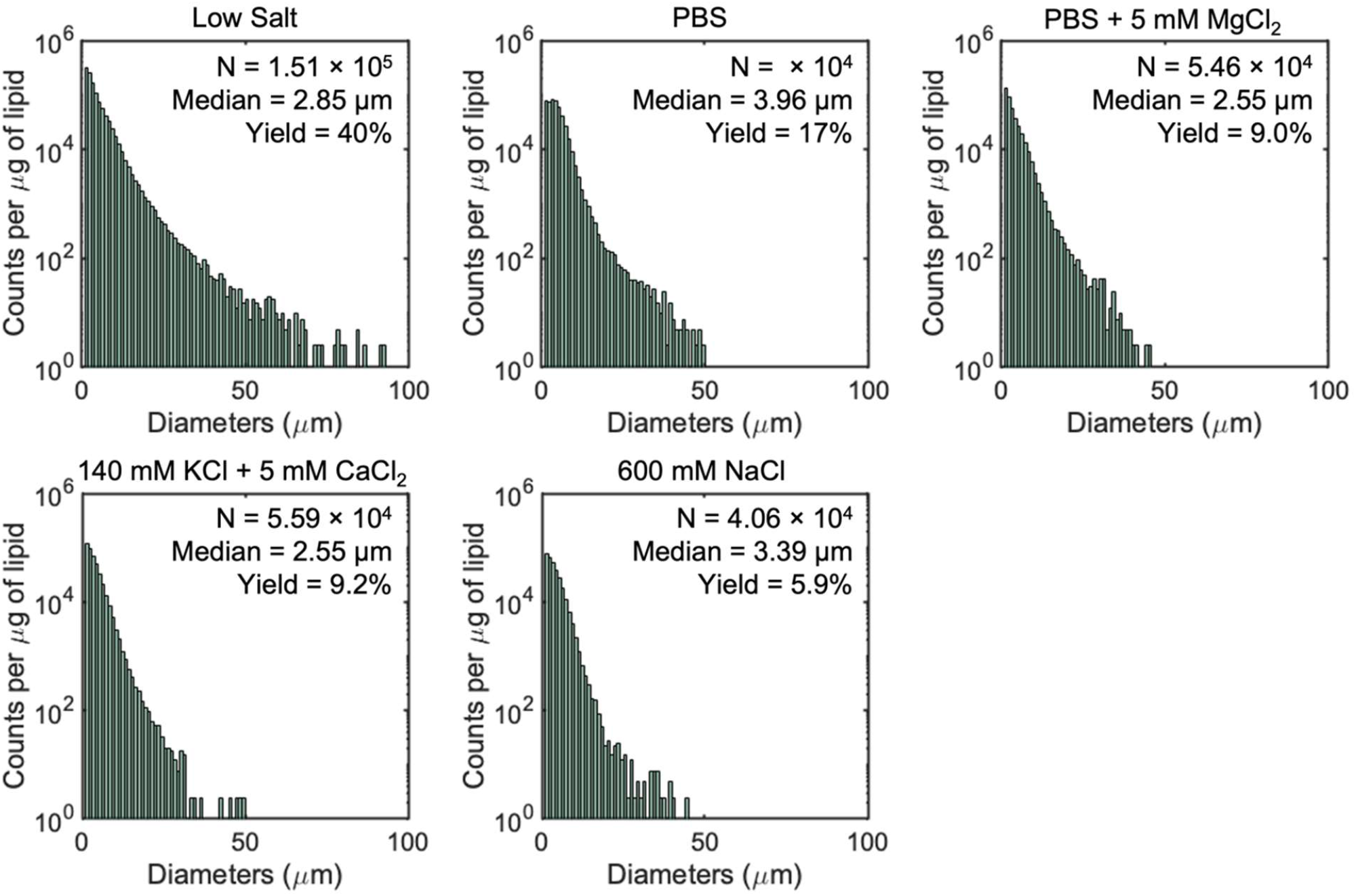
Histograms of GUV diameters of samples shown in Figure 9. The membranes of the GUVs are all composed of DOPC and the assisting compound is low gelling temperature (LGT) agarose. Each histogram is the average of 3 independent repeats per sample. Note the logarithmic scale on the y-axis. Bin widths are 1 μm.

## Supporting Tables

**Table S1.**
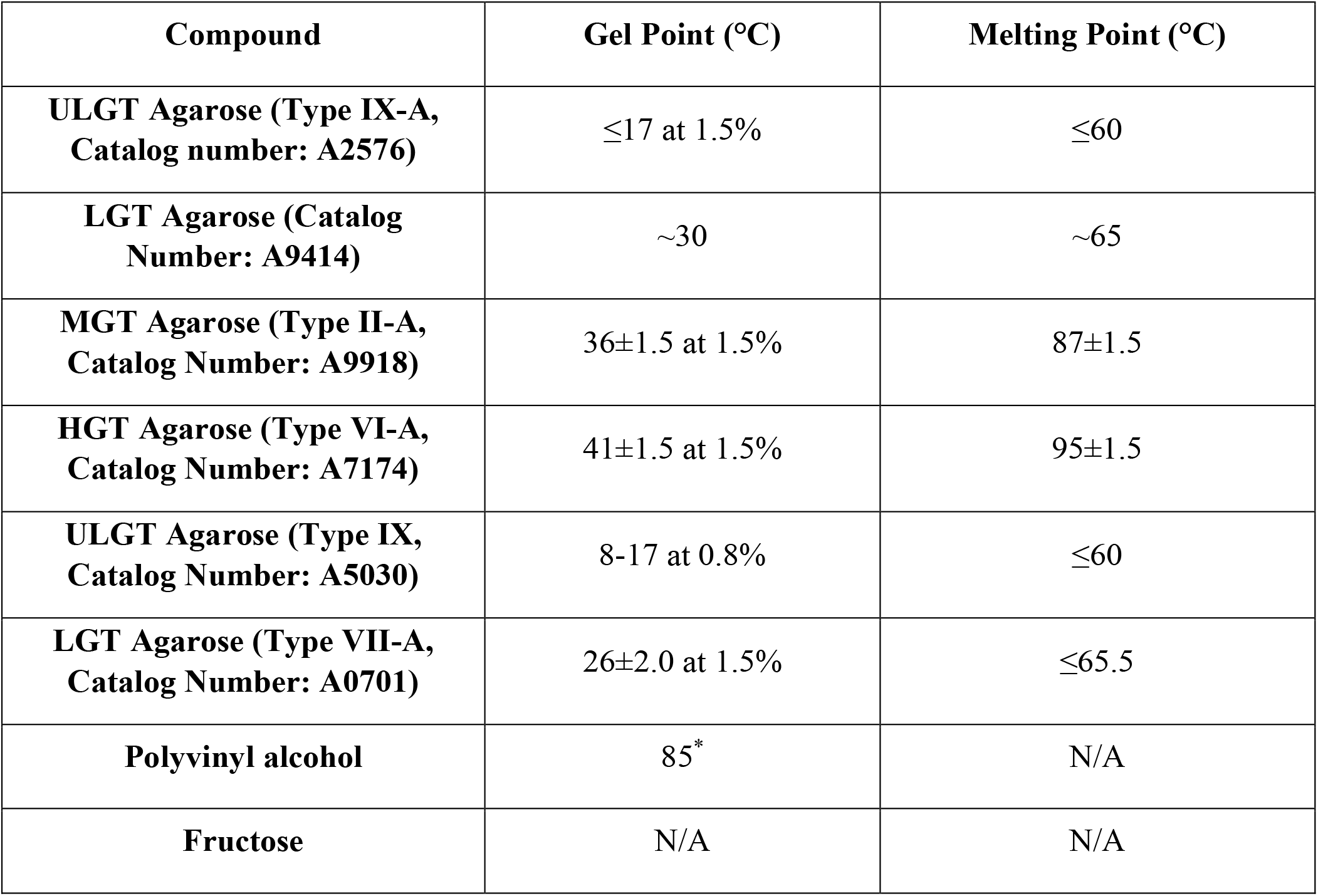
Melting and gelling temperatures of the compounds tested. **^*^**Glass transition temperature. Values for agarose are from ^11^, values for PVA is from ^12^.

**Table S2:**
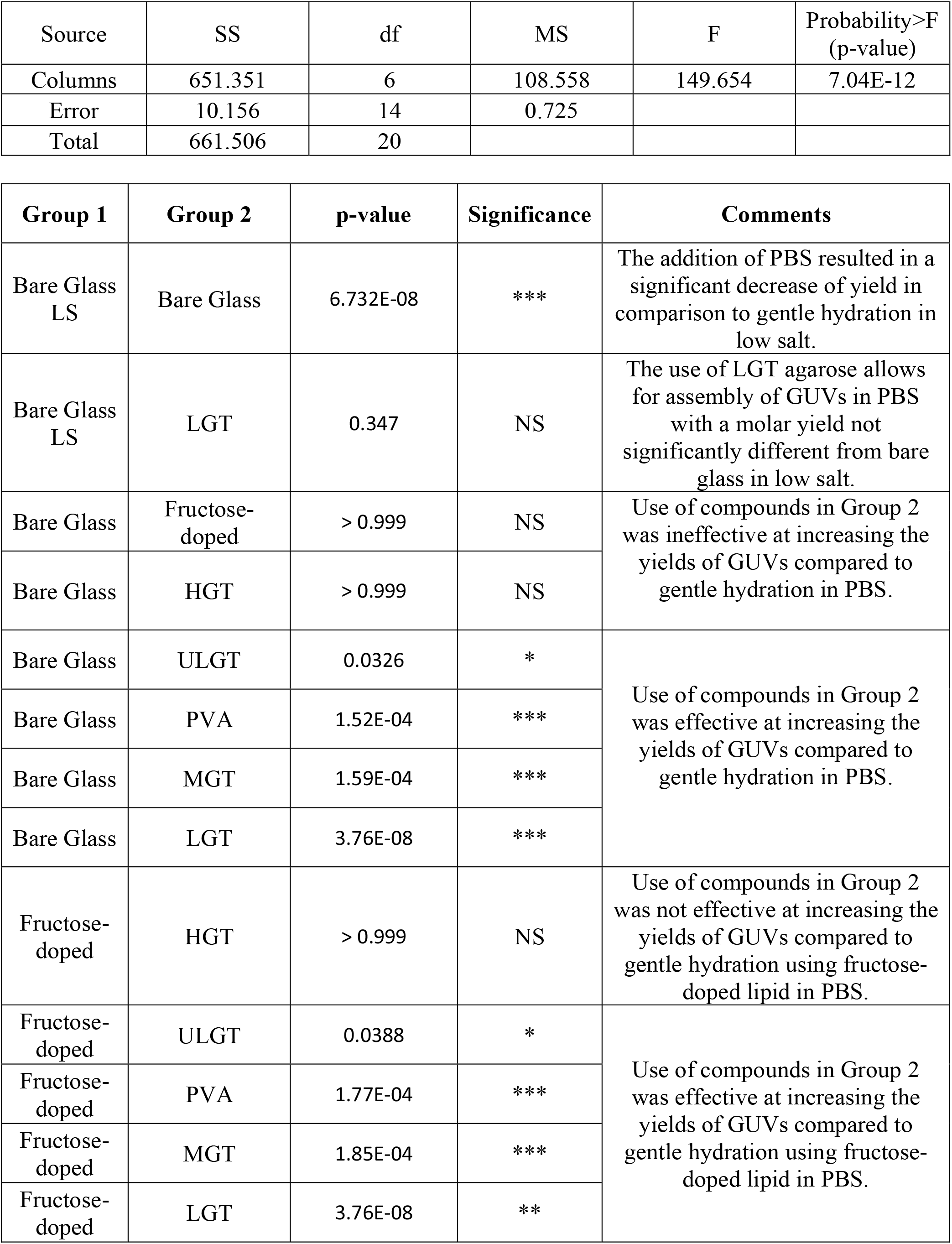
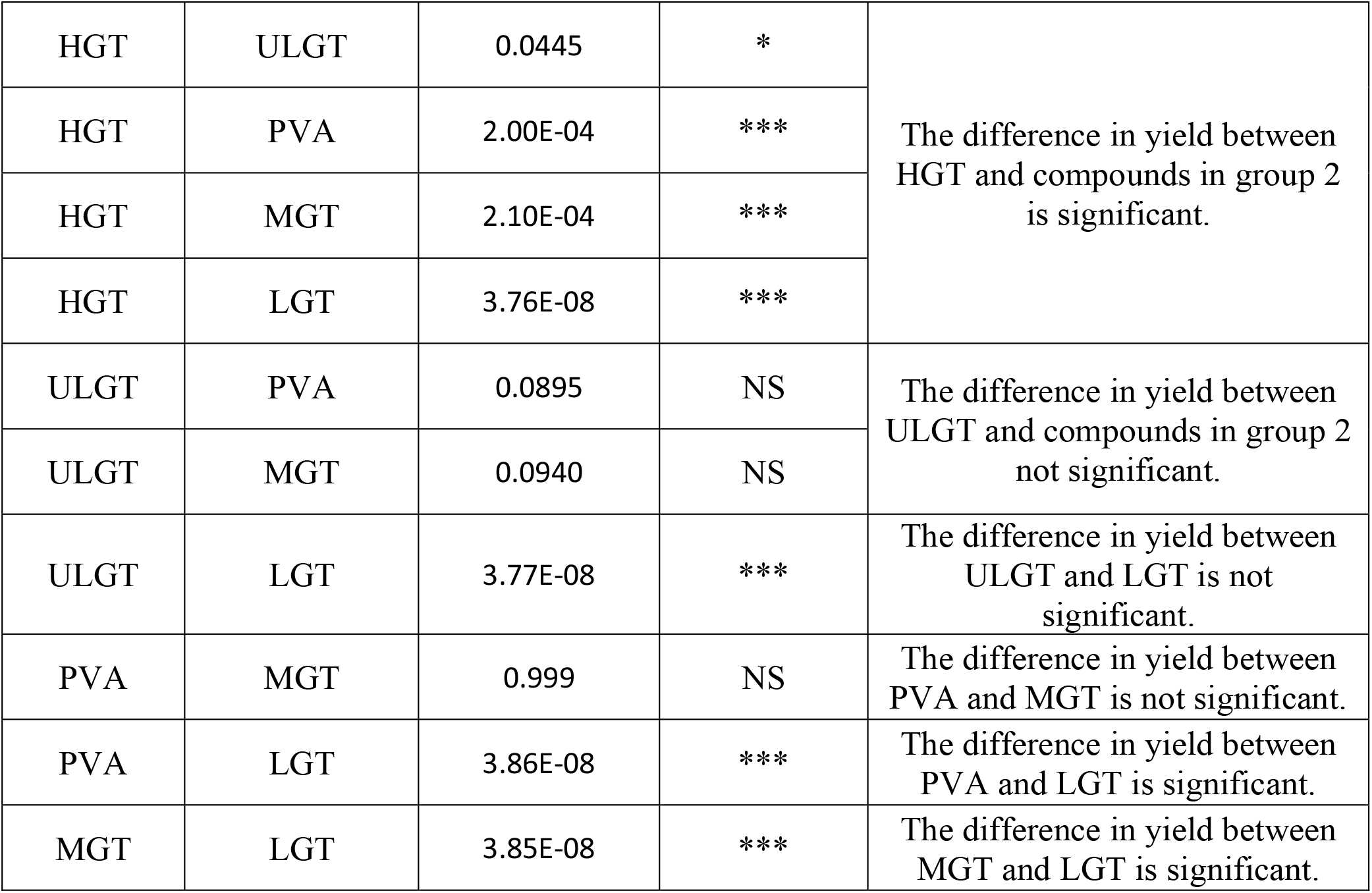
ANOVA table and table of p-values from post hoc Tukey’s HSD tests of the molar yields of GUVs assembled at 22 °C obtained from gentle hydration on bare glass in PBS (Bare Glass), fructose-doped lipid in PBS, and hydration on polymer films of PVA, ULGT agarose, LGT agarose, MGT agarose, and HGT agarose in PBS. Comparison of gentle hydration on bare glass in low salt (Bare Glass LS) and gentle hydration on bare glass in PBS (Bare Glass) was conducted separately from the ANOVA via Student’s t-test. All solutions contain 100 mM of sucrose. * = p < 0.05, ** = p < 0.01, *** = p < 0.001, NS = not significant.

**Table S3:**
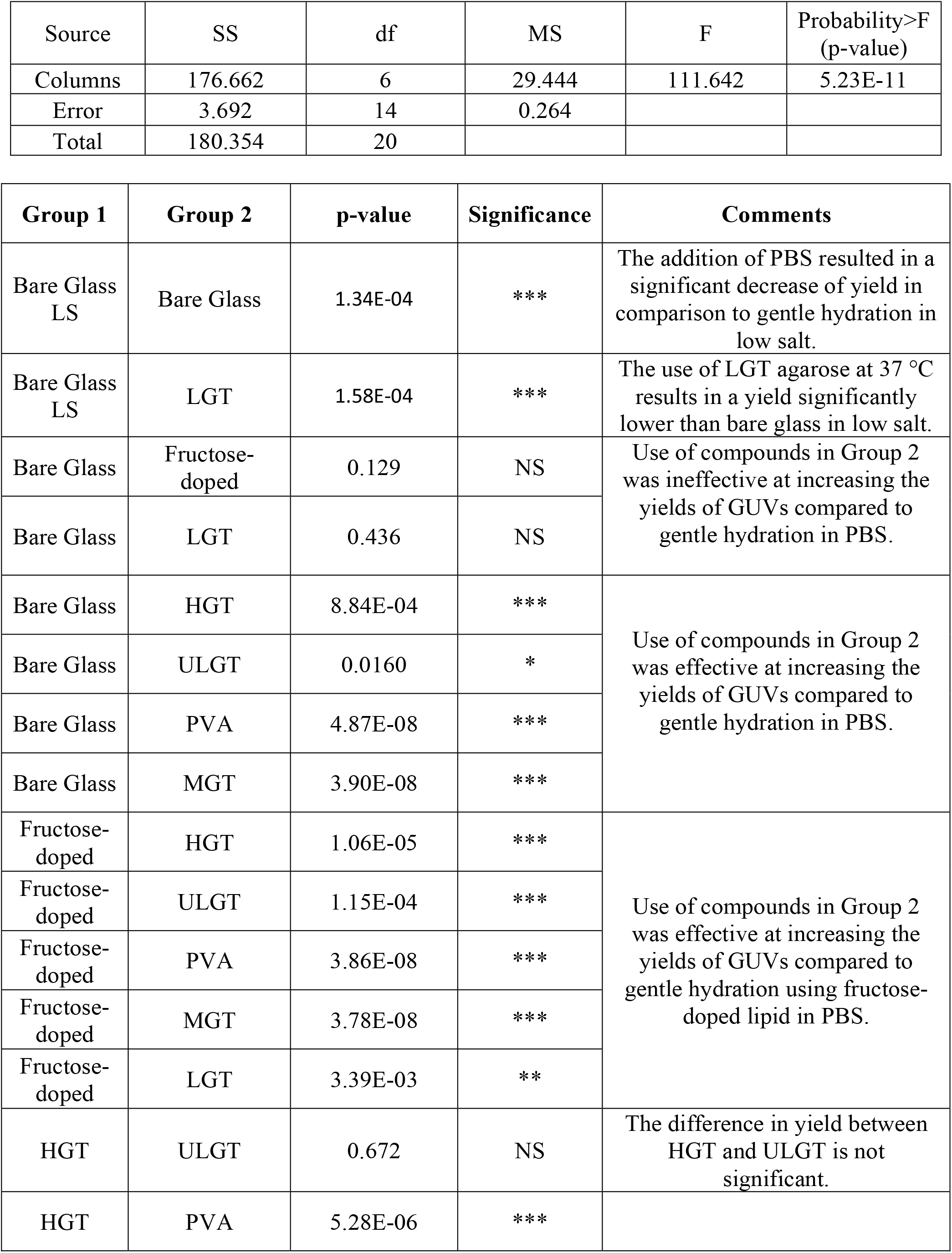
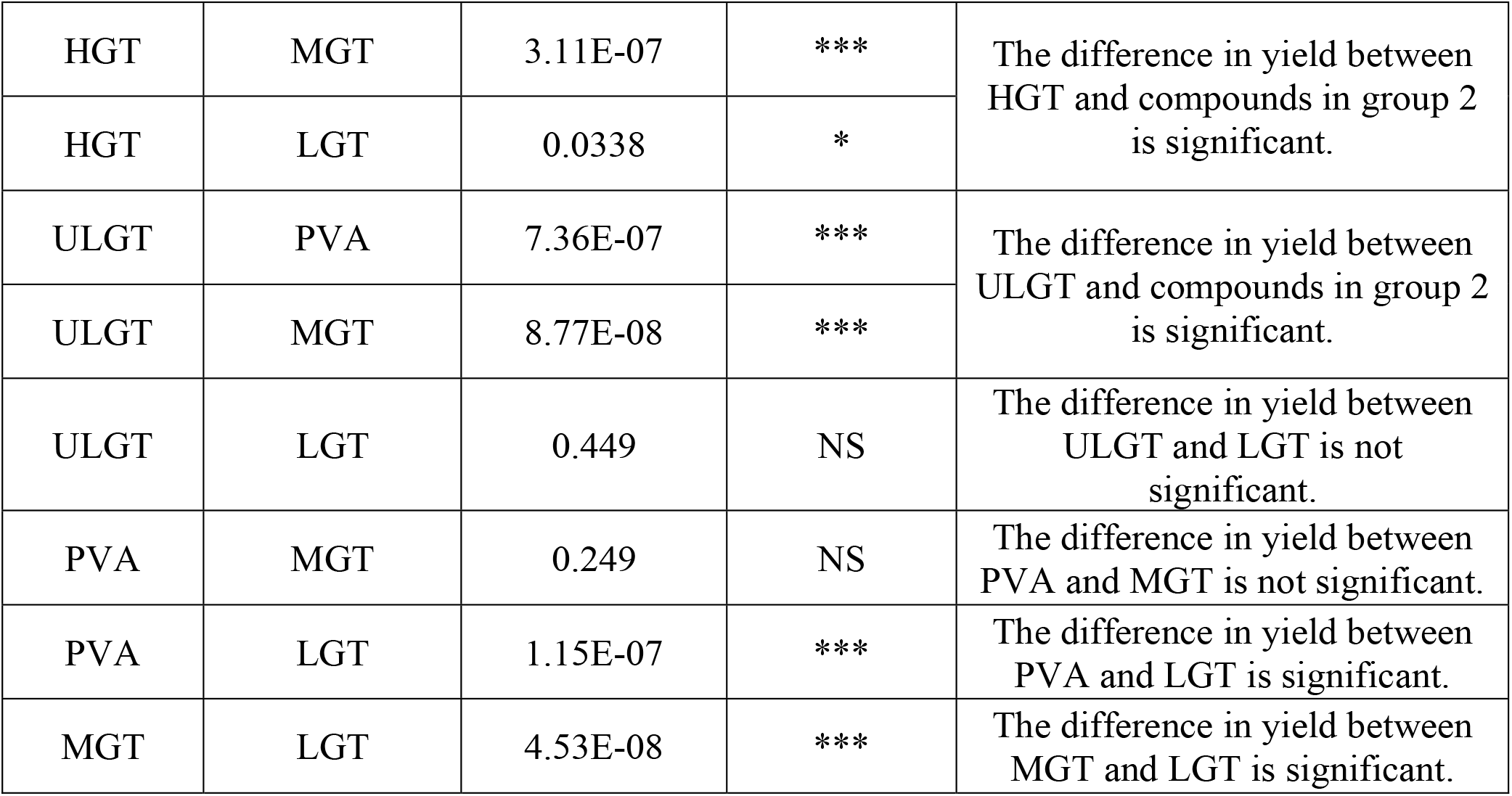
ANOVA table and table of p-values from post hoc Tukey’s HSD tests of the molar yields of GUVs grown at 37 °C obtained from gentle hydration on bare glass in PBS (Bare Glass), fructose-doped lipid in PBS, and hydration on polymer films of PVA, ULGT agarose, LGT agarose, MGT agarose, and HGT agarose in PBS. Comparison of gentle hydration on bare glass in low salt (Bare Glass LS) and gentle hydration on bare glass in PBS (Bare Glass) was conducted separately from the ANOVA via Student’s t-test. All solutions contain 100 mM of sucrose. * = p < 0.05, ** = p < 0.01, *** = p < 0.001, NS = not significant.

**Table S4:**
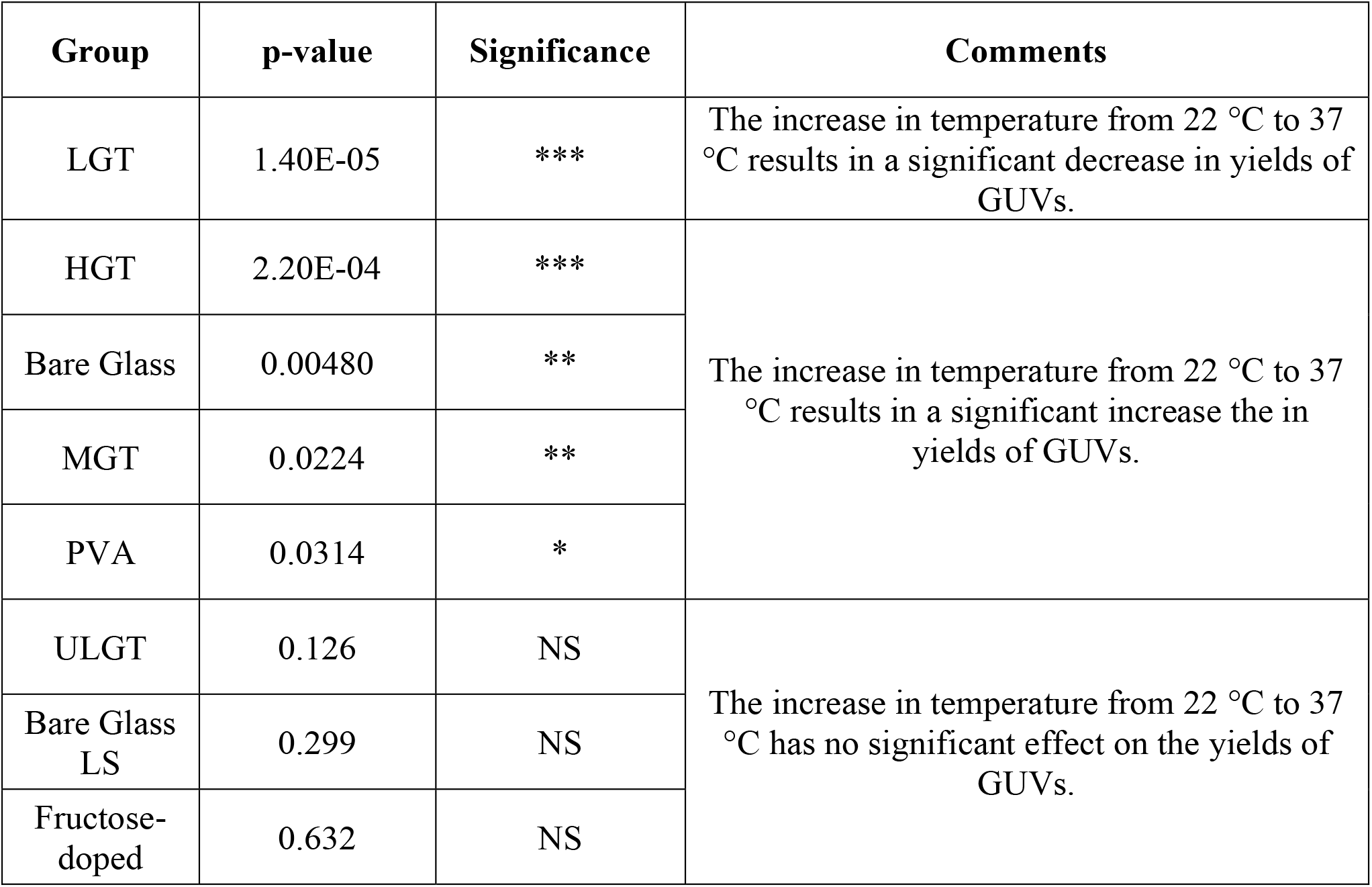
Table of p-values from Student’s t-tests of the molar yields of GUVs at 22 °C and 37 °C obtained from gentle hydration on bare glass in 100 mM sucrose (Bare Glass LS), gentle hydration on bare glass in PBS (Bare Glass), fructose-doped lipid in PBS, and hydration on polymers of PVA, ULGT agarose, LGT agarose, MGT agarose, and HGT agarose in PBS. All solutions contain 100 mM of sucrose. *= p < 0.05, ** = p < 0.01, *** = p < 0.001, NS = not significant.

**Table S5:**
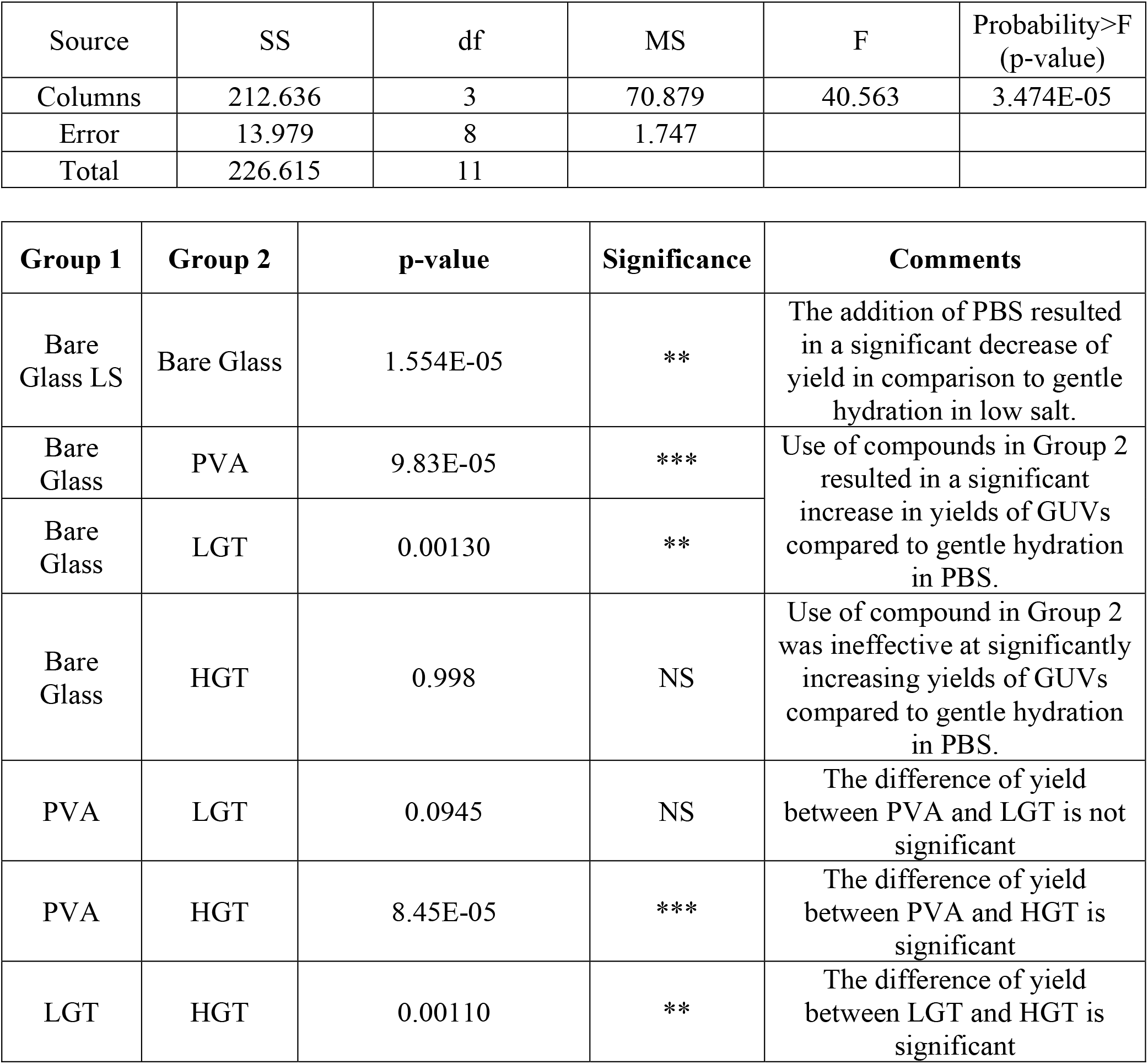
ANOVA table and table of p-values from post hoc Tukey’s HSD tests of the molar yields of GUVs composed of the ERGIC mixture grown at 22 °C obtained from gentle hydration on bare glass in 100 mM sucrose (Bare Glass LS), gentle hydration on bare glass in PBS (Bare Glass) and hydration on the polymer films of PVA, LGT agarose, and HGT agarose in PBS. All solutions contain 100 mM of sucrose. * = p < 0.05, ** = p < 0.01, *** = p < 0.001, NS = not significant.

**Table S6:**
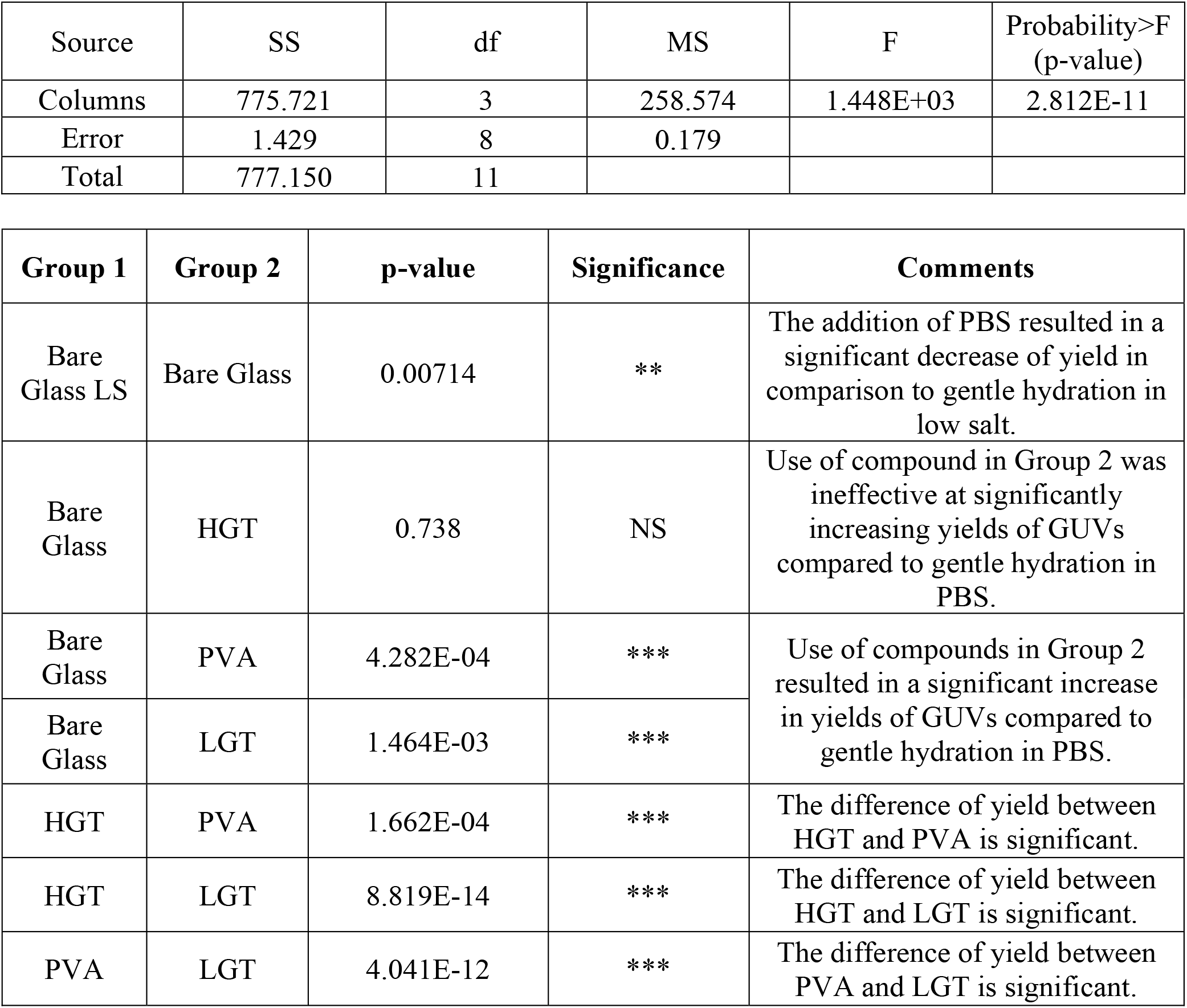
ANOVA table and table of p-values from post hoc Tukey’s HSD tests of the molar yields of GUVs composed of the MEL mixture grown at 22 °C obtained from gentle hydration on bare glass in 100 mM sucrose (Bare Glass LS), gentle hydration on bare glass in PBS (Bare Glass) and hydration on polymer films of PVA, LGT agarose, and HGT agarose in PBS. All solutions contain 100 mM of sucrose. * = p < 0.05, ** = p < 0.01, *** = p < 0.001, NS = not significant.

**Table S7:**
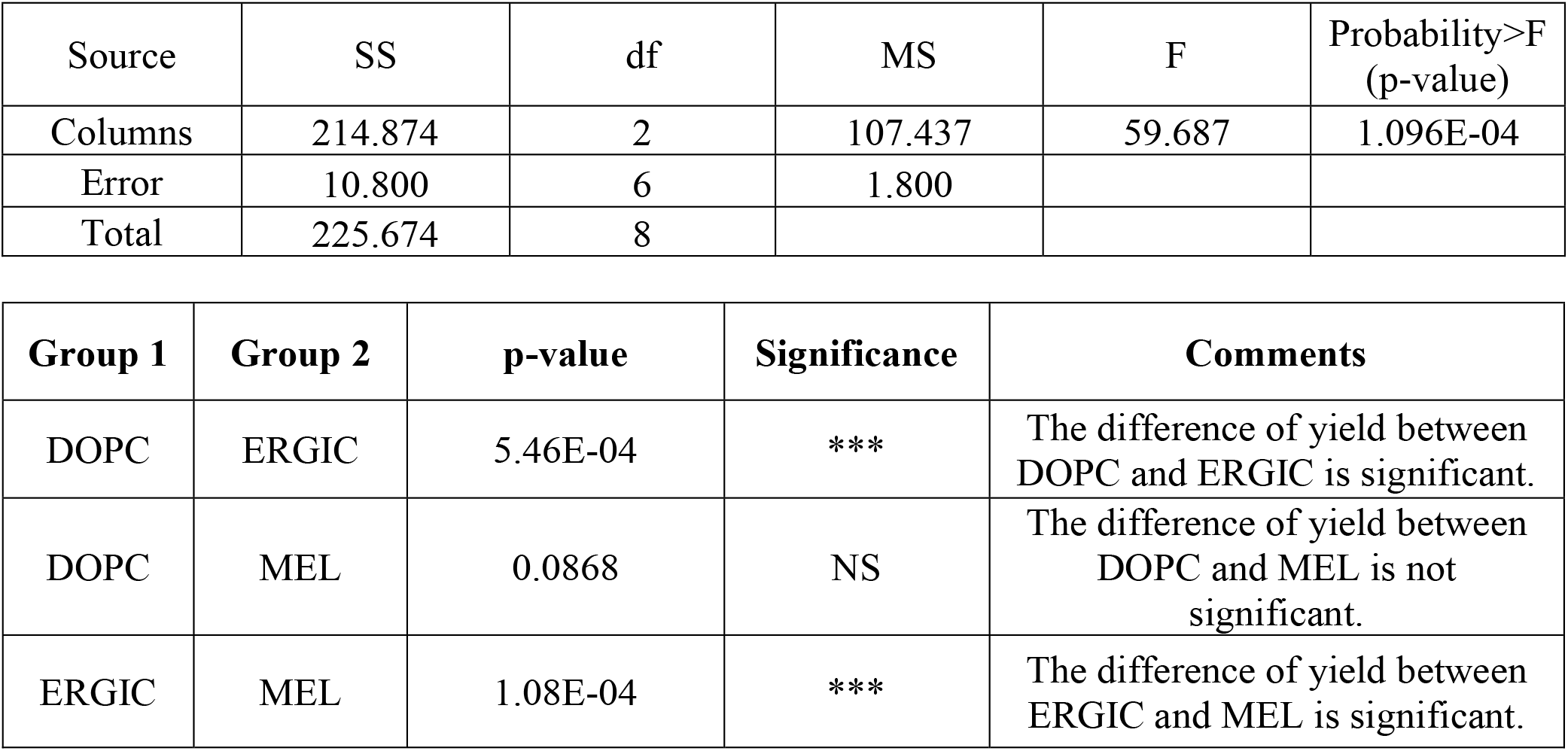
ANOVA table and table of p-values from post hoc Tukey’s HSD tests of the molar yields of GUVs composed of the MEL mixture, DOPC mixture and ERGIC mixture assembled at 22 °C on using LGT agarose as the assisting compound in PBS. All solutions contain 100 mM of sucrose. * = p < 0.05, ** = p < 0.01, *** = p < 0.001, NS = not significant.

**Table S8:**
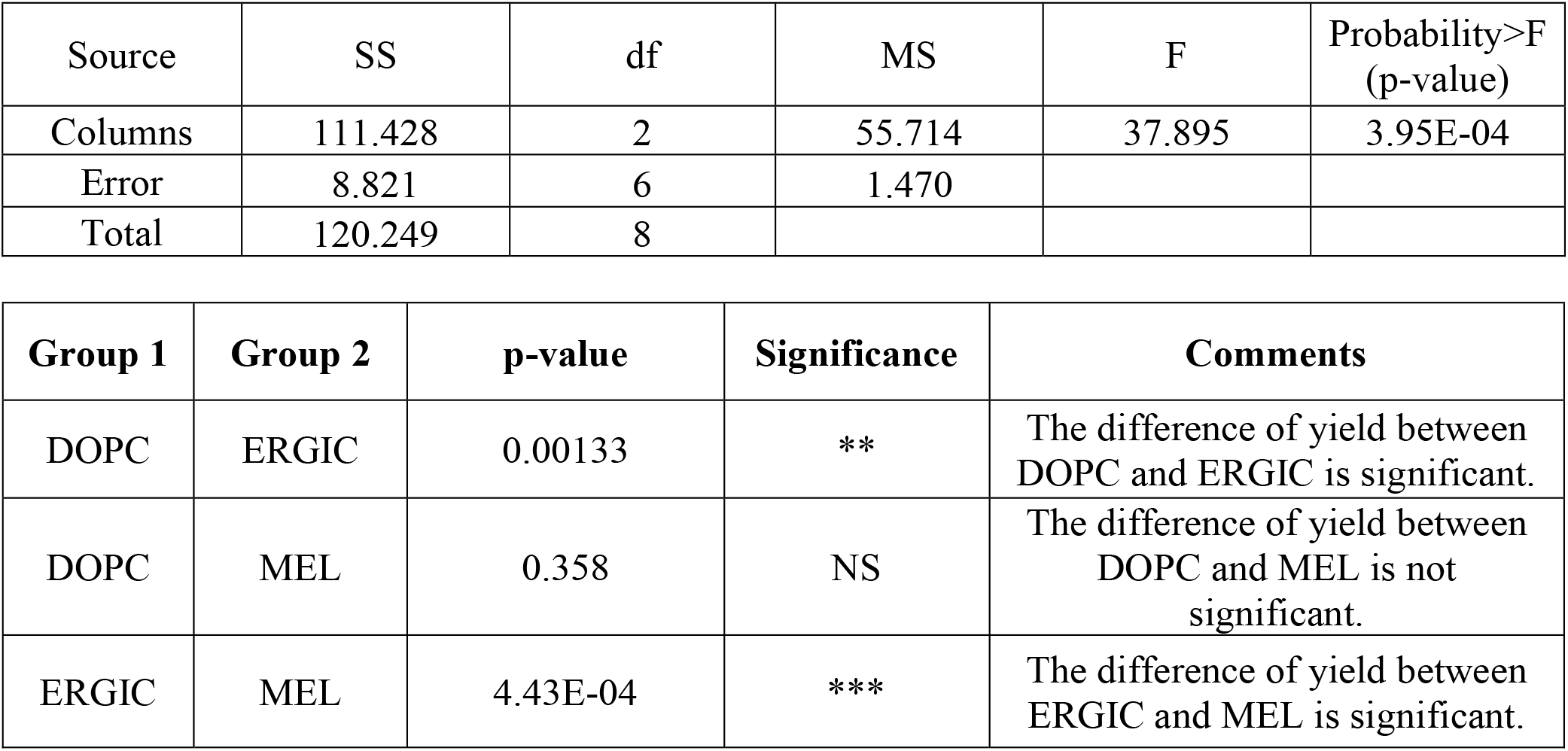
ANOVA table and table of p-values from post hoc Tukey’s HSD tests of the molar yields of GUVs composed of the MEL mixture, DOPC mixture and ERGIC mixture assembled at 22 °C on using PVA as the assisting compound in PBS. All solutions contain 100 mM of sucrose. * = p < 0.05, ** = p < 0.01, *** = p < 0.001, NS = not significant.

**Table S9:**
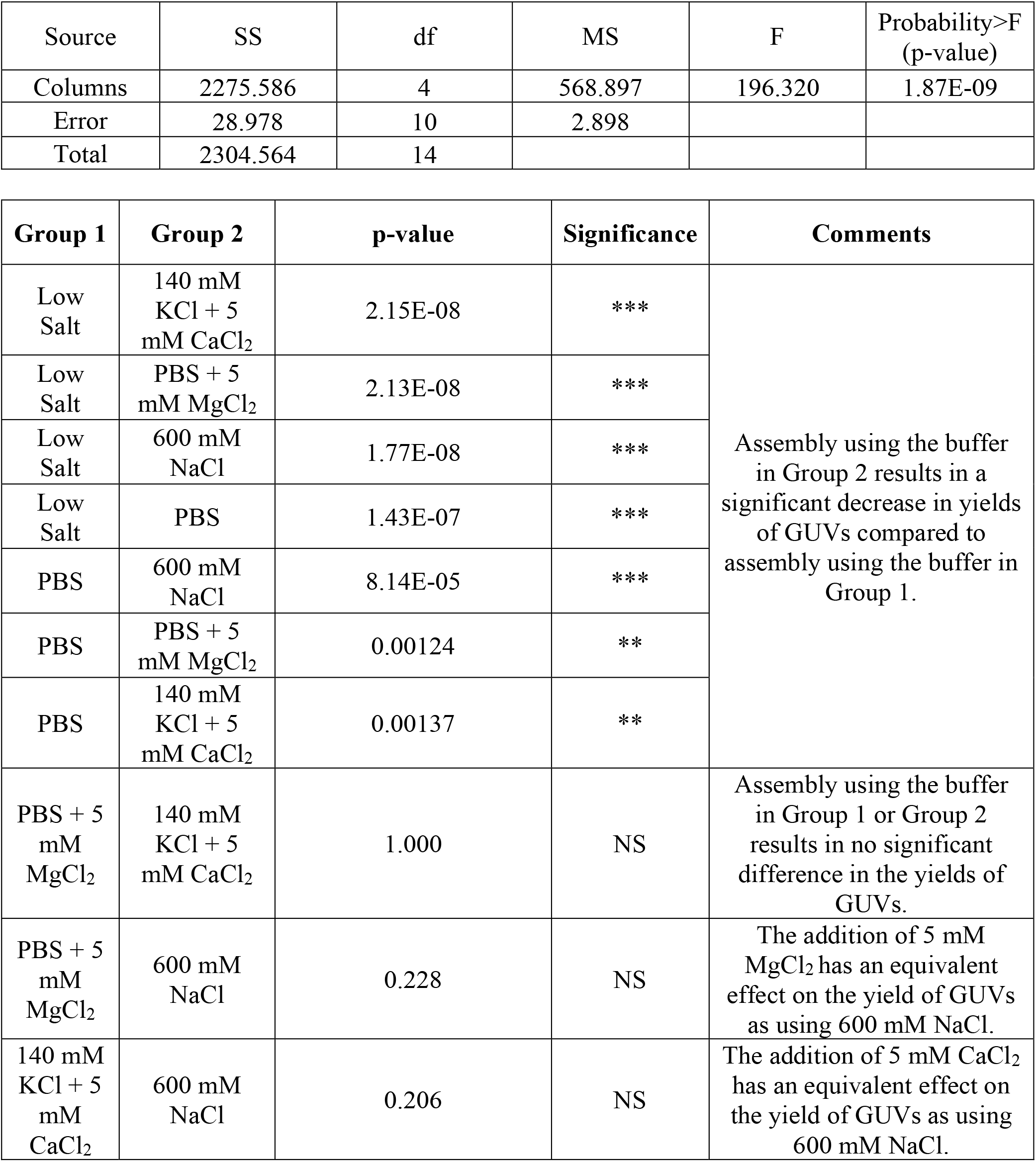
ANOVA table and table of p-values from post hoc Tukey’s HSD tests of the molar yields of GUVs assembled obtained using LGT agarose as the assisting compound in buffers of different ionic compositions. All solutions contain 100 mM of sucrose. * = p < 0.05, ** = p < 0.01, *** = p < 0.001, NS = not significant.

**Table S10.**
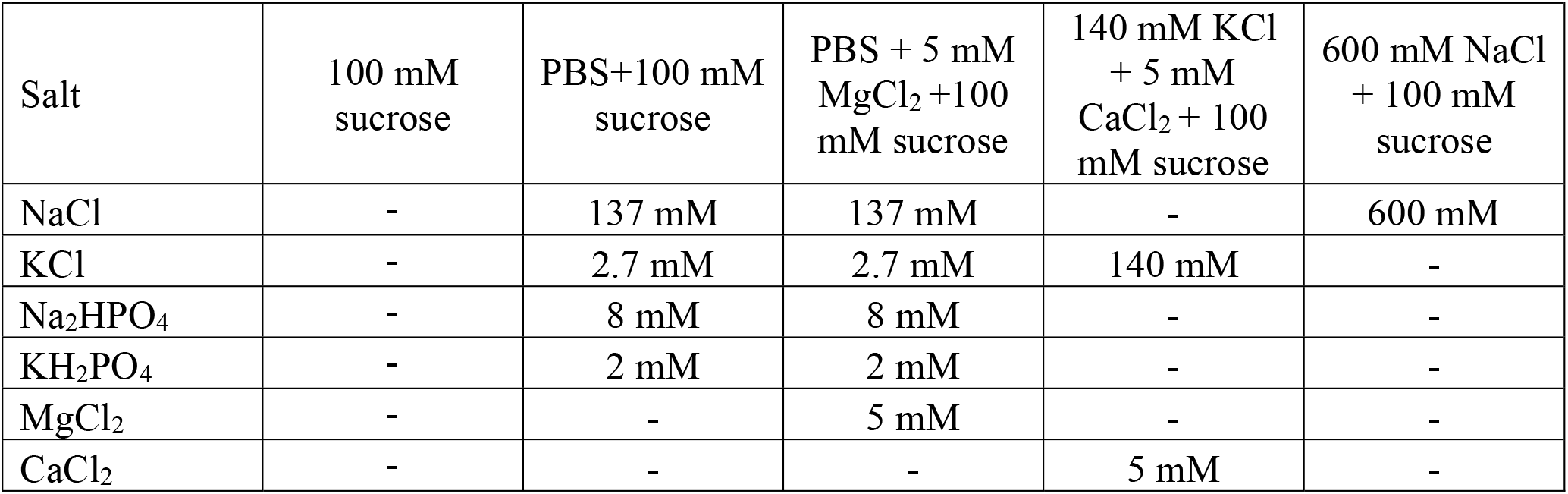
Table of the concentration of different salts in the hydration solutions used to gather the data in Figure 9.

**Table S11.**
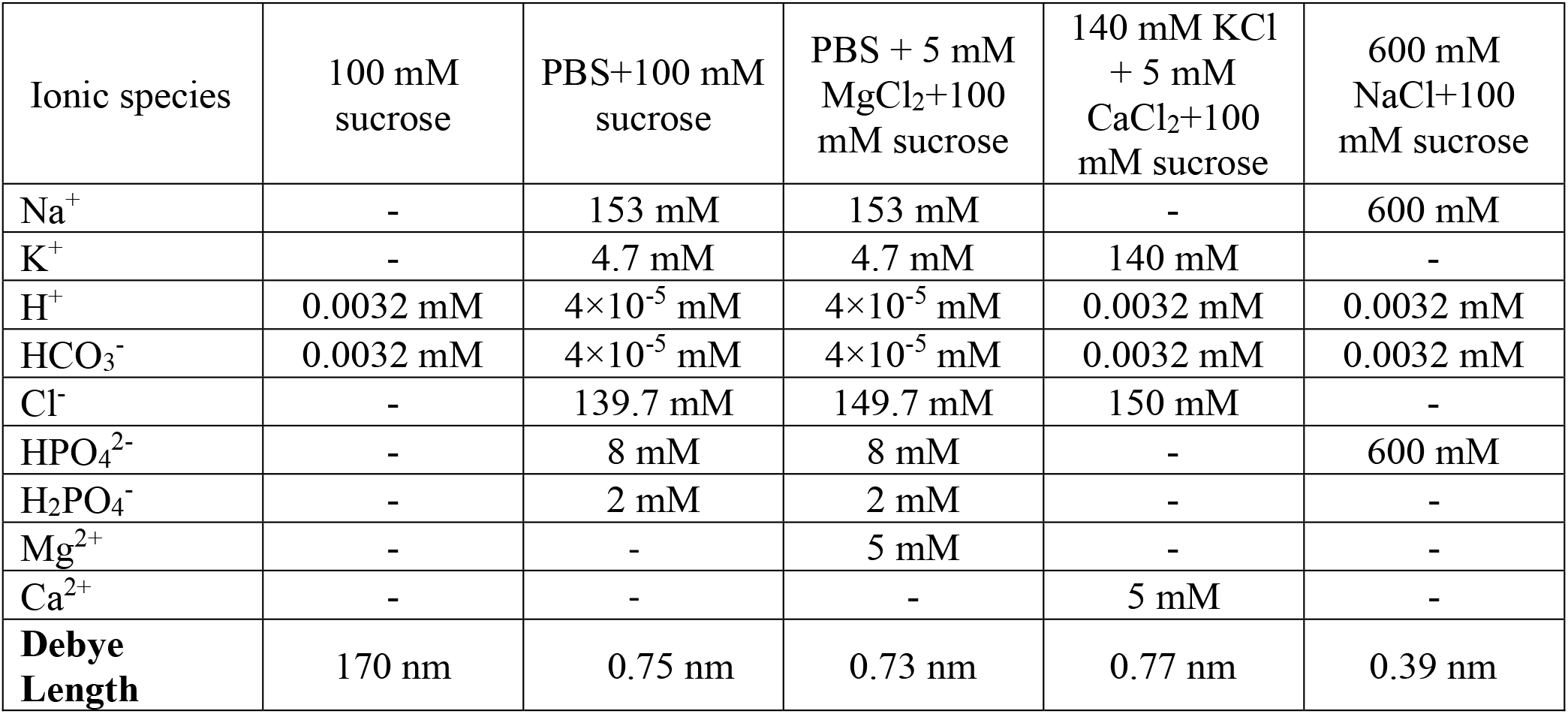
Table of the concentration of ions in the hydration solutions. We assume complete dissociation of all ions. The last row shows the calculated Debye screening length.

## Notes

### Competing Interest Statement

The authors have declared no competing interest.

### Summary of Updates

Added new lipid mixtures. Added estimates of budding energy and the amount of polymer in the interlamellar space.

