## SupportingInfo for "Osmotic pressure enables high yield assembly of giant vesicles in solutions of physiological ionic strengths"

### Supporting Text

**Determining the effect of dissolved polymers on the sedimentation of GUVs.** High concentrations of dissolved polymers in solution could in principle decrease the sedimentation velocity of the GUVs by increasing the viscosity of the solution. The polymers are present at low concentration in the sedimentation chamber.

To demonstrate, we calculate the maximum possible concentration of polymer in the solution containing the harvested GUVs (assuming all the polymers dissolve) using Equation S1. We assume all solutions have the density of water $\rho = 1\ \mathrm{g/mL}$.

$$w_{dissolved}^{max} = \frac{w_{ps} m_{ps}}{A_{coverslip}} \frac{A_{chamber}}{m_h} \tag{S1}$$

In this equation, $w_{dissolved}^{max}$ is the maximum concentration (w/w %) of dissolved polymer in the hydrating solution, $w_{ps}$ is the concentration of the polymer solution (1 w/w %) deposited on the glass coverslip, $m_{ps}$ is the mass of the polymer solution ($0.3\ \mathrm{mL} \approx 0.3\ \mathrm{g}$) deposited on the glass coverslip, $A_{coverslip}$ is the area of the coverslip (484 mm$^2$), $A_{chamber}$ is the area of the hydration chamber (113 mm$^2$), and $m_h$ is the mass of the hydrating buffer ($0.15\ \mathrm{mL} \approx 0.15\ \mathrm{g}$). $w_{dissolved}^{max} \approx 0.47\ \mathrm{w/w}\ \%$. Further, in the sedimentation chamber, the harvested vesicle suspension is diluted 30 times in the sedimentation buffer. The sedimentation buffer is devoid of polymers. Thus, the maximum concentration of dissolved polymers in the sedimentation chamber in the extreme case of complete dissolution is 0.016 w/w%.

The sedimentation chamber that we used has a height of 1000 µm. The vesicles are initially present in all locations in the chamber since they are well-mixed. To test for differences, after 3 hours, we image the solution at the imaging plane (5 µm above the coverslip) and the bulk at 100 µm and 900 µm above the imaging plane for GUVs composed of DOPC harvested from bare glass without any assisting compounds and when LGT agarose is used as the assisting compound. In both cases, we observe a large number of GUVs in the imaging plane and no GUVs in the locations in the bulk (Fig. S1). Our results confirm that the low concentration of polymers, $\ll 0.016$ wt %, in the solution in the sedimentation chamber does not have any measurable effect on the sedimentation behavior of GUVs.

**Calculation of molar yield from the literature.** Reference[1] reports that the total number of GUVs with diameters between 10 μm and 80 μm in a 20 μL aliquot from a harvested volume of 500 μL is 63 ± 14. The molar yield is defined as the mols of lipids in GUV membranes divided by the mols of lipids initially used[2]. The mols of lipid initially used in Reference 1 was $1\times 10^{-9}$ mols [1]. We estimate the maximum possible mols of lipids in the harvested vesicles reported in reference[1] using Equation S2.

$$moles\ of\ lipid = \frac{2\pi d^2 N_{GUV} V_h}{A_{hg} N_A V_{al}} \qquad \text{(S2)}$$

Here $N_{GUV}$ is the number of GUVs, taken as 77, $d_{GUV}$ is the diameter of the GUVs, taken to be 80 μm, $A_{hg}$ is the headgroup area of the lipid which for DOPC is $72.4 \times 10^{-4}$ μm$^2$, $N_A = 6.023 \times 10^{23}$ is Avogadro's number, $V_h$ is the harvested volume which is 500 μL, and $V_{al}$ is the aliquot counted which is 20 μL. Choosing the largest diameter of GUV reported and the upper range of GUVs counted in the experiment ensures that we are calculating the maximum possible mols of lipids harvested as GUVs from reference 1.

Dividing the moles of lipid in the harvested vesicles with the mols of lipids initially used gives a maximum molar yield of $6.0 \times 10^{-4}$ % for Reference 1. Notably, the diameters of GUVs quantified in Reference 1 were limited to 10 μm $\leq d <$ 100 μm [1]. When we similarly limit our measurements, we find that the yield of GUVs from the fructose-doped technique that we performed is $3.9 \times 10^{-2}$ %. This result is ~ 2 orders of magnitude greater than that calculated from reference 1. Nevertheless, these values show that the fructose-doped method produces a very low yield of GUVs in physiological salt solutions.

**Energy for forming spherical buds from flat surfaces.** The free energy of a budding membrane that is templated on a surface is modeled by Equation S3 [2].

$$\mathrm{E} = \int_{\boldsymbol{S}} d\boldsymbol{S} \left\{ \frac{\kappa_b}{2}(2H - H_0)^2 + \kappa_G G + \kappa_a \right\} + \int_V dV\, \Delta\mathrm{P} + \int_{\boldsymbol{S}} d\boldsymbol{S} \left\{ \xi\,(\boldsymbol{h}) \right\} \tag{S3}$$

In this equation, $\kappa_b$ is the bending modulus, $\kappa_G$ is the Gaussian bending modulus, $H = \frac{1}{2}(\kappa_1 + \kappa_2)$ is the mean curvature where $\kappa_1$ and $\kappa_2$ are the principal curvatures on the surface, $G = \kappa_1 \kappa_2$ is the Gaussian curvature, $H_0$ is the spontaneous curvature, $\kappa_a$ is the area expansion modulus, Δp is the difference in osmotic pressure, and $\xi(\boldsymbol{h})$ is the microscopic interaction potential normal to the surface of the membrane. The magnitude of $\xi(\boldsymbol{h})$ depends on the distance, $\boldsymbol{h}$, between the membrane and the surface[3]. These quantities are integrated as appropriate over the surface, $\boldsymbol{S}$, of the membrane, or the volume, $V$, that the membrane encloses.

Equation S3 can be simplified by noting that $H_0 = 0$ for a symmetric bilayer and that there is no change in the Gaussian curvature for spherical buds that remain attached to the surface[2]. We further simplify by replacing the microscopic interaction potential, $\xi(\boldsymbol{h})$, with an effective adhesion contact potential, $\xi$ [2]. We obtain Equation S4 by i) integrating Equation S3 to obtain the energy for State 1, the geometry of a flat disk of radius $R_d$, the energy for State 2, the geometry of a spherical bud of radius $R_b = \frac{R_d}{2}$, and ii) subtracting the energy of State 1 from State 2.

$$\Delta\mathrm{E} = 8\pi\kappa_B + 2\pi R_d \lambda - \pi {R_d}^2 \xi + \Delta P \Delta V \tag{S4}$$

The second term on the RHS of Equation S4 introduces a constraint for a section of the membrane to transition into a spherical bud at a constant area. If there is a lipid source, the membrane can transition without requiring breaks by recruiting lipids from the source[3]. In the absence of a lipid source, the membrane must form breaks, with an edge energy $\lambda$, to allow the lipids to reconfigure to form a spherical bud. In the main manuscript, for simplicity, we assume that the membrane has a source and does not break during budding, thus $\lambda = 0$. For gentle hydration on bare glass, $\Delta p = 0$. For polymer-coated surfaces $\Delta p$ depends on the concentration of dissolved polymer.

**Estimate of the osmotic pressure contribution of the dissolved polymer.** The behavior of the polymer and lipid is expected to be far from ideal due to the small length scales, the high concentrations, the large molecular weight of the polymer, and the time dependent process of lipid hydration and polymer dissolution. The polymers appear to interact with the lipid, which further changes the behavior. To make progress, we consider a simplified scaling argument. We assume ideal behavior and use characteristic length and energy scales to determine if the osmotic pressure exerted by the dissolving polymer at the concentrations that we use in our experiments can overcome an increase in adhesion between the membranes. Specifically, we estimate the concentration of dissolved polymer that is needed in the interlamellar space of the bilayer stacks so that the contribution of the osmotic pressure results in no change in the free energy between polymer-free budding in low salt solutions and polymer-assisted budding in salty solutions.

We assume that the effects of the salt and polymers on the edge energy, $\lambda$, and the bending rigidity, $\kappa_B$, to be negligible. The change in energy in the low-salt condition, $\Delta E_{LS}$ and salty condition, $\Delta E_{HS}$ is given by Equations S5 and S6.

$$\Delta \mathrm{E}_{\mathrm{LS}} = 8\pi\kappa_B + 2\pi R_d \lambda - \pi {R_d}^2 \xi_{LS} \quad \text{(S5)}$$

$$\Delta \mathrm{E}_{\mathrm{HS}} = 8\pi\kappa_B + 2\pi R_d \lambda - \pi {R_d}^2 \xi_{HS} + \Delta P \Delta V \quad \text{(S6)}$$

In these equations, $\xi_{LS}$ is the adhesion energy in the low salt solution, $\xi_{HS}$ is the adhesion energy in the salty solution, $\Delta P$ is the osmotic pressure difference, assumed to be constant, and $\varDelta V$ is the change in volume.

The change in volume from a disk-shaped bilayer on the surface with radius $R_d$ and interlamellar spacing height, $z$ to a spherical bud of radius $R_b$ is given by Equation S7a

$$\Delta V = \frac{1}{6}\pi R_d^3 - \pi R_d^2 z \qquad (S7a)$$

We used $R_b = \frac{R_d}{2}$ for the transition between a disk to a spherical bud at a constant surface area to express Equation S3a in terms of $R_d$.

For an ideal small molecule osmolyte at low concentrations, $\Delta P$ is related to the concentration of the osmolyte, $c$ by Equation S7b.

$$\Delta P = cRT \qquad (S7b)$$

In this equation, $R$ is the ideal gas constant, and $T$ is the temperature.

We obtain Equations S7c and S7d which satisfies the condition that there be no change in energy between assembly of GUVs in the low salt solution and the salty solution.

$$\Delta\Delta E = \Delta E_{HS} - \Delta E_{LS} = 0 \qquad (S7c)$$

$$\Delta P = -\frac{\pi R_d^2}{\varDelta V}(\xi_{HS} - \xi_{LS}) \qquad (S7d)$$

We substitute Equation S7a and S7b into equation S7d to get Equation S8. Equation S8 relates the concentration of the polymer in the interlamellar space that is needed to balance the increase in the adhesion energy in the salty solution compared to the low salt solution.

$$c = -\frac{6(\xi_{HS} - \xi_{LS})}{RT(R_d - 6z)} \quad \text{(S8)}$$

We take $\xi_{LS} = 1 \times 10^{-6}$ J m$^{-2}$ for DOPC membranes in low-salt solutions and $\xi_{HS} = 1 \times 10^{-4}$ J m$^{-2}$ for DOPC membranes in salty solutions[3]. We use $z = 4$ nm and $R_d = 1$ µm for a GUV bud 1 µm in diameter. We get $c \approx 0.2407\ \frac{\text{mol}}{\text{m}^3} = 0.24$ mM. The use of the ideal expression for osmotic pressure in the dilute limit does not change our conclusion that low amounts dissolved polymer is sufficient to exert an osmotic pressure that balances the adhesion energy in high salt solutions. For macromolecular polymers at moderate concentrations, the osmotic pressure is often expressed as $\Delta P = (c + A_2(Mc)^2)RT$, where $A_2$ is the second virial coefficient and $M$ is the molecular weight of the polymer[4]. The magnitude of the osmotic pressure, $\Delta P$, is thus expected to be higher for polymers compared to small molecule solutes for the same dissolved concentration, $c$. A lower amount of dissolved polymer than predicted by Equation S8 will be sufficient to balance the increased adhesion between membranes in salty solutions.

We next consider the expected concentration of dry polymer, $c_d$, in the interlamellar volume of the bilayer stack. The mass of polymer molecules per unit area of substrate (0.003 g of the polymer spread over a coverslip area of $4.84 \times 10^{-4}$ m$^2$) is 6.2 $\frac{\text{g}}{\text{m}^2}$. We assume that upon hydration the polymer and lipid form uniform stacks composed of 5 lipid bilayers with an interlamellar spacing of 4 nm. The concentration of dry polymer in the stack with a molecular weight, $M = 120{,}000$ is $c_d = 2583\ \frac{\text{mol}}{\text{m}^3} = 2.58$ M. Thus, in this ideal calculation, approximately 0.009 % of the polymer must dissolve to form GUV buds in salty solutions.

**Electrostatic interaction between membranes.** The adhesion of membranes is affected by the presence of ions in solutions[5–7]. Ions promote adhesion by screening repulsive electrostatic interactions. The effectiveness of screening can be estimated using the Debye screening length[3]. Smaller screening lengths reflect a shorter range in which electrostatic repulsion is felt, thus allowing attractive van der Waals interaction to dominate[3]. We use Equation S9[3] to calculate the Debye screening length, $\frac{1}{\kappa_D}$.

$$\kappa_D = \sqrt{\frac{N_a e^2}{\varepsilon_0 \varepsilon k_B T} \sum_i [C]_i z_i^2} \tag{S9}$$

Here $N_A$ is Avogadro's number, $e$ is the elementary charge, $[C]$ is the concentration of ionic species $i$, $z$ is the charge of ionic species $i$, $\varepsilon_0$ is the permittivity of free space, $\varepsilon$ is the dielectric constant, $k_B$ is the Boltzmann constant, and $T$ is the absolute temperature.

We report the ionic composition of the buffers used for the experiments in Figure 9 in Table S10. We report the ionic species and the calculated Debye screening lengths in Table S11. Note that the solutions consisting of 100 mM sucrose and 140 mM KCl + 5 mM $CaCl_2$ are unbuffered. Our measurements showed that they have a pH of 5.5. This value of pH is consistent with the formation of carbonic acid when ultrapure water is in equilibrium with atmospheric carbon dioxide[8]. The screening length drops from 170 nm in the solution consisting of 100 mM sucrose to 0.75 nm in the solution consisting of PBS + 100 mM sucrose. The screening length is further reduced to 0.39 nm in the solution consisting of 600 mM NaCl + 100 mM sucrose. The calculated Debye screening lengths of PBS + 5 mM $MgCl_2$ and 140 mM KCl + 5 mM $CaCl_2$ are 0.73 nm and 0.77 nm respectively. Although the values of the Debye lengths are similar to those of PBS, adhesion is significantly enhanced because mM concentration of divalent cations can bind and neutralize surface charges and serve as ionic bridges between charges[9,10].

**Supporting Figures**

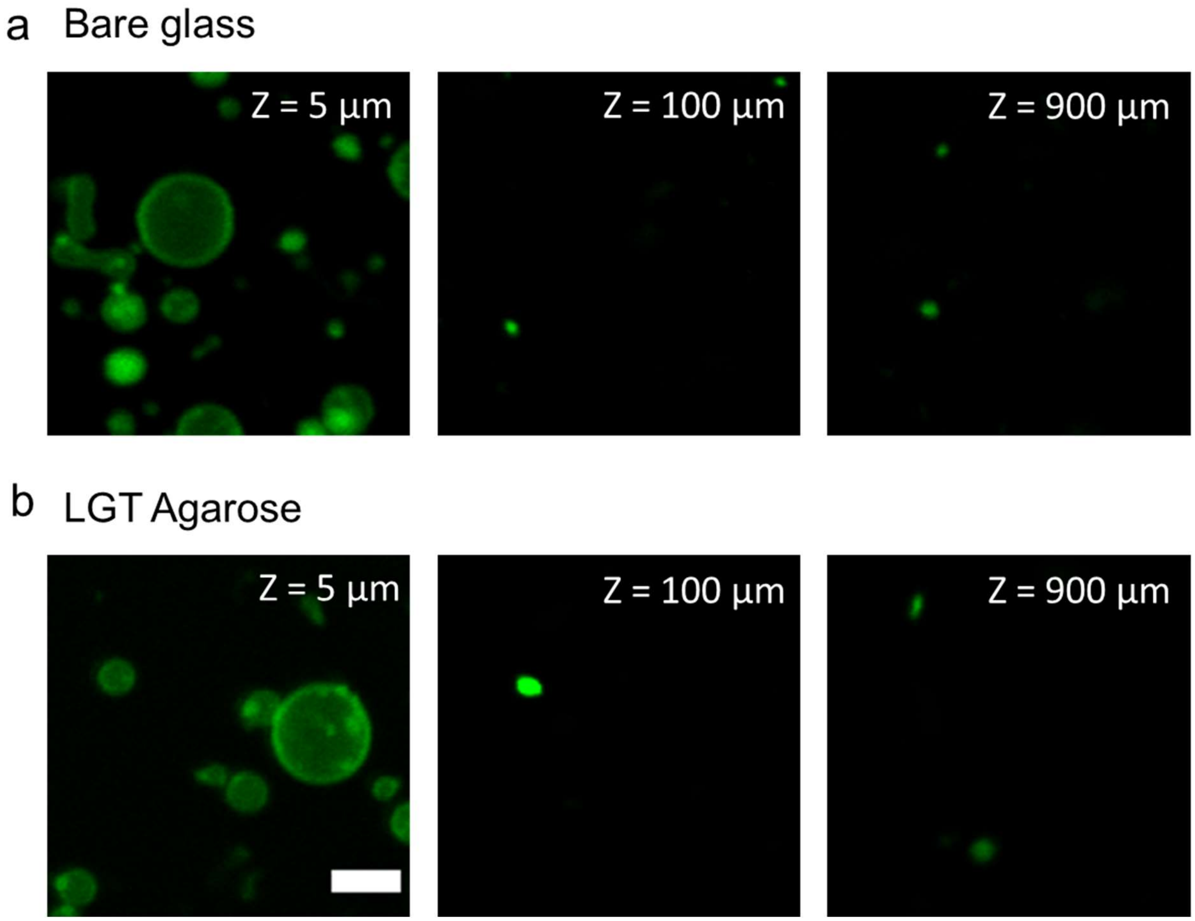


**Figure S1.** GUVs sediment in imaging chambers similarly in the presence and absence of polymers. Representative images at various *Z* planes of our imaging chamber after 3 hours of sedimentation. a) GUVs obtained from a bare glass surface without the use of assisting compounds. b) GUVs obtained when LGT agarose was used as the assisting compound. The Z position relative to the location of the surface of the bottom glass slide of the imaging chamber is indicated in the images. The scale bar is 10 µm.

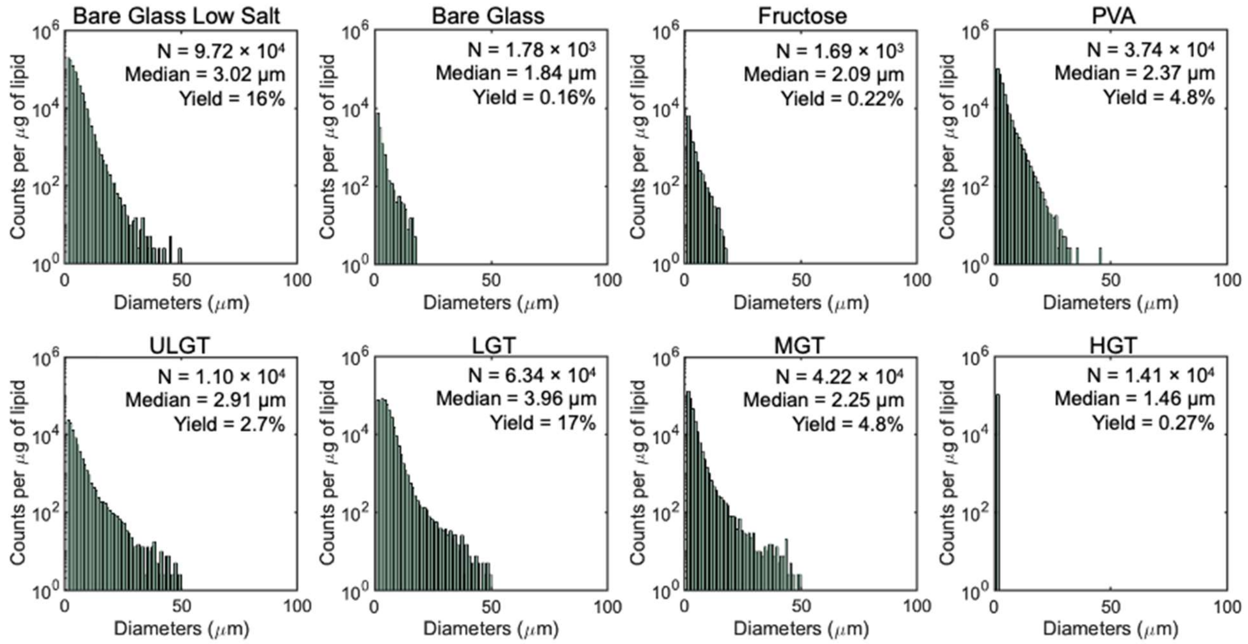


**Figure S2.** Histograms of GUV diameters of samples shown in Figure 2. Each histogram is the average of 3 independent repeats per sample. Note the logarithmic scale on the y-axis. Bin widths are 1 μm.

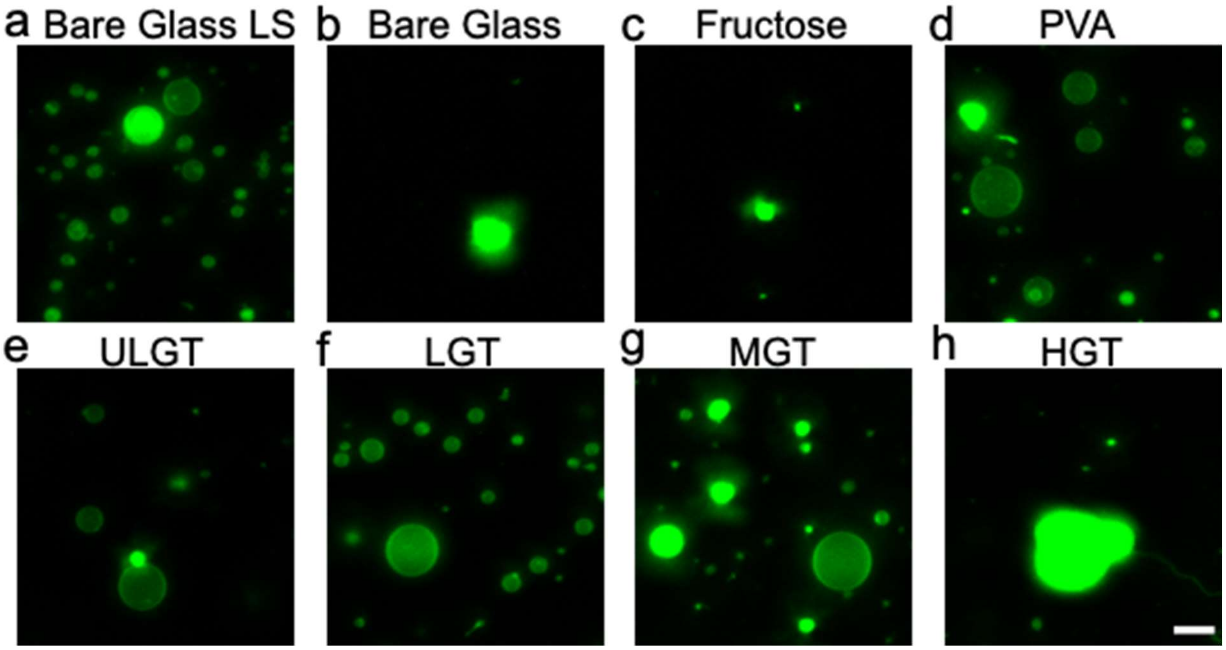


**Figure S3.** Representative images of the harvested objects for samples shown in Figure 2. Samples hydrated at 22 °C. Bright spots in all samples are lipid aggregates or MLVs. The scale bars are 15 μm.

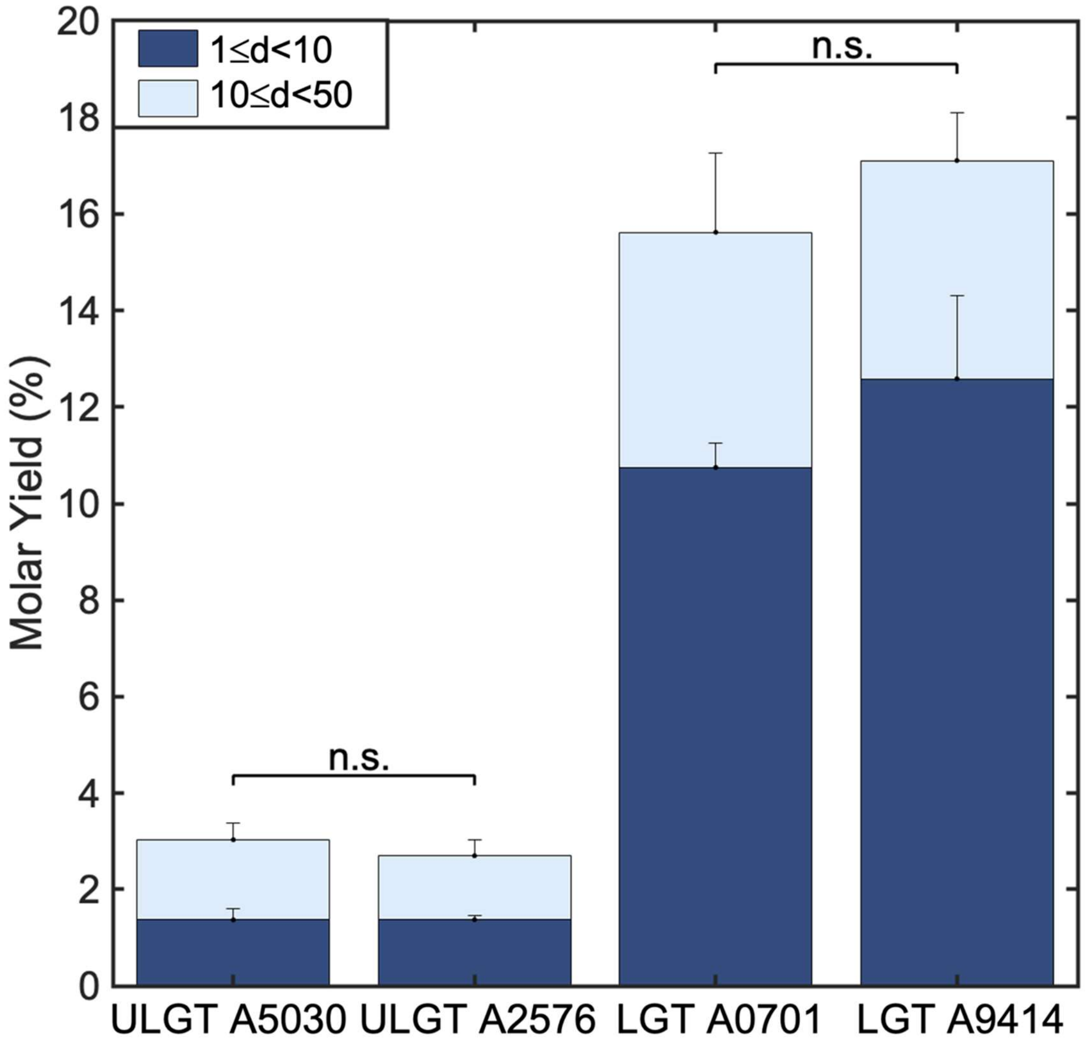


**Figure S4.** There is no significant difference in yields of GUVs obtained from ULGT and LGT agaroses with different catalog numbers (ultra-low gelling: A5030, A2576 and low gelling: A0701, A9414). Each bar is an average of 3 independent repeats per sample. Statistical significance determined by t-test. * = $p < 0.05$, ** = $p < 0.01$, *** = $p < 0.001$, ns = not significant. The data for A2576 and A9414 are from Figure 2.

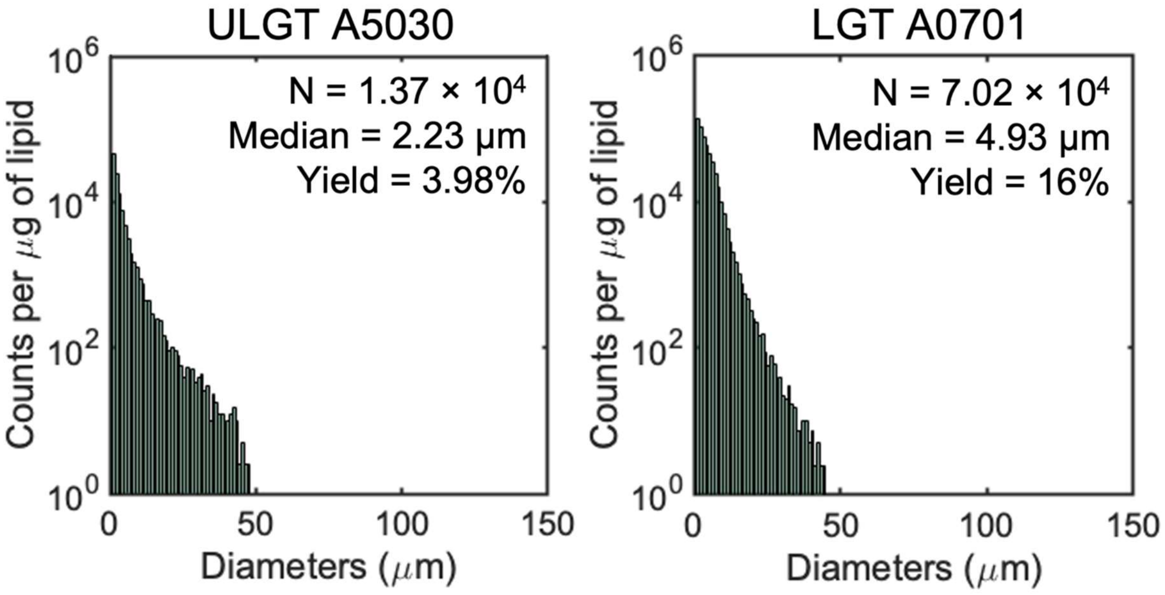


**Figure S5.** Histograms of GUV diameters of the samples shown in Figure S4. Each histogram is the average of 3 independent repeats per sample. Note the logarithmic scale on the y-axis. Bin widths are 1 μm.

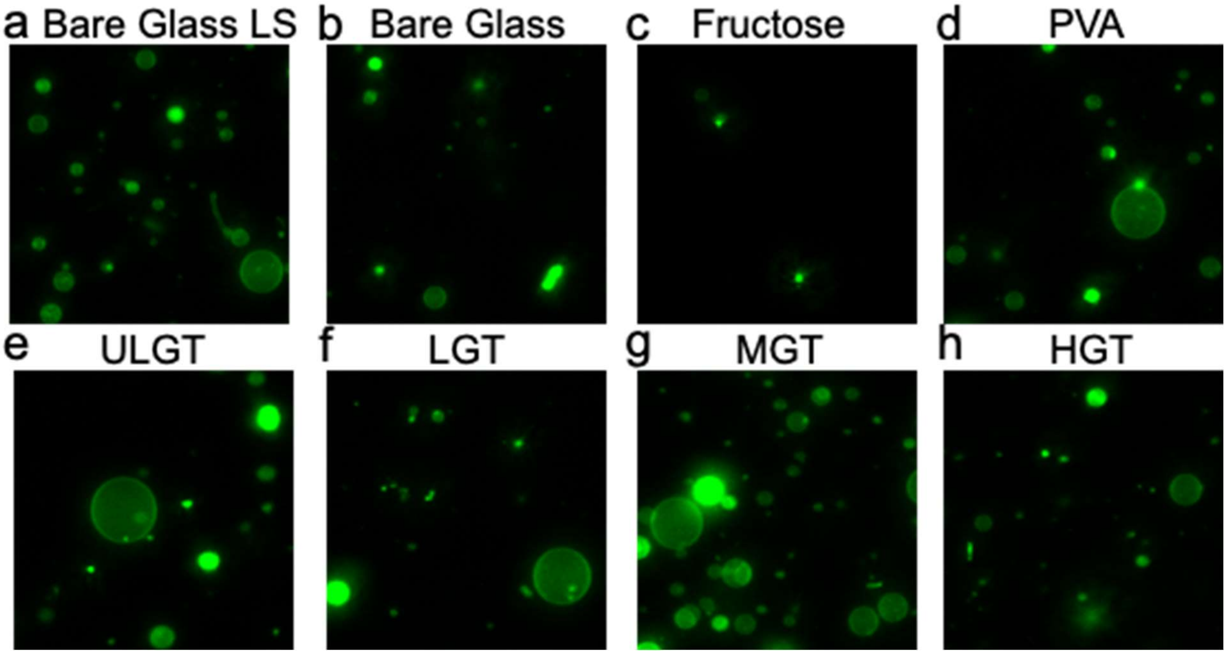


**Figure S6**. Representative images of the harvested objects for samples shown in Figure 3. Samples hydrated at 37 °C. Bright spots in all samples are lipid aggregates or MLVs. The scale bars are 15 μm.

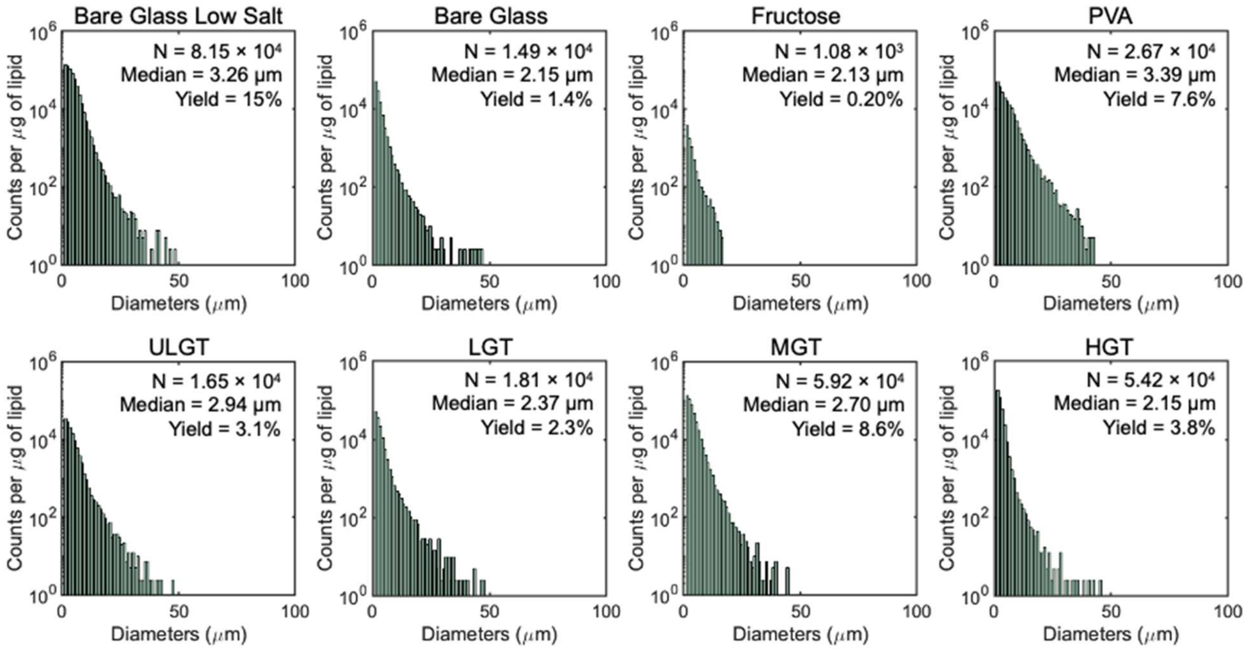


**Figure S7.** Histograms of GUV diameters of samples shown in Figure 3 for GUVs assembled at 37 °C. Each histogram is the average of 3 independent repeats per sample. Note the logarithmic scale on the y-axis. Bin widths are 1 μm.

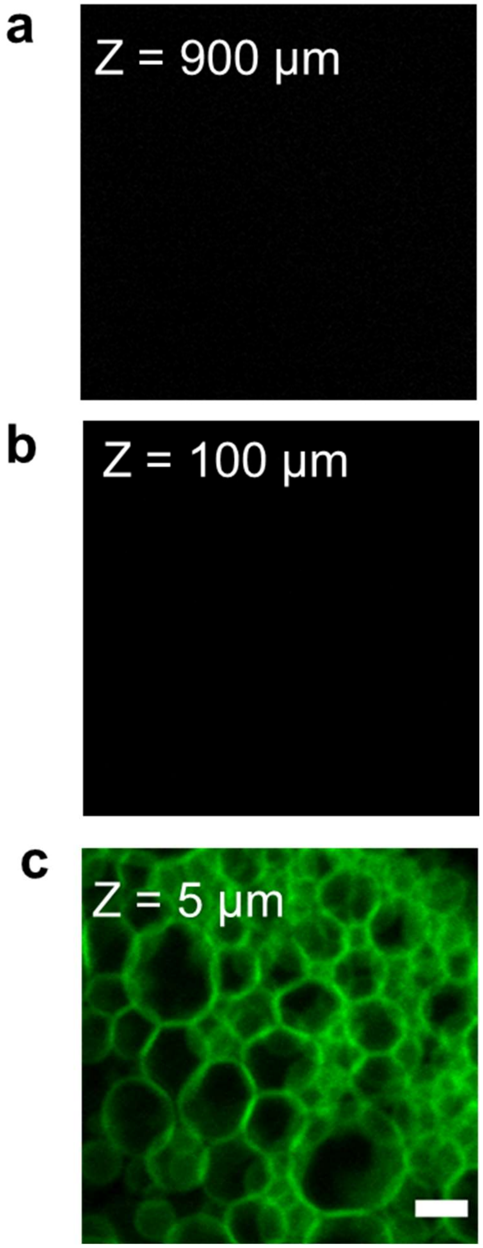


**Figure S8.** GUV buds remain attached to the surface in the absence of flow. Representative images after 2 hours of incubation at a) 900 µm and b) 100 µm above the LGT-agarose coated coverslip. Both regions had few to no floating structures. c) The surface of the agarose is covered with a high density of buds. The supporting glass coverslip is at Z = 0 µm. The scale bar is 10 µm.

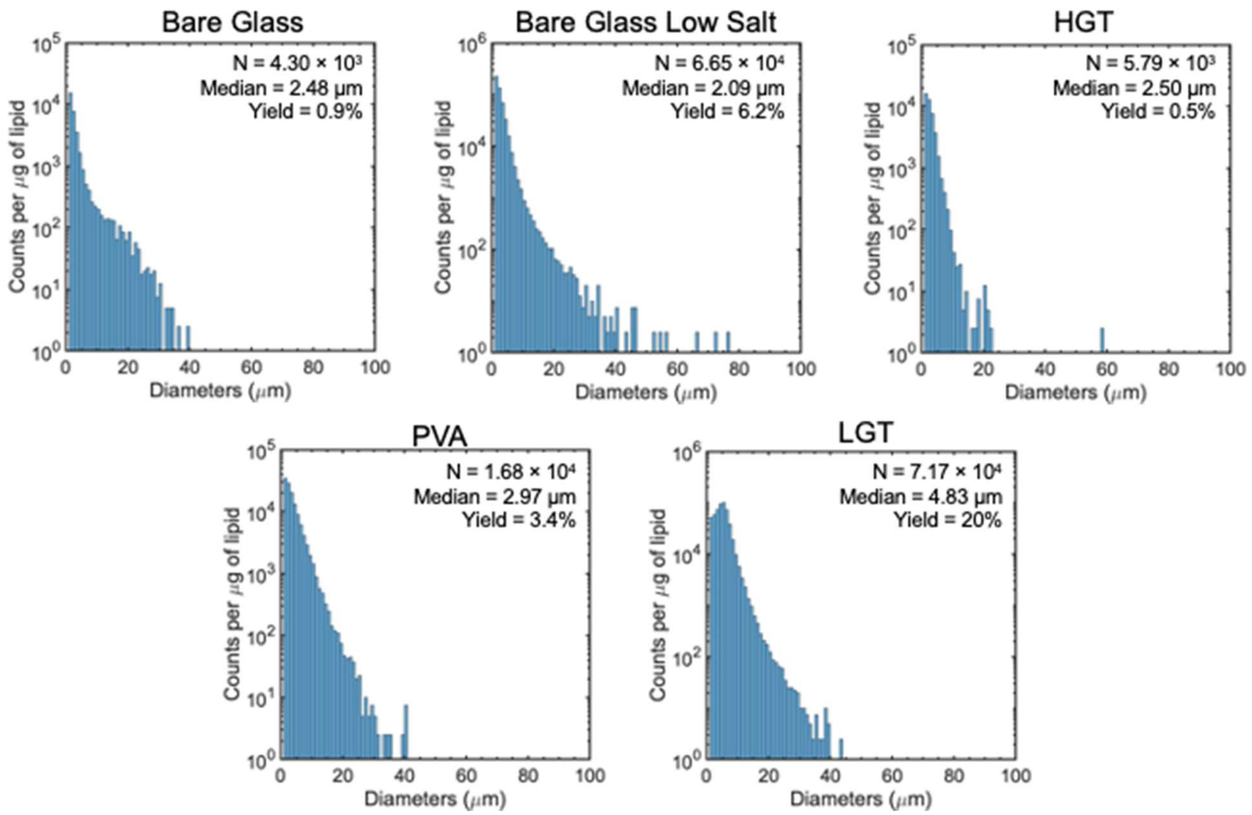


**Figure S9.** Histograms of GUV diameters of the samples shown in Figure S8a. The membranes of the GUVs are composed of a lipid mixture that minimally mimics the composition of the exoplasmic leaflet of the mammalian cellular membrane (mammalian exoplasmic leaflet (MEL)). Each histogram is the average of 3 independent repeats per sample. Note the logarithmic scale on the y-axis. Bin widths are 1 µm.

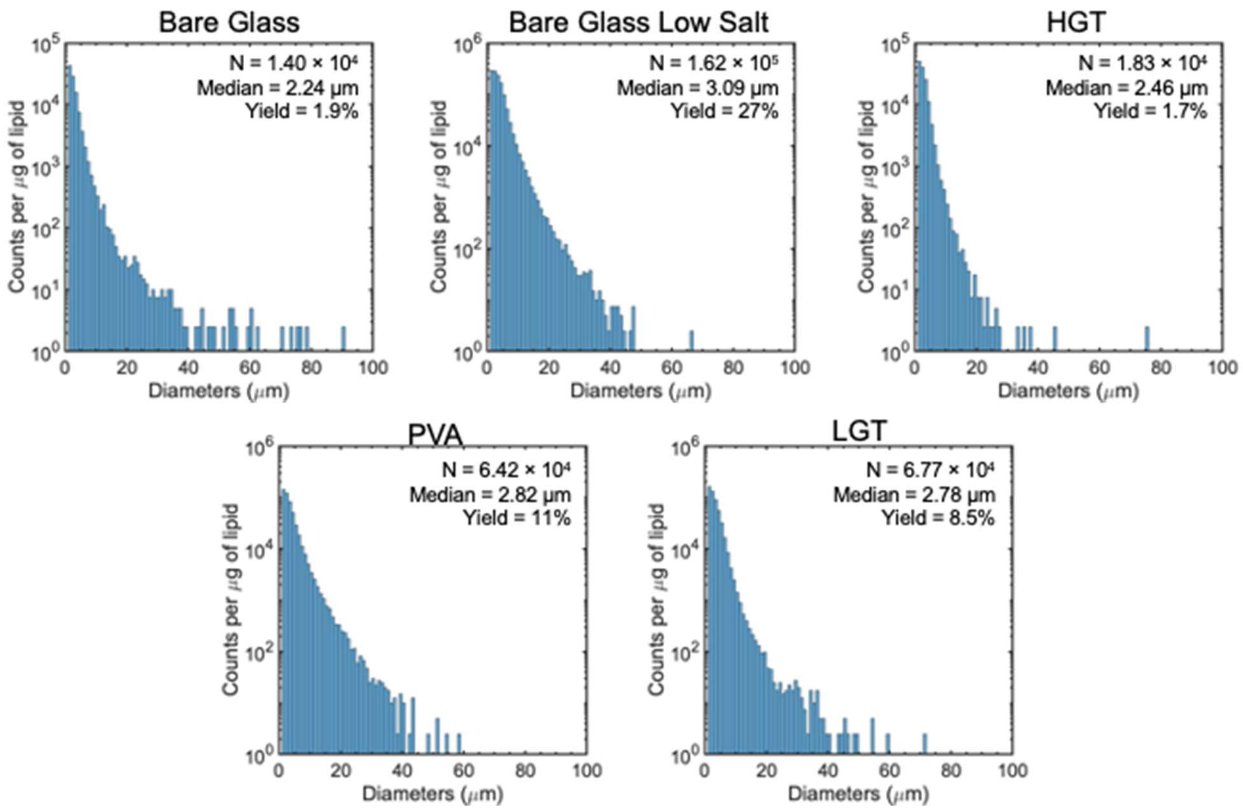


**Figure S10.** Histograms of GUV diameters of the samples shown in Figure S8b. The membranes of the GUVs are composed of a lipid mixture that minimally mimics the composition of the endoplasmic-reticulum-Golgi intermediate compartment (ERGIC) membrane. Each histogram is the average of 3 independent repeats per sample. Note the logarithmic scale on the y-axis. Bin widths are 1 µm.

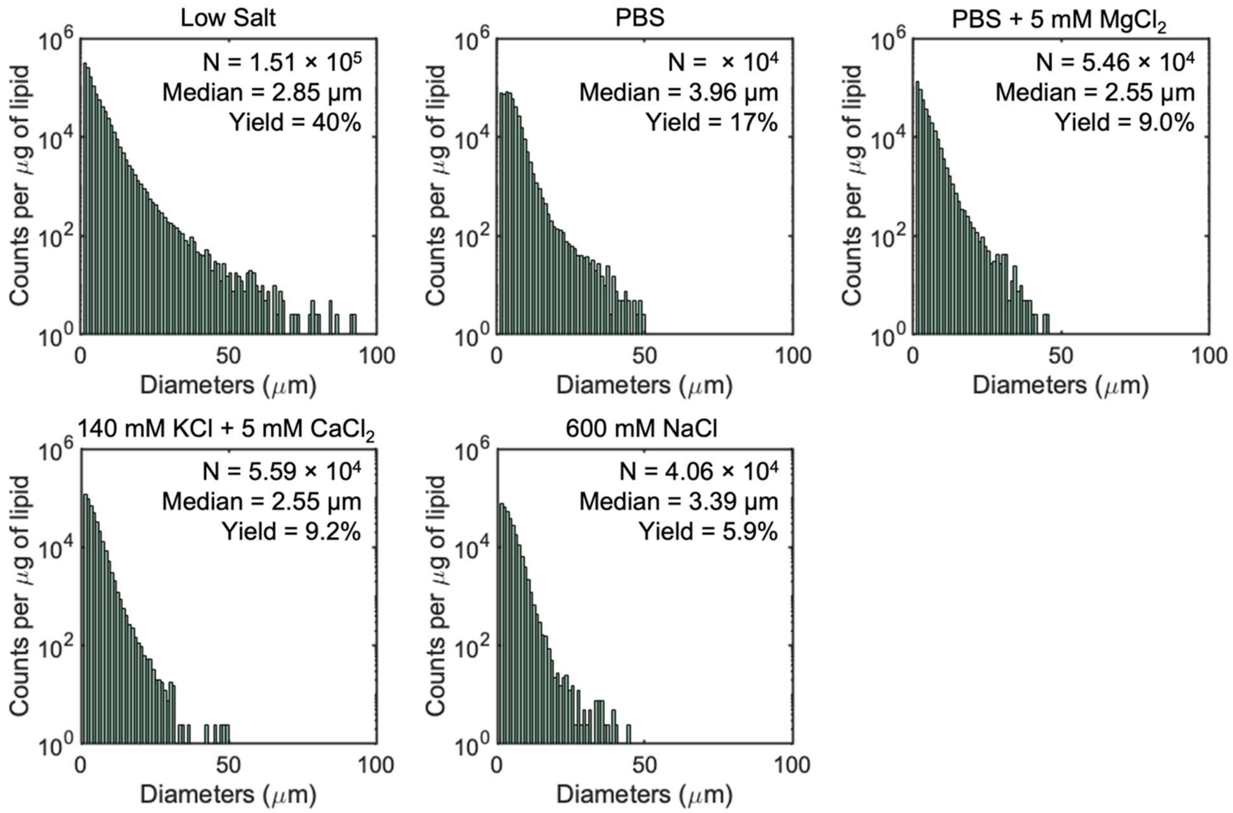


**Figure S11.** Histograms of GUV diameters of samples shown in Figure 9. The membranes of the GUVs are all composed of DOPC and the assisting compound is low gelling temperature (LGT) agarose. Each histogram is the average of 3 independent repeats per sample. Note the logarithmic scale on the y-axis. Bin widths are 1 µm.

**Supporting Tables**

| Compound | Gel Point (°C) | Melting Point (°C) |
|---|---|---|
| **ULGT Agarose (Type IX-A, Catalog number: A2576)** | ≤17 at 1.5% | ≤60 |
| **LGT Agarose (Catalog Number: A9414)** | ~30 | ~65 |
| **MGT Agarose (Type II-A, Catalog Number: A9918)** | 36±1.5 at 1.5% | 87±1.5 |
| **HGT Agarose (Type VI-A, Catalog Number: A7174)** | 41±1.5 at 1.5% | 95±1.5 |
| **ULGT Agarose (Type IX, Catalog Number: A5030)** | 8-17 at 0.8% | ≤60 |
| **LGT Agarose (Type VII-A, Catalog Number: A0701)** | 26±2.0 at 1.5% | ≤65.5 |
| **Polyvinyl alcohol** | 85* | N/A |
| **Fructose** | N/A | N/A |

**Table S1.** Melting and gelling temperatures of the compounds tested. *Glass transition temperature. Values for agarose are from [11], values for PVA is from [12].

| Source | SS | df | MS | F | Probability>F (p-value) |
|---|---|---|---|---|---|
| Columns | 651.351 | 6 | 108.558 | 149.654 | 7.04E-12 |
| Error | 10.156 | 14 | 0.725 | | |
| Total | 661.506 | 20 | | | |

| **Group 1** | **Group 2** | **p-value** | **Significance** | **Comments** |
|---|---|---|---|---|
| Bare Glass LS | Bare Glass | 6.732E-08 | *** | The addition of PBS resulted in a significant decrease of yield in comparison to gentle hydration in low salt. |
| Bare Glass LS | LGT | 0.347 | NS | The use of LGT agarose allows for assembly of GUVs in PBS with a molar yield not significantly different from bare glass in low salt. |
| Bare Glass | Fructose-doped | > 0.999 | NS | Use of compounds in Group 2 was ineffective at increasing the yields of GUVs compared to gentle hydration in PBS. |
| Bare Glass | HGT | > 0.999 | NS | |
| Bare Glass | ULGT | 0.0326 | * | Use of compounds in Group 2 was effective at increasing the yields of GUVs compared to gentle hydration in PBS. |
| Bare Glass | PVA | 1.52E-04 | *** | |
| Bare Glass | MGT | 1.59E-04 | *** | |
| Bare Glass | LGT | 3.76E-08 | *** | |
| Fructose-doped | HGT | > 0.999 | NS | Use of compounds in Group 2 was not effective at increasing the yields of GUVs compared to gentle hydration using fructose-doped lipid in PBS. |
| Fructose-doped | ULGT | 0.0388 | * | Use of compounds in Group 2 was effective at increasing the yields of GUVs compared to gentle hydration using fructose-doped lipid in PBS. |
| Fructose-doped | PVA | 1.77E-04 | *** | |
| Fructose-doped | MGT | 1.85E-04 | *** | |
| Fructose-doped | LGT | 3.76E-08 | ** | |

| | | | | |
|---|---|---|---|---|
| HGT | ULGT | 0.0445 | * | The difference in yield between HGT and compounds in group 2 is significant. |
| HGT | PVA | 2.00E-04 | *** | |
| HGT | MGT | 2.10E-04 | *** | |
| HGT | LGT | 3.76E-08 | *** | |
| ULGT | PVA | 0.0895 | NS | The difference in yield between ULGT and compounds in group 2 not significant. |
| ULGT | MGT | 0.0940 | NS | |
| ULGT | LGT | 3.77E-08 | *** | The difference in yield between ULGT and LGT is not significant. |
| PVA | MGT | 0.999 | NS | The difference in yield between PVA and MGT is not significant. |
| PVA | LGT | 3.86E-08 | *** | The difference in yield between PVA and LGT is significant. |
| MGT | LGT | 3.85E-08 | *** | The difference in yield between MGT and LGT is significant. |

**Table S2:** ANOVA table and table of p-values from post hoc Tukey's HSD tests of the molar yields of GUVs assembled at 22 °C obtained from gentle hydration on bare glass in PBS (Bare Glass), fructose-doped lipid in PBS, and hydration on polymer films of PVA, ULGT agarose, LGT agarose, MGT agarose, and HGT agarose in PBS. Comparison of gentle hydration on bare glass in low salt (Bare Glass LS) and gentle hydration on bare glass in PBS (Bare Glass) was conducted separately from the ANOVA via Student's t-test. All solutions contain 100 mM of sucrose. * = $p < 0.05$, ** = $p < 0.01$, *** = $p < 0.001$, NS = not significant.

| Source | SS | df | MS | F | Probability>F (p-value) |
|---|---|---|---|---|---|
| Columns | 176.662 | 6 | 29.444 | 111.642 | 5.23E-11 |
| Error | 3.692 | 14 | 0.264 | | |
| Total | 180.354 | 20 | | | |

| **Group 1** | **Group 2** | **p-value** | **Significance** | **Comments** |
|---|---|---|---|---|
| Bare Glass LS | Bare Glass | 1.34E-04 | *** | The addition of PBS resulted in a significant decrease of yield in comparison to gentle hydration in low salt. |
| Bare Glass LS | LGT | 1.58E-04 | *** | The use of LGT agarose at 37 °C results in a yield significantly lower than bare glass in low salt. |
| Bare Glass | Fructose-doped | 0.129 | NS | Use of compounds in Group 2 was ineffective at increasing the yields of GUVs compared to gentle hydration in PBS. |
| Bare Glass | LGT | 0.436 | NS | |
| Bare Glass | HGT | 8.84E-04 | *** | Use of compounds in Group 2 was effective at increasing the yields of GUVs compared to gentle hydration in PBS. |
| Bare Glass | ULGT | 0.0160 | * | |
| Bare Glass | PVA | 4.87E-08 | *** | |
| Bare Glass | MGT | 3.90E-08 | *** | |
| Fructose-doped | HGT | 1.06E-05 | *** | Use of compounds in Group 2 was effective at increasing the yields of GUVs compared to gentle hydration using fructose-doped lipid in PBS. |
| Fructose-doped | ULGT | 1.15E-04 | *** | |
| Fructose-doped | PVA | 3.86E-08 | *** | |
| Fructose-doped | MGT | 3.78E-08 | *** | |
| Fructose-doped | LGT | 3.39E-03 | ** | |
| HGT | ULGT | 0.672 | NS | The difference in yield between HGT and ULGT is not significant. |
| HGT | PVA | 5.28E-06 | *** | |

| | | | | |
|---|---|---|---|---|
| HGT | MGT | 3.11E-07 | *** | The difference in yield between HGT and compounds in group 2 is significant. |
| HGT | LGT | 0.0338 | * | |
| ULGT | PVA | 7.36E-07 | *** | The difference in yield between ULGT and compounds in group 2 is significant. |
| ULGT | MGT | 8.77E-08 | *** | |
| ULGT | LGT | 0.449 | NS | The difference in yield between ULGT and LGT is not significant. |
| PVA | MGT | 0.249 | NS | The difference in yield between PVA and MGT is not significant. |
| PVA | LGT | 1.15E-07 | *** | The difference in yield between PVA and LGT is significant. |
| MGT | LGT | 4.53E-08 | *** | The difference in yield between MGT and LGT is significant. |

**Table S3:** ANOVA table and table of p-values from post hoc Tukey's HSD tests of the molar yields of GUVs grown at 37 °C obtained from gentle hydration on bare glass in PBS (Bare Glass), fructose-doped lipid in PBS, and hydration on polymer films of PVA, ULGT agarose, LGT agarose, MGT agarose, and HGT agarose in PBS. Comparison of gentle hydration on bare glass in low salt (Bare Glass LS) and gentle hydration on bare glass in PBS (Bare Glass) was conducted separately from the ANOVA via Student's t-test. All solutions contain 100 mM of sucrose. * = $p < 0.05$, ** = $p < 0.01$, *** = $p < 0.001$, NS = not significant.

<table>
<tr><th>Group</th><th>p-value</th><th>Significance</th><th>Comments</th></tr>
<tr><td>LGT</td><td>1.40E-05</td><td>***</td><td>The increase in temperature from 22 °C to 37 °C results in a significant decrease in yields of GUVs.</td></tr>
<tr><td>HGT</td><td>2.20E-04</td><td>***</td><td rowspan="4">The increase in temperature from 22 °C to 37 °C results in a significant increase the in yields of GUVs.</td></tr>
<tr><td>Bare Glass</td><td>0.00480</td><td>**</td></tr>
<tr><td>MGT</td><td>0.0224</td><td>**</td></tr>
<tr><td>PVA</td><td>0.0314</td><td>*</td></tr>
<tr><td>ULGT</td><td>0.126</td><td>NS</td><td rowspan="3">The increase in temperature from 22 °C to 37 °C has no significant effect on the yields of GUVs.</td></tr>
<tr><td>Bare Glass LS</td><td>0.299</td><td>NS</td></tr>
<tr><td>Fructose-doped</td><td>0.632</td><td>NS</td></tr>
</table>

**Table S4:** Table of p-values from Student's t-tests of the molar yields of GUVs at 22 °C and 37 °C obtained from gentle hydration on bare glass in 100 mM sucrose (Bare Glass LS), gentle hydration on bare glass in PBS (Bare Glass), fructose-doped lipid in PBS, and hydration on polymers of PVA, ULGT agarose, LGT agarose, MGT agarose, and HGT agarose in PBS. All solutions contain 100 mM of sucrose. *= $p < 0.05$, ** = $p < 0.01$, *** = $p < 0.001$, NS = not significant.

| Source | SS | df | MS | F | Probability>F (p-value) |
|---|---|---|---|---|---|
| Columns | 212.636 | 3 | 70.879 | 40.563 | 3.474E-05 |
| Error | 13.979 | 8 | 1.747 | | |
| Total | 226.615 | 11 | | | |

| **Group 1** | **Group 2** | **p-value** | **Significance** | **Comments** |
|---|---|---|---|---|
| Bare Glass LS | Bare Glass | 1.554E-05 | ** | The addition of PBS resulted in a significant decrease of yield in comparison to gentle hydration in low salt. |
| Bare Glass | PVA | 9.83E-05 | *** | Use of compounds in Group 2 resulted in a significant increase in yields of GUVs compared to gentle hydration in PBS. |
| Bare Glass | LGT | 0.00130 | ** | |
| Bare Glass | HGT | 0.998 | NS | Use of compound in Group 2 was ineffective at significantly increasing yields of GUVs compared to gentle hydration in PBS. |
| PVA | LGT | 0.0945 | NS | The difference of yield between PVA and LGT is not significant |
| PVA | HGT | 8.45E-05 | *** | The difference of yield between PVA and HGT is significant |
| LGT | HGT | 0.00110 | ** | The difference of yield between LGT and HGT is significant |

**Table S5:** ANOVA table and table of p-values from post hoc Tukey's HSD tests of the molar yields of GUVs composed of the ERGIC mixture grown at 22 °C obtained from gentle hydration on bare glass in 100 mM sucrose (Bare Glass LS), gentle hydration on bare glass in PBS (Bare Glass) and hydration on the polymer films of PVA, LGT agarose, and HGT agarose in PBS. All solutions contain 100 mM of sucrose. * = $p < 0.05$, ** = $p < 0.01$, *** = $p < 0.001$, NS = not significant.

| Source | SS | df | MS | F | Probability>F (p-value) |
|---|---|---|---|---|---|
| Columns | 775.721 | 3 | 258.574 | 1.448E+03 | 2.812E-11 |
| Error | 1.429 | 8 | 0.179 | | |
| Total | 777.150 | 11 | | | |

| **Group 1** | **Group 2** | **p-value** | **Significance** | **Comments** |
|---|---|---|---|---|
| Bare Glass LS | Bare Glass | 0.00714 | ** | The addition of PBS resulted in a significant decrease of yield in comparison to gentle hydration in low salt. |
| Bare Glass | HGT | 0.738 | NS | Use of compound in Group 2 was ineffective at significantly increasing yields of GUVs compared to gentle hydration in PBS. |
| Bare Glass | PVA | 4.282E-04 | *** | Use of compounds in Group 2 resulted in a significant increase in yields of GUVs compared to gentle hydration in PBS. |
| Bare Glass | LGT | 1.464E-03 | *** | |
| HGT | PVA | 1.662E-04 | *** | The difference of yield between HGT and PVA is significant. |
| HGT | LGT | 8.819E-14 | *** | The difference of yield between HGT and LGT is significant. |
| PVA | LGT | 4.041E-12 | *** | The difference of yield between PVA and LGT is significant. |

**Table S6:** ANOVA table and table of p-values from post hoc Tukey's HSD tests of the molar yields of GUVs composed of the MEL mixture grown at 22 °C obtained from gentle hydration on bare glass in 100 mM sucrose (Bare Glass LS), gentle hydration on bare glass in PBS (Bare Glass) and hydration on polymer films of PVA, LGT agarose, and HGT agarose in PBS. All solutions contain 100 mM of sucrose. * = $p < 0.05$, ** = $p < 0.01$, *** = $p < 0.001$, NS = not significant.

| Source | SS | df | MS | F | Probability>F (p-value) |
|---|---|---|---|---|---|
| Columns | 214.874 | 2 | 107.437 | 59.687 | 1.096E-04 |
| Error | 10.800 | 6 | 1.800 | | |
| Total | 225.674 | 8 | | | |

| **Group 1** | **Group 2** | **p-value** | **Significance** | **Comments** |
|---|---|---|---|---|
| DOPC | ERGIC | 5.46E-04 | *** | The difference of yield between DOPC and ERGIC is significant. |
| DOPC | MEL | 0.0868 | NS | The difference of yield between DOPC and MEL is not significant. |
| ERGIC | MEL | 1.08E-04 | *** | The difference of yield between ERGIC and MEL is significant. |

**Table S7:** ANOVA table and table of p-values from post hoc Tukey's HSD tests of the molar yields of GUVs composed of the MEL mixture, DOPC mixture and ERGIC mixture assembled at 22 °C on using LGT agarose as the assisting compound in PBS. All solutions contain 100 mM of sucrose. * = $p < 0.05$, ** = $p < 0.01$, *** = $p < 0.001$, NS = not significant.

| Source | SS | df | MS | F | Probability>F (p-value) |
|---|---|---|---|---|---|
| Columns | 111.428 | 2 | 55.714 | 37.895 | 3.95E-04 |
| Error | 8.821 | 6 | 1.470 | | |
| Total | 120.249 | 8 | | | |

| **Group 1** | **Group 2** | **p-value** | **Significance** | **Comments** |
|---|---|---|---|---|
| DOPC | ERGIC | 0.00133 | ** | The difference of yield between DOPC and ERGIC is significant. |
| DOPC | MEL | 0.358 | NS | The difference of yield between DOPC and MEL is not significant. |
| ERGIC | MEL | 4.43E-04 | *** | The difference of yield between ERGIC and MEL is significant. |

**Table S8:** ANOVA table and table of p-values from post hoc Tukey's HSD tests of the molar yields of GUVs composed of the MEL mixture, DOPC mixture and ERGIC mixture assembled at 22 °C on using PVA as the assisting compound in PBS. All solutions contain 100 mM of sucrose. * = $p < 0.05$, ** = $p < 0.01$, *** = $p < 0.001$, NS = not significant.

| Source | SS | df | MS | F | Probability>F (p-value) |
|---|---|---|---|---|---|
| Columns | 2275.586 | 4 | 568.897 | 196.320 | 1.87E-09 |
| Error | 28.978 | 10 | 2.898 | | |
| Total | 2304.564 | 14 | | | |

| **Group 1** | **Group 2** | **p-value** | **Significance** | **Comments** |
|---|---|---|---|---|
| Low Salt | 140 mM KCl + 5 mM $CaCl_2$ | 2.15E-08 | *** | Assembly using the buffer in Group 2 results in a significant decrease in yields of GUVs compared to assembly using the buffer in Group 1. |
| Low Salt | PBS + 5 mM $MgCl_2$ | 2.13E-08 | *** | |
| Low Salt | 600 mM NaCl | 1.77E-08 | *** | |
| Low Salt | PBS | 1.43E-07 | *** | |
| PBS | 600 mM NaCl | 8.14E-05 | *** | |
| PBS | PBS + 5 mM $MgCl_2$ | 0.00124 | ** | |
| PBS | 140 mM KCl + 5 mM $CaCl_2$ | 0.00137 | ** | |
| PBS + 5 mM $MgCl_2$ | 140 mM KCl + 5 mM $CaCl_2$ | 1.000 | NS | Assembly using the buffer in Group 1 or Group 2 results in no significant difference in the yields of GUVs. |
| PBS + 5 mM $MgCl_2$ | 600 mM NaCl | 0.228 | NS | The addition of 5 mM $MgCl_2$ has an equivalent effect on the yield of GUVs as using 600 mM NaCl. |
| 140 mM KCl + 5 mM $CaCl_2$ | 600 mM NaCl | 0.206 | NS | The addition of 5 mM $CaCl_2$ has an equivalent effect on the yield of GUVs as using 600 mM NaCl. |

**Table S9:** ANOVA table and table of p-values from post hoc Tukey's HSD tests of the molar yields of GUVs assembled obtained using LGT agarose as the assisting compound in buffers of different ionic compositions. All solutions contain 100 mM of sucrose. * = $p < 0.05$, ** = $p < 0.01$, *** = $p < 0.001$, NS = not significant.

| Salt | 100 mM sucrose | PBS+100 mM sucrose | PBS + 5 mM $MgCl_2$ +100 mM sucrose | 140 mM KCl + 5 mM $CaCl_2$ + 100 mM sucrose | 600 mM NaCl + 100 mM sucrose |
|---|---|---|---|---|---|
| NaCl | - | 137 mM | 137 mM | - | 600 mM |
| KCl | - | 2.7 mM | 2.7 mM | 140 mM | - |
| $Na_2HPO_4$ | - | 8 mM | 8 mM | - | - |
| $KH_2PO_4$ | - | 2 mM | 2 mM | - | - |
| $MgCl_2$ | - | - | 5 mM | - | - |
| $CaCl_2$ | - | - | - | 5 mM | - |

**Table S10.** Table of the concentration of different salts in the hydration solutions used to gather the data in Figure 9.

| Ionic species | 100 mM sucrose | PBS+100 mM sucrose | PBS + 5 mM $MgCl_2$+100 mM sucrose | 140 mM KCl + 5 mM $CaCl_2$+100 mM sucrose | 600 mM NaCl+100 mM sucrose |
|---|---|---|---|---|---|
| $Na^+$ | - | 153 mM | 153 mM | - | 600 mM |
| $K^+$ | - | 4.7 mM | 4.7 mM | 140 mM | - |
| $H^+$ | 0.0032 mM | $4\times10^{-5}$ mM | $4\times10^{-5}$ mM | 0.0032 mM | 0.0032 mM |
| $HCO_3^-$ | 0.0032 mM | $4\times10^{-5}$ mM | $4\times10^{-5}$ mM | 0.0032 mM | 0.0032 mM |
| $Cl^-$ | - | 139.7 mM | 149.7 mM | 150 mM | - |
| $HPO_4^{2-}$ | - | 8 mM | 8 mM | - | 600 mM |
| $H_2PO_4^-$ | - | 2 mM | 2 mM | - | - |
| $Mg^{2+}$ | - | - | 5 mM | - | - |
| $Ca^{2+}$ | - | - | - | 5 mM | - |
| **Debye Length** | 170 nm | 0.75 nm | 0.73 nm | 0.77 nm | 0.39 nm |

**Table S11.** Table of the concentration of ions in the hydration solutions. We assume complete dissociation of all ions. The last row shows the calculated Debye screening length.
